# 4D multimodal wound healing atlas reveals organ-level controls of repair phase transitions

**DOI:** 10.64898/2026.01.15.699736

**Authors:** Jonathan Chin Cheong, Simon Van Deursen, Dreyton Amador, Shannon Hiner, Yvon Woappi

## Abstract

Deep skin wounds demand tightly coordinated communication across diverse tissue systems, yet knowledge of the molecular logic governing organ-scale injury response remains incomplete. Existing wound atlases profile fragments of this process, capturing limited tissue groups and healing phases, obscuring how whole organs synchronize repair. Here, we present the Organ-Scale Wound Healing Atlas (OWHA), a 4D multimodal omnibus that integrates snRNA-seq, scRNA-seq, CITE-seq and high-definition spatial transcriptomics to reconstruct the complete spatial and temporal choreography of mammalian wound healing at single cell resolution. OWHA profiles over 725,000 murine single-cell and spatial transcriptomes encompassing the entire wound healing process from early to late healing phases across the vast skin microanatomical tissue niches. This omnibus overcomes long-standing technical limitations, enabling robust resolution of adipocytes, Schwann cells, fragile epithelial intermediates, and over 100 precisely annotated cell states, including populations missed in prior wound databases. This revealed that wound repair proceeds through sharp transcriptional and cellular inflection points driven by Central Orchestrator populations that coordinate healing via synchronized transcriptional activation and direct cross-tissue signaling. Key among these is a *Sox6*^+^ *Tspear*^+^ *Il20ra*^+^ keratinocyte subpopulation (Basal IV), detectable only through snRNA-seq but entirely missed by conventional wound atlasing. After injury, Basal IV cells deviate from canonical differentiation programs and adopt a neurovasculogenic signaling state during the proliferation phase, forming a transient spatially privileged regulatory hub at the wound edge. This epithelial-anchored niche spatially aligns Basal IV keratinocytes with proliferative endothelial cells, Pericytes, and Repair Schwann Cells, synchronizing re-epithelialization, angiogenesis, and neurite guidance. Mechanistically, this is orchestrated by a conserved Sema3C–Nrp1/Nrp2 axis that coordinates epithelial–vascular–neuronal crosstalk at the wound site. Cross-species integration confirms that the Basal IV/SEMA3C axis is conserved in human skin, yet undetected by conventional scRNA-seq human atlases due to dissociation-induced artifacts – underscoring the critical need for multimodal atlasing to accurately capture organ-scale physiology. Notably, the Basal IV/SEMA3C circuitry is selectively disrupted in human diabetic wounds, but topical Sema3C treatments restores peri-wound angiogenic sprouting and accelerates re-epithelialization of diabetic ulcers *in vivo*. OWHA establishes the first 4D, organ-scale molecular blueprint of mammalian wound healing, creating a foundational platform for decoding systems-level principles of repair and regeneration for tissue wounds.

## INTRODUCTION

Mammalian organs are highly complex biological structures, organized into multiscale hierarchies of functional units maintained by tightly regulated interactions between distinct cells and tissues. Restoration of organ function after injury therefore requires precise spatial and functional coordination to ensure that correct cell types reintegrate into their appropriate tissue environment^1–4^. Yet, decoding this process in mammals remains challenging due to physiological complexity, which has often confined wound studies to a limited set of tissue compartments and phases^5–10^. As the largest organ in mammals, the skin is an intricate composite of specialized stem cell niches and mini-organs that have evolved for protection, sensory perception, and repair^11–13^. The integument encapsulates cellular, structural, and functional analogs of multiple organ systems, enabling the study of regeneration, immune dynamics^14,15^, neurosensory feedback^13^, vascular remodeling^16^, and musculoskeletal interactions^17,18^ within a single accessible tissue^17,19,20^. Consequently, the skin wound healing process is an ideal model to investigate systems-level logic of mammalian organ repair.

Despite decades of extensive study of the wound healing process, the coordinated interactions between structural, adnexal, neuronal, vascular, and immune compartments throughout injury response remain incompletely understood. More recently, rigorous molecular wound atlases have provided valuable insights into the processes involved in injury response^5–10,21^, but they have often lacked critical late remodeling timepoints, exhibited strong sex imbalance, or consistently underrepresented key cell populations, including fascia, adipocytes, Schwann cells, melanocytes, and red blood cells due to dissociation and microfluidic-based constraints^22,23^. Moreover, past studies have largely relied on unimodal profiling strategies and lacked high resolution spatial information, restricting the analysis of multi-compartment coordination to general cell groups and obscuring the fine-grained spatial interactions essential for organ-level response. Together, these limitations have constrained the comprehensive understanding of how diverse skin tissue groups synchronize repair.

To overcome this, we constructed the Organ-Scale Wound Healing Atlas (OWHA), a comprehensive, time-resolved, multimodal omnibus of full-thickness murine skin wound repair. OWHA integrates rigorously benchmarked publicly available single cell RNA sequencing (scRNA-seq) data with newly acquired, sex-balanced datasets generated using three complementary modalities: cellular indexing of transcriptomes and epitopes by sequencing (CITE-seq), single nuclei RNA sequencing (snRNA-seq), and Visium HD. This multimodal approach overcomes long-standing technical barriers, enabling the recovery of fragile, rare, and extra-large cell types, along with their precise positions in tissue space. Leveraging OWHA, we uncovered that wound healing is organized around sharp transcriptional and cellular inflection points that demarcate distinct phases of repair. These transitions are coordinated by a hierarchical regulatory architecture featuring Central Orchestrator (CO) populations that integrate transcriptional activation with directed cross-tissue signaling.

Notably, we identify a wound-adaptive basal keratinocyte subpopulation, termed Basal IV, characterized by elevated expression of *Sox6, Tspear,* and *Il20ra.* Basal IV keratinocytes represent a key CO population frequently underrepresented or missing in conventional scRNA-seq datasets due to dissociation-induced artifacts. Following injury, Basal IV keratinocytes deviate from canonical differentiation trajectories during the proliferative phase and adopt a transient neurovasculogenic signaling program at the wound edge, supported by epithelial-derived SEMA3C. Cross-species analysis with human skin wounds^24^ confirms that *Sox6*^+^ epidermal lineages are conserved in both mouse and human skin. Importantly, the Basal IV state – undetectable in conventional human scRNA-seq atlases – becomes readily identifiable in human spatial transcriptomic data, underscoring the necessity of multimodal approaches for accurately capturing organ-scale transcriptomics. Finally, we demonstrate that the SEMA3C axis is impaired in human diabetic ulcers^25^, and that topical recombinant SEMA3C treatment rescues defective re-epithelialization and peri-wound angiogenesis *in vivo*. Together, OWHA provides a comprehensive, organ-scale exploration of the mammalian wound healing process and reveals phase-resolved, higher-order principles governing integument physiology and reparative fate decisions.

## RESULTS

### Multimodal single-cell transcriptomic atlasing comprehensively captures the murine wound healing process at organ scale

To construct OWHA, we first conducted a comprehensive survey of all publicly available single-cell sequencing datasets that sampled 4–6 mm full-thickness mouse skin wounds across several healing timepoints (**Table S1; See Methods**). Three scRNA-seq datasets met inclusion criteria that provided comprehensive representation of full thickness wound healing in mice (**Figure S1A**). However, integrative analysis of these datasets revealed several major limitations: (i) absence of late remodeling timepoints and phases, such as day 15 and 30 post-wounding (D15PW, D30PW) (ii) consistent underrepresentation of cell types including Schwann cells, adipocytes, and red blood cells and (iii) a pronounced sex imbalance, with most datasets derived exclusively from female mice (**Figures S1B–S1D, S2G–S2I**). Together, these gaps underscored the practical challenges in constructing an organ-scale atlas^26,27^ and the critical need for more comprehensive wound datasets, as achieving this requires complete tissue sampling across multiple healing phases and tissue groups, all while ensuring sample sex balance^28,29^. Given that our primary objective was to investigate organ-level responses, rather than mere isolated tissue compartments, we set out to address these limitations directly. We therefore employed an integrative strategy that combined rigorously benchmarked public scRNA-seq datasets with newly generated sex-balanced multimodal sequencing datasets from our group (**Figures S2C-S2I and S3**). All public data were realigned to GRCm39 (mm39)^30^ (**Figures S2A and S2B**) to maximize seamless integration, and newly generated wound samples (batches) were collected with comparable tissue harvesting protocols (**Table S2; See Methods**)^29^. Next, to expand cellular resolution at timepoints missing from prior studies, we performed CITE-seq^31^ on unwounded (UW) skin and late-stage wounds at D15PW and D30PW, yielding 37,894 cellular transcriptomes with simultaneous surface epitope expression (**Figures S2C-S2E**).

Nonetheless, we found that adipocytes remained missing from both scRNA-seq and CITE-seq, hereafter referred to as whole-cell sequencing, datasets including UW samples (**Figures S4A-S4F**). This is consistent with prior reports demonstrating the incompatibility of adipocytes’ large size, buoyancy, and lipophilic composition with lipid-based droplet encapsulations used in conventional single-cell platforms^32^. To overcome this, we performed paired snRNA-seq of full-thickness skin wounds across matching timepoints (i.e. UW, D4PW, D7PW, D15PW, and D30PW) captured by whole-cell sequencing. This yielded an additional 54,749 single-nuclei transcriptomes captured across the entire wound healing time course (**Figures S2C and S4C; Table S2**). Notably, our snRNA-seq data enabled the identification of multiple adipocyte subpopulations and rare cell types otherwise challenging to recover with whole-cell sequencing workflows^32,33^ (**Figures S4G-S4H**). Altogether, we profiled 193,524 single cell transcriptomes obtained from 28 sequencing batches generated by three complementary single-cell transcriptomic modalities – snRNA-seq, CITE-seq, and scRNA-seq – all within an integrated framework we termed OWHA. This unified 4D wound healing omnibus captures 11 major cell classes (metaclusters) encompassing all major integumentary tissue sub-compartments and their respective activities throughout wound repair (UW, D1PW, D2PW, D4PW, D7PW, D15PW, D30PW) (**Figures 1A-1G and S4**). Stringent integration benchmarking and rigorous quality control (QC) analyses revealed that OWHA was sex-balanced and demonstrated consistent cell-cluster distributions and mean cell recovery across all modalities, wound conditions, sexes, and chemistries (**Figures S2F-S2G, S3, S5 and S6**), nearly doubling the molecular and cellular resolution previously available in comparable wound-healing datasets (**Figure S2E**).

**Figure 1.**
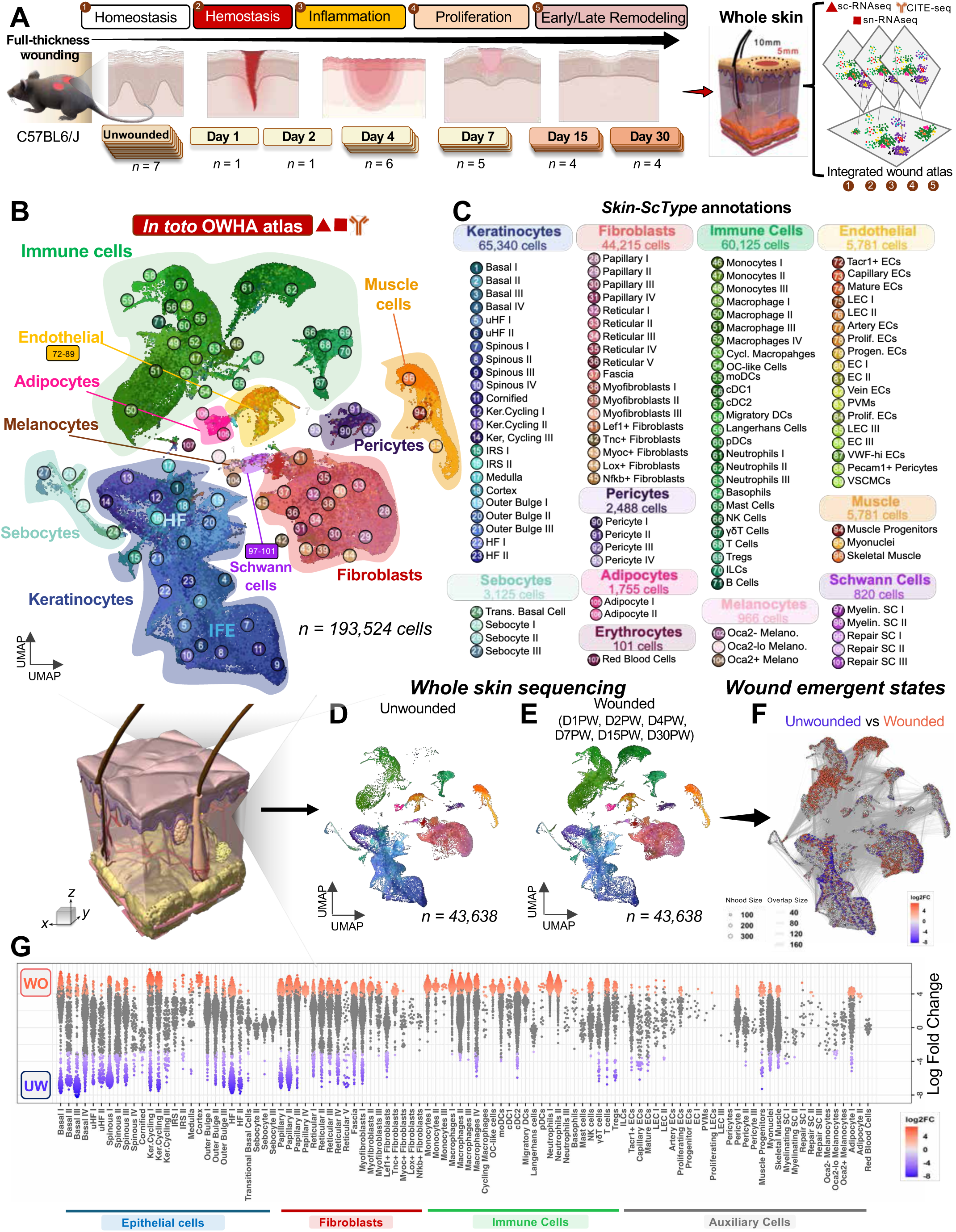
Multimodal transcriptomic skin atlas captures all major tissue groups active throughout wound healing. (**A**) Experimental schematic detailing the wound samples collection timeline and subsequent sequencing modalities used to generate an integrated single-cell atlas. (**B**) UMAP embedding of the integrated organ-scale wound healing atlas (OWHA) visualizing 193,524 single cells collected from full-thickness skin wounds. (**C**) Skin-ScType annotations for major skin cell types and corresponding subtypes represented in OWHA. A total of 11 major cell groups and 107 cell subtypes are captured at each phase of healing. (**D**) Integrated UMAP visualization of cells from unwounded timepoint, down sampled to 43,638 cells. (**E**) Integrated UMAP visualization of wounded timepoints at D1PW, D2PW, D4PW, D7PW, D15PW, and D30PW down sampled to 43,638 cells. (**F**) Milo differential abundance analysis highlighting wound emergent states (red) and cell states enriched during the unwounded (blue) timepoint. (**G**) Beeswarm plot of Milo differential abundance across all OWHA subclusters found in unwounded (blue) or wounded (red) timepoints. FDR = 0.10.

### *Skin-ScType* enhances the annotation accuracy of wound-associated cell clusters

Mammalian wounds contain a mixture of skin-resident cells and transient populations recruited from circulation and distal tissues, creating a cellular ecosystem of markedly high complexity^5,7^. This heterogeneity, combined with the diverse physiological states present across spatial compartments of the integument, makes transcriptional annotation of wound-derived cells particularly challenging^34,35^. We thus sought to develop a robust methodology that could accurately label all cellular populations captured by OWHA. First, we evaluated three well-established single-cell annotation methods, including *ScType*^36^, *GPTcelltype*^37^, and *SingleR*^38^ which rely on marker-gene expression and are routinely used for skin and other organ groups (**Figures S7A-S7D**). While these methods reliably labeled hematopoietic cell types and certain epithelial-derived populations, we observed substantial misannotations of cell types critical to the skin wound healing response, including fibroblasts, keratinocytes, and neurovascular cell types (**Figures S7B-S7D**). In particular, these annotation databases largely relied on human marker genes, which proved suboptimal for murine skin cell annotations (**Figures S7F-S7K**). To address this gap, we constructed a mouse-optimized *ScType* database (*Skin-ScType*), specifically designed to capture the complex murine skin cell taxonomy and its compositional changes throughout healing (**Table S3**). Notably, *Skin-ScType* accurately identified all major canonical cell groups involved in skin wound response^2,20^, including distinct keratinocyte and immune subpopulations (**Figures S7E and S7K**). To validate this annotation, we leveraged the multimodal profiling within OWHA, verifying cells’ RNA based annotations with CITE-seq protein marker expression which confirmed key metacluster identities including keratinocytes (CD49f+), leukocytes (CD45+), fibroblasts (Ly6A/E+), endothelial cells (CD31+), and Schwann cells (CD9+) (**Figures S7L-S7N**). This combined automated transcriptomic annotation with single cell marker protein validations provided a reproducible and precise cell classification system that corrected cluster misannotations (**Figures S7F-S7K**). Using this approach, we classified all major cell types captured by OWHA (**Figures 1B and 1C**) and established a comprehensive taxonomy of the mammalian wound across all of its healing stages (**Figures 2A and 2B**). In total, the 11 *Skin ScType*-annotated metaclusters identified by OWHA included fibroblasts (*Pdgfra*, *Col1a2*), immune cells (*Ptprc*), skeletal muscle (*Mylpf*, *Des*), pericytes (*Rgs5*), endothelial cells (*Pecam1*), interfollicular epidermis (IFE) keratinocytes (*Krt14*, *Itga6*), sebocytes (*Pparg*), and additional hair follicle (HF)-associated populations (*Krt14*, *Sox9*, *Krt15*) (**Figure 1C**). Importantly, OWHA also captured 107 skin subpopulations including previously missed groups, such as Schwann cells (*Mpz*), adipocytes (*Adipoq*), melanocytes (*Mlana*), and red blood cells (*Hba-a1*, *Hbb-bt*) (**Figures 1B, 1C, 2B and S8; Table 1**).

**Figure 2.**
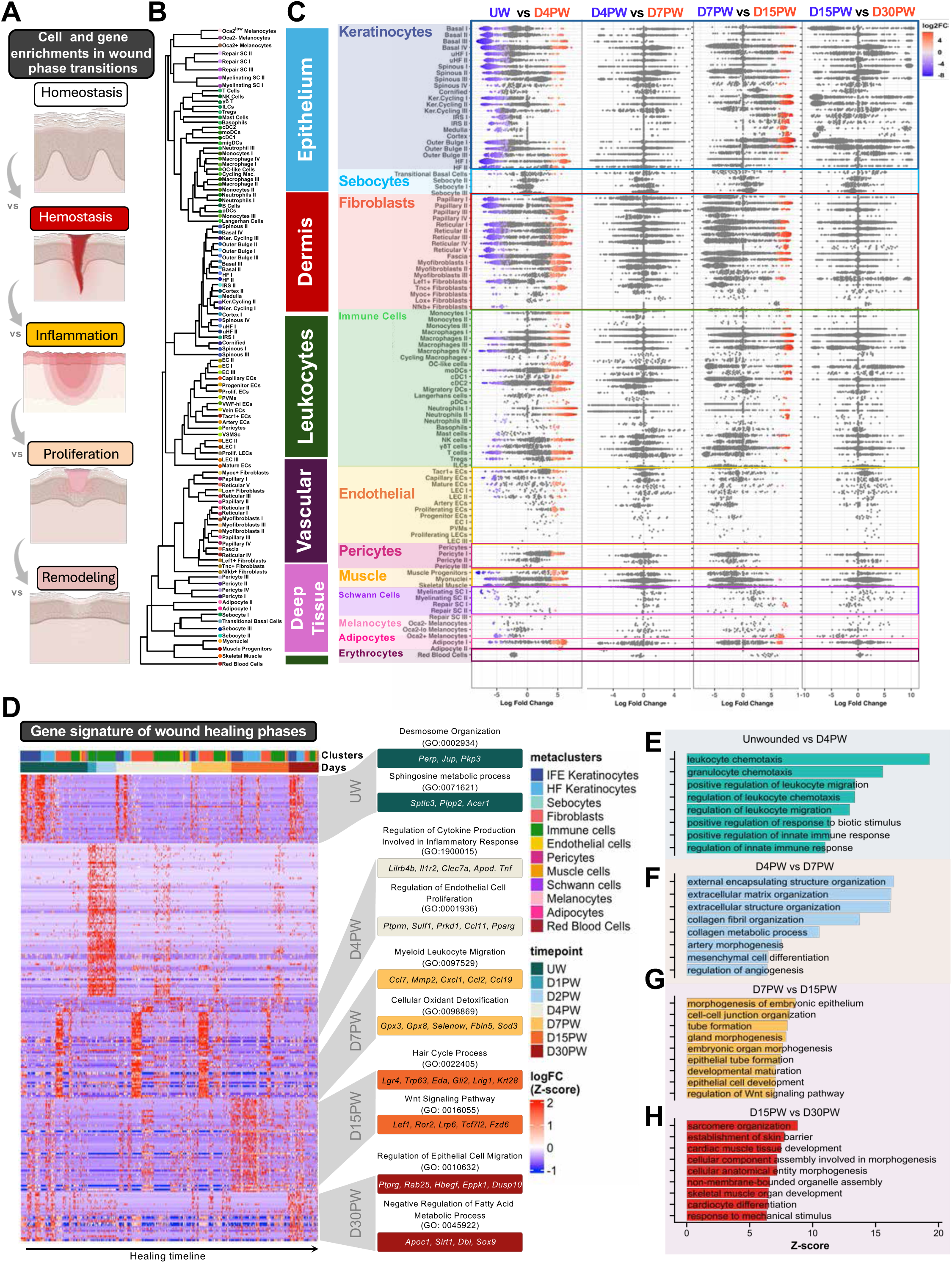
All major skin compartments actively contribute to wound phase transitions in full-thickness wounds. (**A**) Schematic summarizing differential cell and gene abundance analysis across wound phase transitions captured by OWHA. (**B**) Dendrogram illustration of all 107 OWHA-identified subclusters. (**C**) Phase-specific Beeswarm plots of Milo differential abundance analysis reveals dynamic enrichment and depletion of cell types across successive wound healing phase transitions (UW vs. D4PW, D4PW vs. D7PW, D7PW vs. D15PW, and D15PW vs. D30PW; FDR = 0.10). This highlights specific cell subclasses that emerge at distinct transition points, uncovering both transient and sustained activation of unique cell states during healing. (**D**) Heatmap displaying top 200 differentially expressed genes marking each wound timepoint across skin Metaclusters (left), and Gene Ontology (GO) pathways and genes associated with each timepoint (right). (**E–H**) Z-score bar plots of uniquely upregulated Gene Ontology programs active at each sequential phase transition timepoint (UW→D4PW, D4PW→D7PW, D7PW→D15PW, D15PW→D30PW; p < 0.05).

### Wound repair is driven by emergent biological shifts rather than linear temporal progressions

To define how each skin tissue compartment contributes to wound repair, we quantified differential cell-state abundances across consecutive healing timepoints using Milo^39^. Early after injury (UW to D4PW), epithelial and melanocyte populations were globally depleted two-fold compared to UW (**Figures 2C**, **S9A, S9D, S9E, and S9M**), consistent with the expected physical loss of the epidermis and HF structures immediately after acute injury^20,40^. In contrast, immune and fibroblast compartments expanded sharply, and their activation states were maintained through the proliferation-to-maturation transition (D7PW-D15PW) (**Figures 2C, S9A-S9C, and S9F**). Strikingly, despite large scale tissue loss, Milo revealed extensive wound-emergent cellular states across nearly every skin compartment, including in metaclusters severely depleted after injury (**Figures 2C and S9**). Within the epithelium, Basal IV (*Sox6+, Tspear+, Il20ra+)* and HF I *(Ch25h+, Guca2a+, Serpinb10+)* displayed the strongest enrichment during the UW-to-D4PW transition, marking an early shift toward stress-responsive re-epithelization and HF activation^41–43^ (**Figure 2C and S10A-S10D)**. At the D7PW-to-D15PW transition, this activation was most dominantly seen in Outer Bulge I (*Lgr5+, Kcnb2+*) cells, suggesting that the epithelium undergoes a biphasic activation with early basal/HF activation at D4PW and a second later activation of the HF/outer bulge cells after the D15PW remodeling transition, supporting previous reports^5,41^ **(Figure 2C).**

In the dermis, Papillary I (*Igfbp2*+, *Ly6c1*+, *Pla2g5*+) and Fascia (*Opcml+, Tmeff2+*) fibroblasts showed the highest early (D4PW) enrichments. However, Papillary IV (*Cthrc1+*, *Pcsk5*+), Myofibroblasts II (*Col11a1*+, *Mme*+, *Coch*+), and *Tnc*⁺ fibroblasts were exclusively induced after injury, indicating emergence of fibroblast subsets not typically found during homeostasis^7,44^. Interestingly, both *NfKb*+ and Reticular V fibroblasts (*Procr*+, *Gfpt2*+) exhibited significant depletion following injury, suggesting restricted regenerative plasticity in these cell types early in the repair process. By the D7PW-to-D15PW transition, Reticular III (*Sfrp4+, Fstl1+*) displayed the highest enrichment among fibroblasts, indicating a strong deep dermis wound maturation role (**Figure 2C and S10E-S10H**). In total, the dermis displayed the greatest expansion of injury-induced lineages across all skin tissue groups. Among leukocytes, cDC2 (*Mgl2*+, *Cd209a*+, *H2-Eb1*+) and Neutrophil (*Retnlg*+, *Cxcr2*+) subsets showed the most pronounced early activations, consistent with the inflammatory influx characteristic of the initial wound response^45^, while the late healing phases were mostly dominated by macrophage activations. Among lymphatic and vascular cells, *Tacr1*+ ECs, Proliferating (*Mki67*+) Endothelial cells (PECs), and Pericyte I (*Rgs5*+, *Adra2a*+) were most strongly and nearly exclusively enriched in the UW-to-D4PW transition, matching the activation patterns seen in early keratinocyte transitions^46^ (**Figures S10 and S11)**. In the deep tissue, Adipocyte I (*Plin1*+, *Adipoq*+) and Muscle Progenitors (*Heyl*+, *Myf5*+) displayed the strongest early activation state, while Schwann cells were largely depleted early except for Repair SC II (*Cdh19+, Foxd3+*) (**Figures S12 and S13**). While at late timepoints, Repair SC I (*Cdh19+, Stard13+, Kcna6+*) displayed the highest enrichment (**Figures 2C and S14A-S14D**). Interestingly, several cell populations showed enrichment states during both UW and wounded (WO) conditions (e.g. UW vs D4PW) (**Figures 1G and 2C**), suggesting that many skin populations undergo condition-specific shifts in their transcriptional programs in response to injury relative to their unwounded states. Nonetheless, minimal changes in transcriptional states or cellular abundance were observed between the early to late maturation phase (D15PW-to-D30PW), revealing that this interval represents a terminal or stabilized stage of wound healing with respect to population-level state changes (**Figures 2C, S9-S13**). Altogether, these results reveal that wound repair proceeds through coordinated emergence of several temporal cellular fates across a wide array of tissue groups, culminating in a near-complete restoration of the homeostatic transcriptional landscape during late maturation.

Strikingly, sequential differential abundance analysis across all timepoints revealed that the most robust induction of wound-emergent cell states occurred at major physiological inflection points in the healing process (**Figure 2C**). These included the transition from homeostasis to inflammation/proliferation (UW-D4PW) and from proliferation to maturation (D7PW-D15PW). Conversely, no significant changes in wound-activated cell states were detected within similar physiological wound healing stages, such as early to late proliferation (D4PW-D7PW) and early to late maturation (D15PW-D30PW), even though this stage spans a longer temporal interval (**Figure 2C**). Together, these findings indicate that phases transitions in wound-healing occur at distinct inflection points. Between these transitions however, wounded skin compartments remain remarkably transcriptionally and physiologically stable.

To define the molecular programs underlying each phase of wound repair, we first examined Gene Ontology (GO) signatures uniquely activated at individual healing timepoints (**Figures 2D and S14E**). This revealed that early responses (D4PW) were expectedly dominated by inflammatory cytokine production (*Tnf, Clec7a, Il1r2*) and endothelial cell proliferation (*Ccl11, Pparg, Ptprm*), while the D7PW proliferation phase was defined by myeloid leukocyte migration (*Ccl2*, *Cxcl1*, *Ccl19*) and cellular antioxidant activity (*Gpx3*, *Gpx8*, *Selenow*). At D15PW, transcriptional profiles shifted toward epithelial and glandular morphogenesis, marked by induction of hair cycle (*Gli2, Lrig1, Lgr4*) and Wnt signaling (*Fzd6, Ror2, Lef1*) pathways, indicative of pilosebaceous and epithelial regeneration at the wound periphery. Subsequently, D30PW, corresponding to tissue late maturation, was dominated by strong induction of epithelial migration (*Ptprg, Hbegf, Eppk1*) and fatty acid metabolism (*Apoc1, Sirt1, Sox9*) programs, consistent with epidermal barrier and adnexal structure restoration (**Figures 2D and S14E**). To investigate how these programs are synchronized for phase transition, we performed sequential differential gene expression analysis between consecutive timepoints (**Figures 2A, 2E-2H and S14F**). This revealed that leukocyte chemotaxis dominated the UW-to-D4PW transition (**Figure 2E**), reflecting the immune influx after acute injury. Notably, while the D4PW-to-D7PW phase transition was primarily defined by extracellular matrix (ECM) reorganization, a striking induction of vascular morphogenesis and angiogenesis regulators emerged (**Figure 2F**), processes often disrupted in ischemic wounds^47,48^. The D7PW-to-D15PW transition, another major physiological inflection point, was defined by organotypic pathways related to epithelial morphogenesis, embryonic organ patterning, and gland development, indicating a reengagement of developmental pathways to rebuild specialized skin structures (**Figure 2G**)^20,49^. Finally, the D15PW-to-D30PW transition was characterized by signatures related to muscle development and mechanical stimulus, epidermal barrier establishment, and extracellular component assembly, reflecting structural maturation and mechanical reinforcement of the newly formed tissue (**Figure 2H**). Together, these findings indicate that full-thickness wound healing is governed by interconnected and time-ordered molecular programs that transition from immune mobilization to structural regeneration and ultimately barrier reinforcement by D30PW. This temporal choreography likely ensures that early inflammatory cues prime the tissue microenvironment for vascularization and matrix deposition while later healing programs facilitate epidermal integrity, mechanical resilience, and immune clearance. Importantly, the emergence of developmental and morphogenetic signatures during mid-to-late healing phases indicates that skin repair recapitulates many aspects of embryonic tissue patterning^50,51^. Together, these analyses demonstrate that wound healing phase transitions are driven by coordinated, time-specific transcriptional programs, whereby early immune mobilization creates an inflammatory milieu which progresses into a vascularization state, ultimately giving rise to organ repatterning and tissue restoration.

### Wound healing phase transition integrates distinct signaling hubs across vast anatomical and temporal scales

To investigate how healing-associated signaling is orchestrated across the entire wound healing timescale, we subdivided signaling analysis by specific tissue groups. While extensive characterization of cell signaling at the wound site has been conducted in previous years^3,4,28,29,52,53^, most have focused on isolated cell types or discrete stages of repair, limiting insight into how intercellular communication is coordinated across the full wound healing continuum. We thus examined whether cell–cell crosstalk remained confined to local tissue compartments or extended across distal skin layers, including epidermis, dermis, and deeper fascial tissue (**Figure 3A**). CellChat^54^ interactome analysis revealed that each major skin compartment contained distinct subpopulations with elevated sender and receiver activity that dominated specific timepoints in the repair process (**Figures 3B-3G and S15**). In the UW epidermis, *Krt10*-expressing Spinous I populations were the dominant signaling nodes. However, following injury, Basal IV keratinocytes – one of the strongest wound-emergent subpopulations – were the dominant signalers (**Figure 3B)**. This signaling activity was most pronounced at the early and late proliferation phases (D4PW-D7PW), coinciding with active re-epithelization in the wound site. This suggests Basal IV keratinocytes act as a key coordinator of early epidermal repair (**Figures 3B, S15B and S16A-S16H**). By early remodeling (D15PW), HF-associated Keratinocyte cycling I cells were the dominant signalers, suggestive of a shift to early wound remodeling activity by adjacent HF cells^41,43,55^. By late remodeling (D30PW), Spinous I keratinocytes then returned as top signalers of the epidermis, revealing that in addition to cell number recovery, wound healing also restores tissue-specific signaling hierarchies (**Figure 3B**). We next examined signaling dynamics between all sebaceous lineages associated with the pilosebaceous unit. Interestingly, while transitional basal cells had the largest enrichment (FC >1) of wound-emergent sebocytes, they were the least involved in sebocyte-mediated signaling. Conversely, the less significantly enriched Sebocyte III subpopulation (*Ces4a*+, *Cidea*+) was the dominant signaler across all healing timepoints, with the exception of D30PW where Sebocyte I (*Akr1c19***+,** *Acoxl*+), a population previously shown to be involved in lipid-mediated repair^56^, emerged as the dominant signaler (**Figures 3C, S15C and S16I-S16P**).

**Figure 3.**
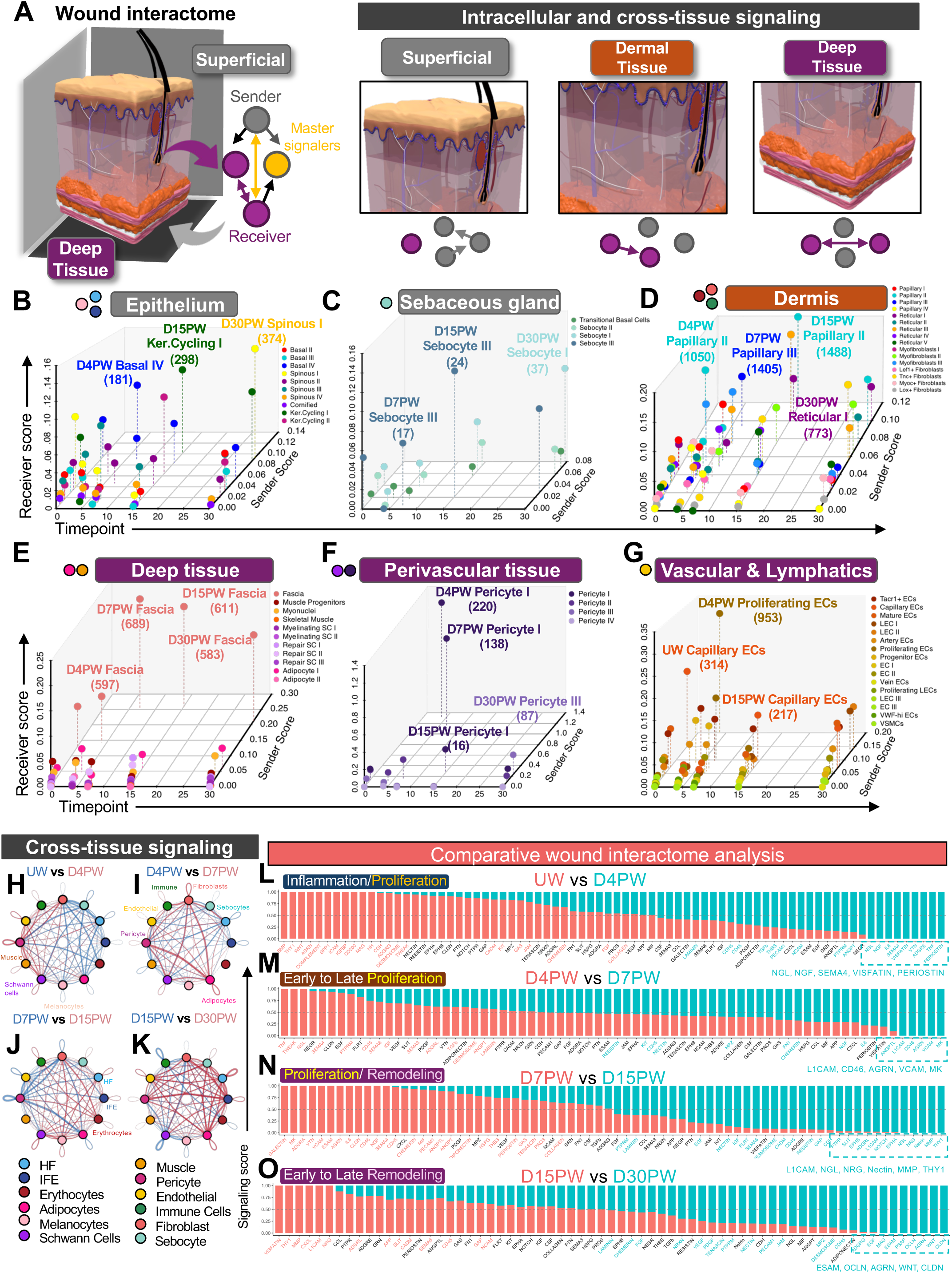
Wound-emergent populations coordinate tissue repair signaling across multiple tissue groups. (**A**) Outline of the experimental workflow for identifying key signaling domains mediating intracellular, cross-tissue, and inter-compartment communication across all skin strata. This encompassed superficial tissues (grey), dermal tissue (orange), and deep tissue structures (purple). (**B-G**) Three-dimensional manifold projections of intracellular signaling landscapes, quantifying sender and receiver strengths across major skin compartments during wound healing. Each plot highlights the predominant signaling populations within each major skin cluster: (**B**) Epithelium, (**C**) sebocytes, (**D**) dermis, (**E**) fascia and deep tissue, (**F**) perivascular tissue, and (**G**) vascular and lymphatic tissues. Points are color-coded by cluster identity, and the number of inferred outgoing interactions for each top signaling population is shown in parentheses, revealing phase-specific reorganization of intercellular communication networks in key signaling hubs throughout healing. **(H-K)** Differential signaling network centrality depicting cross-tissue changes in cell–cell communication roles across sequential wound healing transition phases. Comparisons include (**H**) unwounded (UW) vs. day 4 post-wound (D4PW), (**I**) D4PW vs. D7PW, (**J**) D7PW vs. D15PW, and (**K**) D15PW vs. D30PW. Edge width reflects the strength of predicted ligand–receptor interactions, revealing dynamic rewiring of cellular signaling networks as healing progresses. Red and blue chords indicate increased and decreased signaling, respectively, between groups at the second timepoint compared to the first. (**L–O**) Ranked global differential changes in signaling pathways uniquely enriched at key wound healing transition phases. Each bar represents a signaling pathway ranked by overall communication probability, highlighting phase-specific networks activated during distinct healing stages. Colored pathway names indicate significant enrichment for their respective timepoints. Wilcoxon test was used to determine whether there is significant difference between two datasets (p < 0.05).

Fibroblasts represented the most signaling-diverse cell group within the wound environment, exhibiting the greatest number of signaling interactions and the greatest heterogeneity of dominant signalers over time **(Figures 3D, S15A and S15D**). Reticular IV (*Apod+, Cyp2f2+*) and Papillary I group led signaling activity before injury, but Papillary II cells (*Igfbp2*+, *Eln*+, *Lox*+) dominated early (D4PW) injury response. By D7PW, dermal signaling again changed, now to Papillary III cells (*Cthrc1*+ *Lrrc15*+), before returning to Papillary II cells at D15PW. By D30PW, Reticular I (*Tmeff2*+, *Opcml*+) and Papillary II cells were the dominant signalers (**Figures 3D, S15D and S17A-S17H**). This sequential relay of fibroblast signaler activity highlights dynamic remodeling of stromal communication programs across healing phases. We next examined signaling within deep integument tissue structures, including adipocytes, Schwann cells, muscle, and fascia. Strikingly, while muscle and adipocytes were among the leading signalers in UW deep tissues, fascial cells emerged as dominant and persistent signaling hubs among all deep tissue categories across all timepoints after injury (**Figures 3E, S15G and S17I-S17P**). This supports recent studies implicating fascia as critical for large-scale wound reconstruction^17,57–59^ and reveal a previously under-characterized communication node in deep wound niches. Next, we investigated signaling within the vasculature. This revealed that Pericyte I and PECs exhibit high signaling at D4PW–D7PW before returning toward baseline by D30PW (**Figures 3F, 3G, S15E, S15F, and S18**). Together, these findings reveal a relay of transient but coordinated signaling interactions between designated cell groups within distinct tissue compartments throughout skin repair.

To next determine how signaling facilitated healing phase transitions, we performed differential CellChat analysis over time across all metaclusters detected within the wound environment. Signaling network centrality analysis revealed broad and balanced cross-lineage communication in the skin even before injury (**Figures 3H-3K**). However, these activities were markedly expanded as early as D4PW (**Figure 3H**). Moreover, the UW-to-D4PW transition was characterized by loss of interactions among major epidermal and subcutaneous metaclusters (**Figure 3H**), consistent with the physical loss of tissue after full-thickness injury and mirroring patterns observed in the differential abundance analyses (**Figure 2C**). Simultaneously, we observed induction of broad immune-derived signaling received by virtually all metaclusters, suggestive of a global inflammatory milieu generated and received by all wounded tissues after acute injury. During the D4PW-to-D7PW transition, the signaling infrastructure became more finely tuned, with strong and selective adipocyte-centric interactions that involved reciprocal signaling with dermal fibroblasts and pericytes (**Figure 3I)**. Additionally, distinct signaling pathways emerged between Schwann cells, fibroblasts, and immune cells, indicating targeted immune-stromal-neuronal interactions. The D7PW-to-D15PW transition displayed reduced interactions between adipocytes, endothelial cells, and pericytes while signaling increased between hair follicle-associated groups, including melanocytes, fibroblasts, and HF keratinocytes (**Figure 3J**). By the D15PW-to-D30PW transition, we observed a return to UW signaling patterns with increased interactions between epidermal lineages of the HF and IFE and virtually all other cells, reflecting a return to the broad multi-tissue crosstalk seen in uninjured skin (**Figure 3K**). This analysis revealed that while defined cell groups have dominant signaling roles throughout repair, global communication networks return to UW baseline levels by D30PW, recapitulating restoration trends seen at the population level (**Figures 2C, S9 and S14C**). Together, these findings indicate that wound healing phase transitions are driven by dynamic and selective signaling interactions that engage deep tissue compartments, including the adipose layer, as well as stromal and vascular populations, underscoring the necessity of holistic wound profiling to uncover these inter-tissue responses.

To define the molecular programs associated with the vast cross-tissue communication observed throughout the phase transitions, we performed differential interactome pathway analysis across all metaclusters at successive timepoints (**Figures 3L-3O**). This revealed an enrichment in NGL, NGF, and Sema4 signaling at the UW-to-D4PW transition, indicating activation of neurotrophic and axon-guidance pathways early in the healing processes **(Figure 3L).** Conversely, D4PW-to-D7PW interactomes were mediated by L1CAM, CD46, AGRN, VCAM, and MK signaling, suggesting angiogenesis, adhesion, and ECM remodeling active early in repair (**Figure 3M**). At the D7PW-to-D15PW transition, dominant signaling programs included L1CAM, NGL, NRG, Nectin, and THY1, suggesting progressive engagement of neurogenic and cell-adhesion pathways that may support vascular stabilization and tissue maturation at this transition phase (**Figure 3N**). By the D15PW-to-D30PW transition, ESAM, OCLN, AGRN, CLDN, and WNT pathways returned as dominant signaling programs (**Figure 3O**), reflecting the WNT and BMP signaling expressed in homeostatic skin^49,60^ **(Figure 3L**). This indicates that wound phase transitions rely on synchronized processes that integrate specific communication signals across a vast range of tissue groups. These crosstalks shift in their signaling dynamics, from early neurotrophic and adhesion cues to intermediate adipocyte-pericyte axes and eventually broader multi-tissue crosstalk that restores the homeostatic interactome.

Altogether, our analyses indicate three principal cellular regulatory hierarchies present in full-thickness wounds: (1) emergent transitional effectors (TE) – defined by cell states, such as Papillary I cells, that are sharply activated at discrete phase-transition intervals; (2) dominant signalers (DS), comprising populations such as Capillary ECs that dominantly drive temporally restricted signaling within a tissue compartment but do not constitute the most prominent emergent cell population within that tissue; and (3) Central Orchestrators (COs), a subset exemplified by Basal IV keratinocytes and PECs, which both exhibit the strongest emergence after injury and also assume the most prominent signaling roles within their respective tissue compartments (**Table 2**). We thus reasoned that this convergence of cell state emergence and signaling dominance reflects a core organizational architecture of deep wounds. Within this architecture, a subset of wound-emergent cellular hubs integrate paracrine cross-tissue signals to coordinate the transition of each healing phase.

### Wounding activates specialized tissue niches populated by Central Orchestrator populations

To investigate how wound-emergent cellular regulatory hierarchies are organized throughout the wound landscape we constructed WoundScape, a high-definition spatial transcriptomics (ST) atlas of full-thickness murine wounds at matched phase-resolved intervals (UW, D4PW, D7PW, D30PW) (**Figures 4A and S19-S22**). Using robust cell type decomposition (RCTD)^61^ with OWHA as a reference, we mapped the spatial distribution of all metaclusters and subclusters throughout the injury landscape (**Figures S19F-S19J)**. Next, we applied BANKSY clustering^62^ to delineate spatial neighborhood positions and organization across healing phases (**Figures S20A-S20H**). This revealed a highly ordered cellular neighborhood architecture in UW skin, dominated by large, well-segmented epidermal, hair follicle (HF), and muscle neighborhoods (**Figures 4B and 4F**). By D4PW, however, the neighborhood landscape underwent a dramatic reorganization, with marked increase in neighborhood densities (+12), but a near total loss (−4) of muscle-dominant neighborhoods (**Figures 4C and 4G**). Interestingly, at D4PW several new spatial clusters were dominated by epithelial-rich neighborhoods that progressively disappeared by D7PW (**Figures 4C-4D and 4G-4H**), indicative of rapid emergence of epithelial-dominant spatial hubs early in the wound response (**Figures 2C and 3B**). By D30PW, neighborhood composition dramatically shifted toward fibroblast-enriched spatial domains, including several neighborhoods (11, 14, 20, 27, 29) uniquely localized to the wound center (**Figures 4E, 4I**, **S20G-S20H, S20L and S21D**). These fibroblast-dominant neighborhoods spatially aligned within a late-stage scarring wound region (**Figures 4E**, **S20, and S21**). This is in sharp contrast to our multimodal single-cell data, which depicted a near-return to UW homeostasis by D30PW (**Figures 2C and S14C**). This revealed that although the skin’s transcriptional state may largely normalize by late maturation, its’ spatial organizational landscape remains substantially altered post-injury – an insight best captured by ST profiling and analysis. Correspondingly, we observed a general expansion in the total number of tissue neighborhoods across all WO timepoints, revealing that wounding induces a global reconfiguration of the cutaneous architecture into specialized injury-responsive niches with highly localized functions (**Figures 4F-4I**).

**Figure 4.**
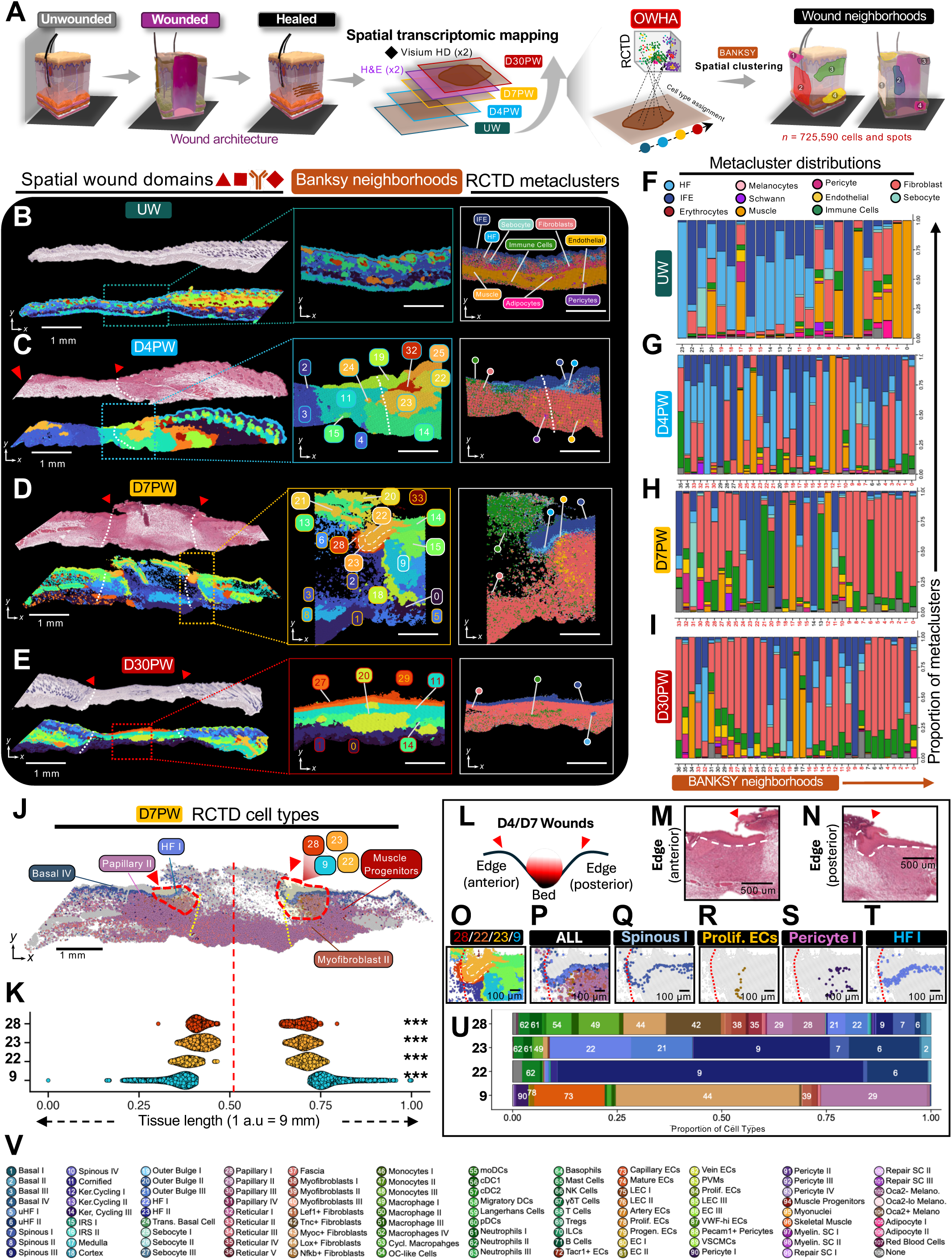
Spatial mapping of dominant master regulators uncovers highly localized wound emergent niches. **(A)** Schematic overview of WoundScape, the organ-scale spatial transcriptomic wound atlas. High-resolution 2 μm Visium HD data were generated for UW, D7PW, D15PW, and D30PW skin, totaling 532,066 spatial barcoded spots. These data were merged with the OWHA omnibus, yielding a comprehensive tetra-modal spatially resolved database encompassing 725,590 total cells and spots organized across all anatomical compartments of the skin. Histological H&E staining was combined with WoundScape spatial profiling to precisely align Visium HD-identified neighborhoods within defined cutaneous wound anatomical regions. **(B–E**) Spatial transcriptomic mapping of Banksy neighborhoods (bottom), and corresponding H&E sections (above), in 8 μm Visium HD sections from unwounded (**B**) (UW), (**C**) D4PW, (**D**) D7PW, and (**E**) D30PW. Middle insets show magnified views of local BANKSY neighborhoods clustering (clustering resolution = 0.5). Right insets depict corresponding RCTD-derived metacluster annotations for all discrete spatial domains within each section. Each section represents a technical replicate from the same biological specimen (UW = 113,587 spots; D4PW = 104,247; D7PW = 176,256; D30PW = 137,976). Red arrowheads indicate the initial wound edge in the suprabasal layer. White guidelines mark the initial subcutis wound boundaries. Scale bars represent 500 um or 1 mm as indicated. **(F–I)** Stacked bar plots showing the proportional metacluster composition within each BANKSY cluster for unwounded skin and wounded mouse skin (D4PW, D7PW, D30PW). Data represents two technical replicates derived from the same biological sample. Statistical similarity of metacluster compositions between replicates was assessed using a chi-square test with Monte Carlo permutation (100,000 simulations). BANKSY clusters that show significant concordance between technical replicates (*P* < 0.05) are highlighted in red font. **(J)** Visium HD localization of Dominant Signalers and Central Orchestrators, including Basal IV, Papillary II, HF I, Myofibroblasts II, and Muscle Progenitor clusters, along BANKSY neighborhoods proximal to the wound bed at D7PW. Red outlines mark wound-associated regions of interest, while yellow guidelines and red arrowheads denote suprabasal and subcutis wound edge boundaries, respectively. Scale bars, 1 mm. **(K)** Anatomical distance quantifications of wound emergent BANKSY clusters along the anterior-posterior transverse plane at D7PW. The x-axis represents arbitrary spatial units (1 a.u. = 9 mm) corresponding to anterior-posterior distance across the entire tissue section. Vertical redline demarcates the wound center. Data represents two technical replicates from the same biological timepoints. A variance-based localization test was used, and multiple testing correction was applied to p-values using the Benjamini–Hochberg procedure. (p < 0.05 = *, p < 0.01 = **, p < .001 = ***). **(L)** Schematic illustration summarizing the spatial geometry of the wound edge (anterior and posterior margins) as visualized at D7PW. Red arrowheads indicate typical location of wound boundaries from the adjacent wound bed. **(M-N)** High-magnification 20x H&E of the D7PW wound edge regions of interest (ROI). Red arrowheads indicate wound boundaries. Scale bars, 500 μm. **(O-T)** BANKSY spatial clustering of respective Dominant Signalers populations within the posterior wound edge ROI: (**O**) BANKSY cluster positions, (**P**) merged overlay of selected Dominant Signaler populations: (**Q**) Spinous I, (**R**) Proliferative Endothelial Cells, (**S**) Pericyte I, and (**T**) HF I. Each overlay highlights discrete but spatially organized domains at the wound front where reparative signaling networks converge. Scale bars, 100 μm. **(U)** Stacked bar plots showing fine cell-type composition of BANKSY clusters localized at the D7PW wound front. Data represents two technical replicates from the same biological timepoints. **(V)** Numbering and classification legend of fine cell types corresponding to panel **(U)**.

To understand how this spatial reorganization supports the activity of CO and DS populations, we examined their localization during the proliferation phase transition (D4PW–D7PW), when the largest increase in emergent cellular states was observed (**Figures S14A-S14C and S22**). This revealed a prominent enrichment of several wound-proximal neighborhoods, including BANKSY neighborhoods 9, 22, 23, and 28, at the D7PW wound edge (**Figures 4D and 4J-4N**). Interestingly, neighborhoods 9 and 28 were composed primarily of fibroblast and vascular–lymphatic populations, forming heterogeneous stromal-vascular hubs at the wound bed (**Figures 4O-4V and S20K**). In contrast, neighborhoods 22 and 23 consisted largely of basal epidermal, spinous, and HF-derived populations (**Figures 4O-4V and S20K).** Notably, Basal IV keratinocytes, one of the earliest and strongest emergent COs, constituted only a minor proportion of neighborhood 23 and 28 but was found in high abundance immediately adjacent to this neighborhood **(Figures 4U-4V and S22)**. This spatial positioning suggests that Basal IV cells appear to influence wound healing not merely by spatial dominance at the wound-edge regions, but largely by exerting influence through proximal signaling interactions.

### Basal IV keratinocytes orchestrate formation of a neurovascular regulatory hub to facilitate barrier restoration

Previous studies have shown that Sox6 expression enables skin keratinocytes to survive and maintain proliferation under acute environmental stress, such as UV exposure^63^. More recently, spatial coordination of diverse cutaneous cell populations, including epidermal and stromal cell movements, has been recognized as essential for an effective healing response across space and time^5,10,64^. To explore how Basal IV keratinocytes contribute to this coordinated multi-tissue repair process, we mapped their potential crosstalk and spatiotemporal distribution at the wound site (**Figures 5A and S23**). Beyond autocrine and epithelial-restricted signaling, Basal IV cells displayed the strongest cross-tissue interactions with endothelial cells, pericytes, and Schwann cells during the D4PW-to-D7PW transition **(Figure S23A).** Subcluster-level communication analysis confirmed that Basal IV most directly engaged PECs, Pericyte I, and Repair Schwann Cell II populations during the proliferation transitions (**Figures S23B-S23D**). ST analysis further validated this communication architecture and revealed that while Basal IV keratinocytes were broadly and evenly distributed across the unwounded IFE (**Figures 5B and 5F**), their localization was highly concentrated at the wound edge by D4PW and D7PW (**Figures 5C-5D and 5G-5H**). This enrichment placed them in direct spatial proximity with PECs, Pericyte I, and Repair Schwann Cell II populations that similarly accumulated at the wound edge (**Figures 5C-5D, 5G-5H and S24A-S24H)**. Notably, this coordinated Basal IV-neurovascular assembly dissipated by D30PW (**Figures 5E, 5I, S24D, and S24H)**, indicating that Basal IV keratinocytes occupy a transient wound-proximal niche that aligns with their wound-emergent CO roles. Together, these findings identify a strong physiological realignment specifically activated during the D4PW-to-D7PW proliferation transition. Given the spatial proximity of Basal IV keratinocytes with vascular-associated cells, particularly PECs, during the D4PW-to-D7PW transition, we next interrogated signaling programs facilitating interaction of Basal IV and surrounding PECs (**Figures 5J and 5K)**. This revealed prominent activation of the trans-synaptic adhesion–GPCR axis *Tenm4–Adgrl2*, as well as the neuronal-vascular patterning axes *Sema3c–Plxnd1* and *Sema3c–Nrp2/Plxna4* during this interval (**Figures 5K and S23E-S23H)**. WoundScape mapping showed strong, localized expression of *Tenm4* exclusively along the basal epidermis, whereas *Adgrl2* was diffusely expressed both in the basal IFE as well as the dermal wound edge (**Figures S24I-S24O)**. Importantly, SEMA3C pathway components (*Sema3c, Nrp1, Nrp2, Plxna4*) displayed strong colocalized expression at the wound edge at D7PW, with *Sema3C* itself being strongly expressed both within the basal epidermis and in the subcutaneous fascia compartment (**Figure 5L**). Single molecule RNA fluorescent *in situ* hybridization (smFISH) analysis further validated these findings and confirmed strong colocalization of *Sema3C* with *Sox6^+^* cells in the basal epidermis by D4PW compared to UW skin (**Figures 5M-5P and S25**). Collectively, these data suggest that Basal IV keratinocytes orchestrate the formation of a specialized, contact-dependent neurovascular niche at the wound edge via SEMA3C and other organizing ligands to tune endothelial outgrowth, pericyte stabilization, and Schwann cell-mediated neurite guidance.

**Figure 5.**
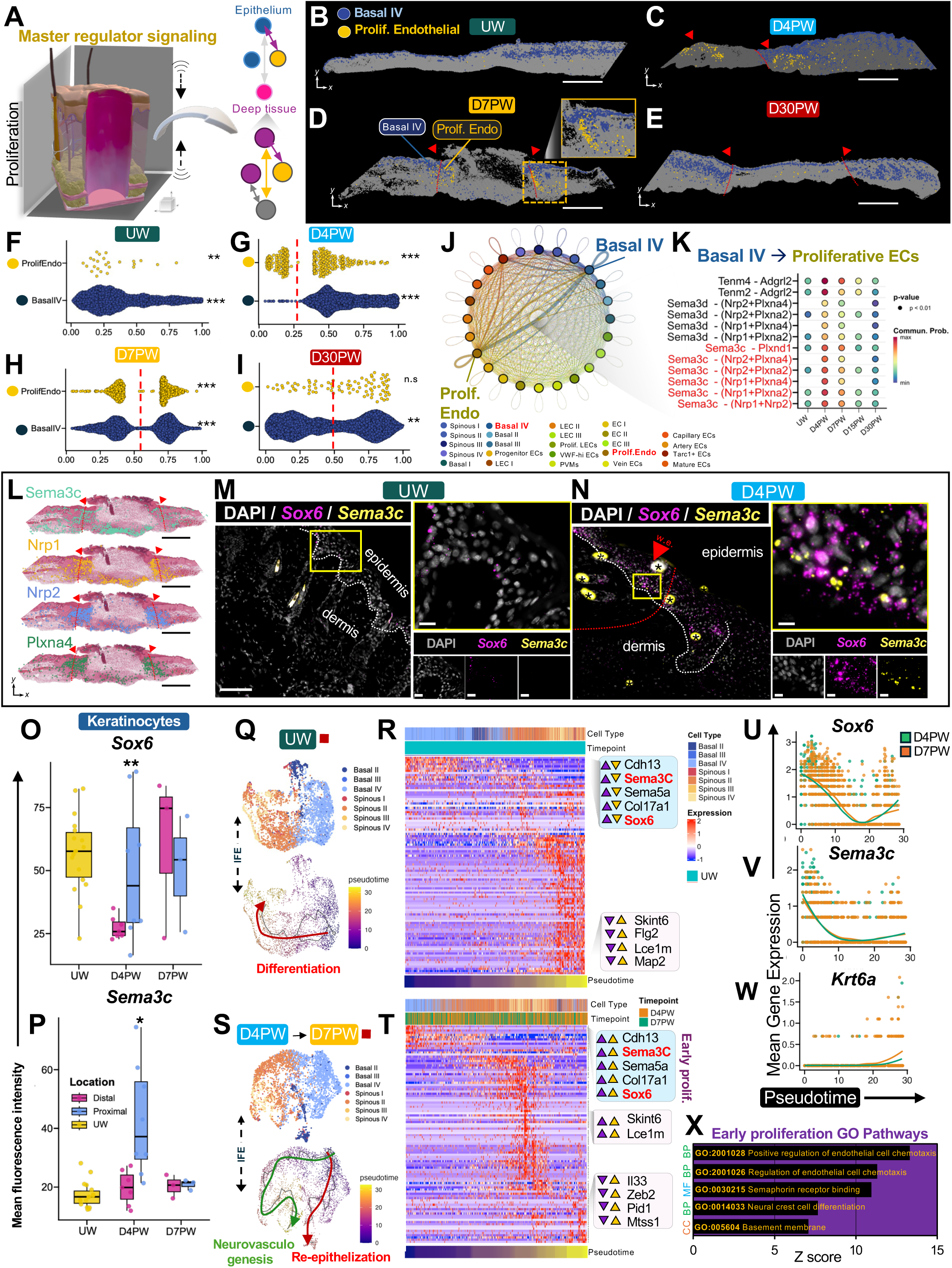
Sox6⁺ basal keratinocytes are master controllers driving wound re-epithelization through dynamic epidermal–neurovascular crosstalk. (**A**) Schematic illustration of cross-tissue wound signaling between epidermal Central Orchestrators and deep tissue population populations. (**B–E**) Visium HD spatial localization of Basal IV keratinocytes (blue) and proliferative endothelial cells (yellow) across the healing time course (**B**) UW, (**C**) D4PW, (**D**) D7PW, (**E**) D30PW. Images represent one biological replicate, with one technical replicate shown for UW. Red arrowheads indicate the suprabasal wound edge; red guidelines denote subcutis wound boundaries. Yellow boxes indicate magnified insets highlighting Basal IV–endothelial interactions at the wound edge. Scale bar, 1 mm. (**F–I**) Quantification of anatomical distance between Basal IV and proliferative endothelial cells along the anterior–posterior wound axis (1 a.u. = 9 mm) at (**F**) UW, (**G**) D4PW, (**H**) D7PW, (**I**) D30PW. Data represent two technical replicates from one biological sample per timepoint. Significance was assessed using a variance-based localization test with Benjamini–Hochberg correction (* p < 0.05; ** p < 0.01; *** p < 0.001). (**J–K**) CellChat-inferred signaling network (**J**) and corresponding ligand–receptor pathways (**K**) transmitted between Basal IV keratinocytes and endothelial cells. Edge width denotes interaction strength. (**L**) Visium HD spatial transcriptomic expression of *Sema3c* pathway components (*Sema3c*, *Nrp1*, *Nrp2*, *Plxna4*) overlaid on corresponding H&E sections at D7PW. Scale bar, 1 mm. (**M–N**) Single-molecule RNA FISH (smFISH) immunofluorescence showing *Sox6* (magenta) and *Sema3c* (yellow) mRNA transcripts with DAPI (white) in (**M**) UW and (**N**) D4PW skin. Boxed regions highlight an Sox6-high epidermal zone. (Right) zoomed in images of the Sox6-high zone with single-channel panels shown. Wound edge (w.e.) indicated by red arrowheads and subcutis wound boundaries indicated by red lines. Scale bars, 100 µm (full images) or 10 µm (zoomed insets). (**O–P**) Boxplots quantifying mean fluorescence intensity of (**O**) *Sox6* and (**P**) *Sema3c* smFISH signals across timepoints (UW, D4PW, D7PW), comparing unwounded, distal, and proximal wound regions. UW includes three biological replicates, D4PW two biological replicates, and <u>D7PW one biological replicate</u>. Statistical testing performed using a two-sided Wilcoxon test. Variability is represented using the interquartile range (IQR). Statistical significance was determined using a Wilcoxon rank-sum test (p < 0.05 = *, p < 0.01 = **). (**Q**) UMAP visualizations and corresponding pseudotime ordering of UW IFE keratinocyte subclusters from OWHA snRNA-seq. *n* = two biological replicates. (**R**) Heatmap of top 100 pseudotime-associated genes in UW snRNA-seq keratinocytes ordered by cluster and pseudotime. Arrowheads mark genes enriched in early (purple) vs. late (yellow) pseudotime. (**S**) UMAP visualizations and corresponding pseudotime ordering of D4PW–D7PW IFE keratinocyte subclusters showing branching into re-epithelization (red) and neurovasculogenesis (green) lineage trajectories. *n* = two biological replicates per timepoint. (**T**) Heatmap of the top 100 pseudotime-associated genes expressed along the neurovasculogenesis pseudotime trajectories in D4PW–D7PW keratinocytes, ordered by cluster and timepoint. Arrows indicate early (purple) and late (yellow) pseudotime gene signatures. (**U–W**) Mean pseudotime expression profiles of (**U**) *Sox6*, (**V**) *Sema3c*, and (**W**) *Krt6a* along the D4PW–D7PW proliferative pseudotime trajectory. (**X**) Gene Ontology terms among genes upregulated at early pseudotime stages of the neurovasculogenesis trajectory, subdivided into molecular function (MF), biological process (BP), and cellular component (CC) categories.

Next, to better define the functional trajectories adopted by Basal IV keratinocytes during healing, we performed pseudotime analysis spanning UW–D7PW (**Figures 5Q–5T**). In UW skin, Basal IV cells exhibited strong expression of *Sox6* and *Sema3c* and progressed along a canonical epithelial differentiation trajectory characterized by increased expression of *Skint6*, *Flg2*, and *Lce1m* accompanied by gene programs associated with keratinization and cornified envelope formation (**Figures 5Q, 5R and S26A**). Correspondingly, pseudotime ordering revealed direct routes from Basal IV cells to a *Sox6^+^* Keratinocyte Cycling III population and a *Sox6^−^* spinous cell population (**Figures 5Q and S26F-S26I**). This mirrors recent reports of slow-cycling *Sox6*^+^ keratinocytes giving rise to rapidly cycling keratinocyte populations in the mouse and human skin adapting to UV exposure^65^. Notably, we found that this canonical slow cycling to spinous differentiation route also remained intact after wounding. However, a second wound-induced trajectory emerged, marked by elevated expression of *Serpinb7*, *Adgrl2*, *Nebl*, and *Grip1* and enriched for axonogenesis– and neurogenesis-associated GO terms (**Figure 5S-5T and S26C-26D**). This suggests a sequential keratinocyte-driven fate specification program involved in vascular stabilization followed by reinnervation during wound resolution. Strikingly, these fate trajectories culminated in the generation of *Krt16⁺* wound-edge keratinocytes (**Figure 5T-5W, S26A-S26B**), indicating that the neurovasculogenic signaling state adopted by Basal IV cells is critical for the wound-induced re-epithelialization required to regenerate barrier epithelium. These findings also support prior reports of enhanced angiogenesis activity in Sox6^+^ keratinocytes^65^ and of a glial-rich niche forming at early stages of repair^66^, indicating that Basal IV cells serve dual, context-dependent roles: in homeostasis, they support barrier maintenance through canonical terminal differentiation, but after injury they generate wound-front *Krt16*⁺ cells and adopt neurovascular functions that help restore periwound vasculature and tissue integrity (**Figures 5Q–5X and S26**).

### Basal IV keratinocytes participate in human wound healing but are excluded from standard scRNA-seq datasets

To examine the broader applicability of OWHA in human biology and determine whether *Sema3c*-expressing Basal IV keratinocytes are conserved in human skin repair, we constructed a Cross-species Organ-Scale Wound Atlas (COWA) by integrating recently published human scRNA-seq datasets^24^ with OWHA at matching time points (UW, D1PW, D7PW, D30PW) (**Figure 6A**). This integrated dataset encompassed 236,930 cells and enabled direct comparison of conserved and divergent cellular lineages and programs across human and murine wounds (**Figures 6A-6D and S27**). COWA revealed a prolonged remodeling phase in human wounds compared to mouse wounds, with sustained expression of ECM remodeling genes as late as D30PW (**Figures S27B-S27C**). In addition, COWA revealed that several populations robustly captured in OWHA were largely absent in the human scRNA-seq data, including skeletal muscle, myonuclei (panniculus carnosus (PC) muscle), adipocytes, fascia, several fibroblast subsets, Schwann cell subsets, lymphatic endothelial cell subsets, and sebocytes (**Figures 6C-6F and S27A**). With the exception of mouse-specific PC cells, these differences reflect known limitations of droplet-based scRNA-seq in recovering large, fragile, or low-RNA cell types^23,67^, rather than true biological differences in tissue composition between the two species. To further quantify inter-species fidelity between the mouse OWHA dataset and human scRNA-seq atlas, we performed anchor-based cross-species projections of skin cell states and visualized lineage correspondence using Sankey diagrams (**Figure 6E; Table S4**). This revealed highly conserved cell type correspondences between both species in which most major mouse skin lineages, including IFE and endothelial clusters, aligned to their analogous human lineages in their transcriptional expression (**Figure 6E; Table S4**). However, specificity of these correspondences varied more at the subcluster level (**Figures 6E, S27D-S27F, and S28A**).

**Figure 6.**
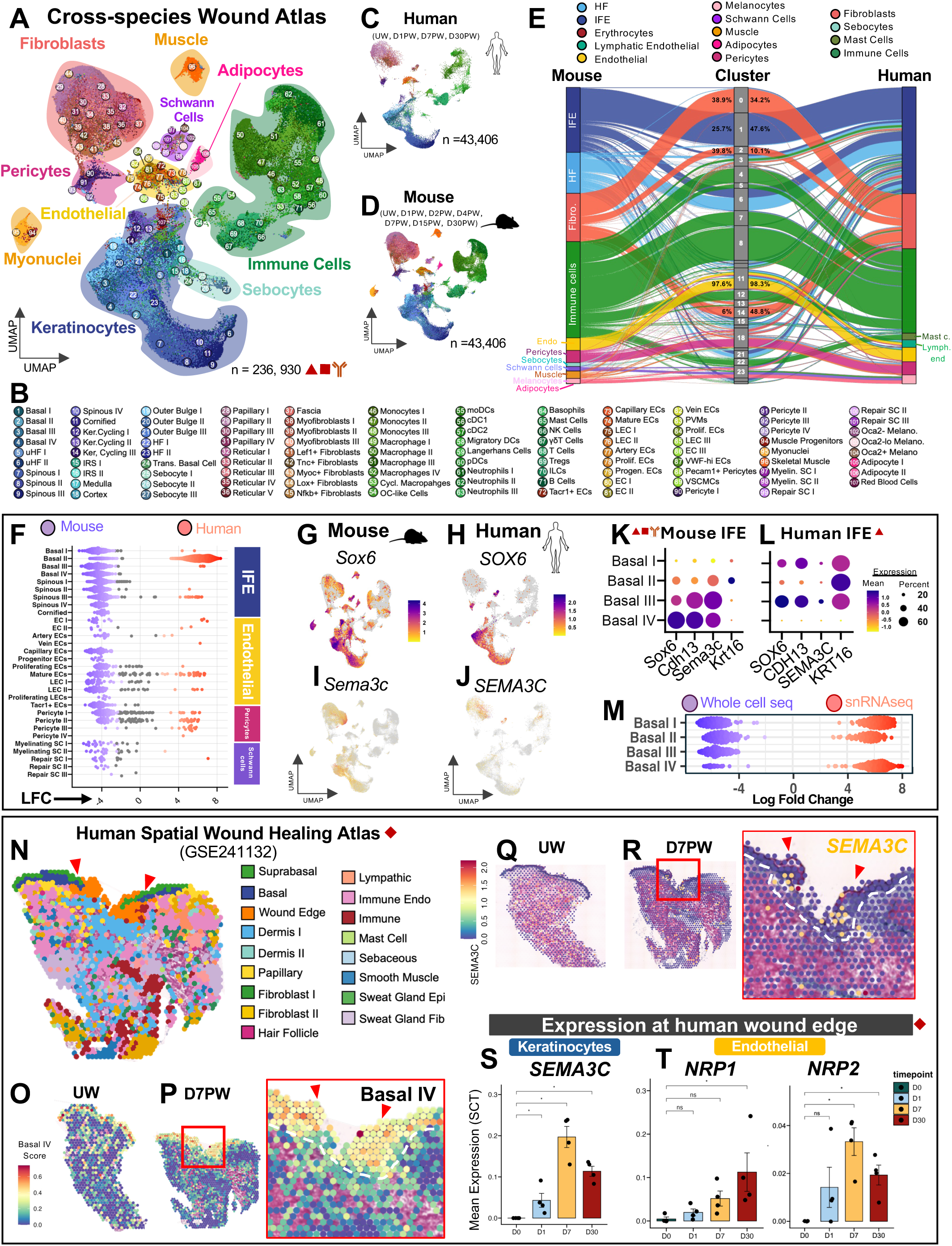
Sox6^+^ Basal IV keratinocytes are active in human wound healing but entirely missed by conventional scRNA-seq capture. (**A**) UMAP embedding of the fully integrated cross-species wound-healing atlas (COWA) containing 236,930 single cells. COWA integrates the multimodal OWHA dataset with the human wound healing dataset (GSE241132), generating a total of 40 sequencing runs (28 murine, 12 human). (**B**) Annotation legend displaying all 107 fine-grained subcluster identities represented in COWA. (**C–D**) UMAP projections showing (**C**) human (n=12) and (**D**) mouse (n=28) wound datasets downsampled to 43,406 cells each for comparable visualization of interspecies cell capture differences. (**E**) Sankey plot showcasing proportional representation of major cell types between mouse and human samples across metaclusters, highlighting shared and species-specific tissue compositions. Percentages display proportion of each respective metacluster assigned to integrated cross-species Harmony clusters. (**F**) MILO differential abundance analysis of select IFE keratinocyte, endothelial, pericyte, and Schwann cell subclusters states significantly enriched in mouse (blue) or human (red) samples (FDR = 0.15). Mouse-enriched (>0 LFC) states represent populations preferentially detected in by multimodal murine atlasing. (**G–H**) UMAP feature plots depicting (**G**) mouse *Sox6* and (**H**) human *SOX6* expression. (**I–J**) UMAP feature plots of (I) mouse *Sema3c* and (J) human *SEMA3C*, showing substantially reduced *SEMA3C* detection in human wound-healing scRNA-seq datasets. (**K–L**) Dot plots of Basal IV–associated markers (*Sox6*, *Cdh13*, *Sema3c*) and the wound-induced keratinocyte gene *Krt16* in IFE keratinocytes from (**K**) the murine OWHA dataset and (**L**) the human wound dataset. Dot size indicates the percentage of expressing cells; color intensity denotes mean expression level. (**M**) Beeswarm plot of MILO differential abundance comparing whole-cell (blue) versus single-nucleus (red) RNA-seq, highlighting modality-specific biases in cell type capture (FDR = 0.15). (**N**) Visium spatial transcriptomic feature plot of D7PW human wound tissue (GSE241132), annotated by regional labels as defined Liu et al. 2024. Red arrowheads mark the wound edge. (**O–P**) Quantification of Basal IV module scores in (**O**) unwounded and (**P**) D7PW human tissue sections (calculated using *AddModuleScore()*). Red box highlights wound edge region zoomed-in in the right panel. Red arrowheads denote the wound edge; white dashed lines outline the epidermal-dermal boundary. (**Q–R**) Quantification of *SEMA3C* pathway signature scores in (**Q**) unwounded and (**R**) D7PW human tissue sections. Red box highlights wound edge region zoomed-in in the right panel. Red arrowheads denote the wound front; white dashed lines indicate the epidermal-dermal boundary. (**S–T**) Mean expression of (***S***) *SEMA3C* in keratinocytes and (**T***) NRP1* and *NRP2* in endothelial cells at the human wound edge across healing timepoints (D0PW, D1PW, D7PW, D30PW). Each dot represents the mean signal per sequencing run. Statistical significance was assessed using a Wilcoxon rank-sum test relative to D0/UW (* p < 0.05; ** p < 0.01). *n*= 4.

We thus sought to determine whether *Sema3c-*expressing Basal IV keratinocytes were conserved in human skin. Strikingly, Basal IV cells were largely missing from the human datasets (**Figures 6F and S28**). To determine whether this reflected technical limitations in transcriptional gene capture, we examined the expression of *Sox6* and *Sema3c,* two key markers defining the Basal IV cell state (**Figure S8**). Both genes displayed robust expression in mouse and human keratinocytes (**Figures 6G-6L**), and *Sema3C* was consistently detected in COWA datasets, though its epithelial expression was comparatively lower in human samples (**Figures 6I–6L**). Strikingly, although canonical Basal IV marker genes were captured in both species, the Basal IV cell population was entirely absent as a discrete cell group in the human scRNA-seq atlas (**Figure 6L**). Given the known limitations of scRNA-seq in capturing the full cell type heterogeneity of the epidermis, particularly in human samples^68^, we next examined whether these results reflected technical deficiencies in cell type capture. While most basal cell groups were detected in both species, Basal IV, Spinous I, and Cornified cells were entirely absent in human scRNA-seq atlas data. Moreover, this data revealed low capture rates of several fragile lineages, such as vascular-lymphatic and glial populations, including PECs and Myelinating Schwann cells (**Figures 6F and S28**). Collectively, these findings suggest that the apparent absence of Basal IV keratinocytes in human datasets likely reflects a technical artifact rather than a failure in gene capture or species-specific scarcity. Such artefactual biases are consistent with recent reports of dissociation-induced cell loss in skin and transcriptional perturbation during enzymatic tissue digestions^68–72^. Given that such bias would considerably alter interpretation of experiments investigating tissue wound healing, we systematically examined modality-specific differences in cell abundances and transcriptomic capture, between whole cell and snRNA-seq batches included in OWHA (**Figures 6M, S29A-S29D**). Notably, among all basal cell groups, Basal IV cells were only robustly captured by snRNA-seq (**Figure 6M**).

Because whole-cell sequencing preparations of mammalian skin require thorough live-cell enzymatic digestion, while snRNA-seq sample preparation employs flash-frozen tissue void of tissue-dissociating enzymatic exposure^68,73,74^ (see **Methods**), we reasoned that differences in sample preparation could substantially influence the representation of the stress-responsive Basal IV population. We thus quantified the expression of stress-induced transcripts in whole cell and snRNA-seq samples at matching timepoints (**Figures S29E-S29H and S30A-S30B**). Quantification of stress-associated genes^69^ detected by all modalities used in OWHA (**Table S5**) revealed that snRNA-seq samples displayed markedly lower stress scores at the UW state relative to time-matched whole-cell datasets (**Figures S29 and S30A**). Although all modalities showed increased stress signatures post-injury, the fold-change increase was substantially lower in snRNA-seq than in whole-cell sequencing preparations (**Figures S29E-S29H and S30A-S30D**). UMAP embedding further revealed elevated stress scores across diverse cell clusters in whole-cell sequencing samples (**Figure S29E**). This indicates that enzymatic dissociation disproportionately perturbs stress-responsive cell states which may selectively distort the representation of fragile or activation-prone populations such as Basal IV cells in wound atlases. To further examine this possibility, we calculated stress scores within the IFE, a tissue compartment well captured by all modalities (**Figure S4G**). Consistent with our prior findings on aggregated clusters, IFE populations derived from snRNA-seq exhibited statistically significant but modest increases in stress scores at the UW state relative to all injury timepoints (**Figure S30C**). Conversely, IFE clusters derived from whole-cell sequencing displayed significantly greater fold change increases in stress scores at the UW state relative to all injury timepoints (**Figure S30C**). We next analyzed stress scores in skin cells obtained from ST modalities generated from Formalin-fixed paraffin-embedded (FFPE) tissues, which are also never exposed to live-cell enzymatic digestions. Expectedly, ST datasets from both human and mouse skin revealed elevated stress scores after injury (**Figures S30E-S30J, S31A, and S31B**); however, these stress signatures were spatially restricted to the wound bed and predominantly localized to immune cells rather than broadly distributed across all clusters (**Figures S30E-S30N**), as observed in whole-cell sequencing datasets. Moreover, ST stress scores remained low in the UW samples and showed modest induction patterns that more closely resembled those observed in snRNA-seq (**Figures S30**).

Importantly, spatial mapping of the Basal IV cell gene signature in human wound ST samples revealed a distinct subpopulation localized near the wound edge, with prominent *SEMA3C* and *NRP2* colocalization observed in D7PW wounds (**Figures 6N-6T**). This spatial and temporal ordering closely matched the murine Basal IV wound activation pattern, where *Sema3c*-expressing Basal IV cells were enriched along the wound margins **(Figures 6N-6T, S24A-S24D, and S31C-S31N**). Altogether, these findings support the presence of a conserved, epidermal-mediated SEMA3C-NRP1/NRP*2* signaling axis in both mouse and human wound repair and highlight the critical need of incorporating multiple transcriptomic modalities to overcome capture bias and dissociation-driven artifacts in wound atlases.

### Topical Sema3C administration rescues re-epithelization in diabetic skin ulcers

A major defect in human wound healing arises from wound neuropathy driven by chronic diabetic metabolic dysregulation that impairs protective skin sensation and neurovascular coupling, thereby promoting the development of chronic, nonhealing diabetic foot ulcers (DFU)^75,76^. We thus choose to investigate the functional impact of SEMA3C in shaping the neurovascular niche at the wound edge by first analyzing the transcriptomic signature of human ischemic wounds using a human DFU dataset.^25^ *SEMA3C* expression in DFU keratinocytes was significantly downregulated in both healing and non-healing diabetic ulcers relative to healthy skin (**Figure 7A**). Strikingly, while keratinocyte *SOX6* expression was most significantly depleted in diabetic ulcers, its expression was also significantly reduced in diabetic but non-ulcerated skin, indicating that chronic metabolic dysregulation resulting from diabetes impairs baseline Basal IV *SOX6* activity (**Figures 7B and S31O-S31R**). Additionally, endothelial cells in non-healing human ulcers showed reduced expression of *NRP1* and *NRP2*, but healing ulcers had significantly higher *NRP2* levels relative to non-healing ulcers (**Figures 7C-7D**). Similarly, healing ulcers were able to significantly upregulate *KRT6A* compared to non-healing ulcers (**Figure S31Q**). Together, these findings suggest that diabetes disrupts the Basal IV SEMA3C–NRP1/NRP2 signaling axis, impairing re-epithelialization and predisposing diabetic skin to nonhealing ulcerations. This supports previous findings which demonstrated that exogenous SEMA3C enhances corneal reinnervation after injury in diabetic mice^77^ and that SEMA3C-NRP2 signaling improves healing of DFUs in rats^78^. The significant increase of *NRP2* and *KRT6A* in samples from healing ulcers suggested that compensation of the SEMA3C–NRP1/NRP2 signaling axis could restore healing potential in diabetic ulcers. To investigate whether SEMA3C treatment could rescue defective inflammatory-to-proliferative transition in diabetic skin *in vivo*, we generated large full-thickness wounds^79^ in *Lepr^db^*/J mice which created DFU-like ulcers (**Figure 7E**). Expectedly, *Lepr^db^*/J mice exhibited markedly delayed healing compared to wild-type controls (**Figure 7J**). Notably, topical delivery of recombinant mouse SEMA3C (mSema3C) or human SEMA3C (hSema3C) significantly accelerated wound closure at timepoints corresponding to active keratinocyte migration and re-epithelialization (D7PW) (**Figures 7F–7J**). Importantly, both hSema3c– and mSema3c-treated wounds also displayed dense periwound angiogenic sprouting, indicating increased revascularization in both treatment groups relative to vehicle controls (**Figures 7K–7O**). Altogether, these findings reveal that while diabetes impairs epidermal-derived SEMA3C production and the SEMA3C–NRP1/NRP2 signaling circuitry required for timely proliferation phase transition, topical administration of recombinant SEMA3C can restore this signaling deficit, rescuing periwound angiogenesis and re-epithelialization of DFUs (**Figure 7P**).

**Figure 7.**
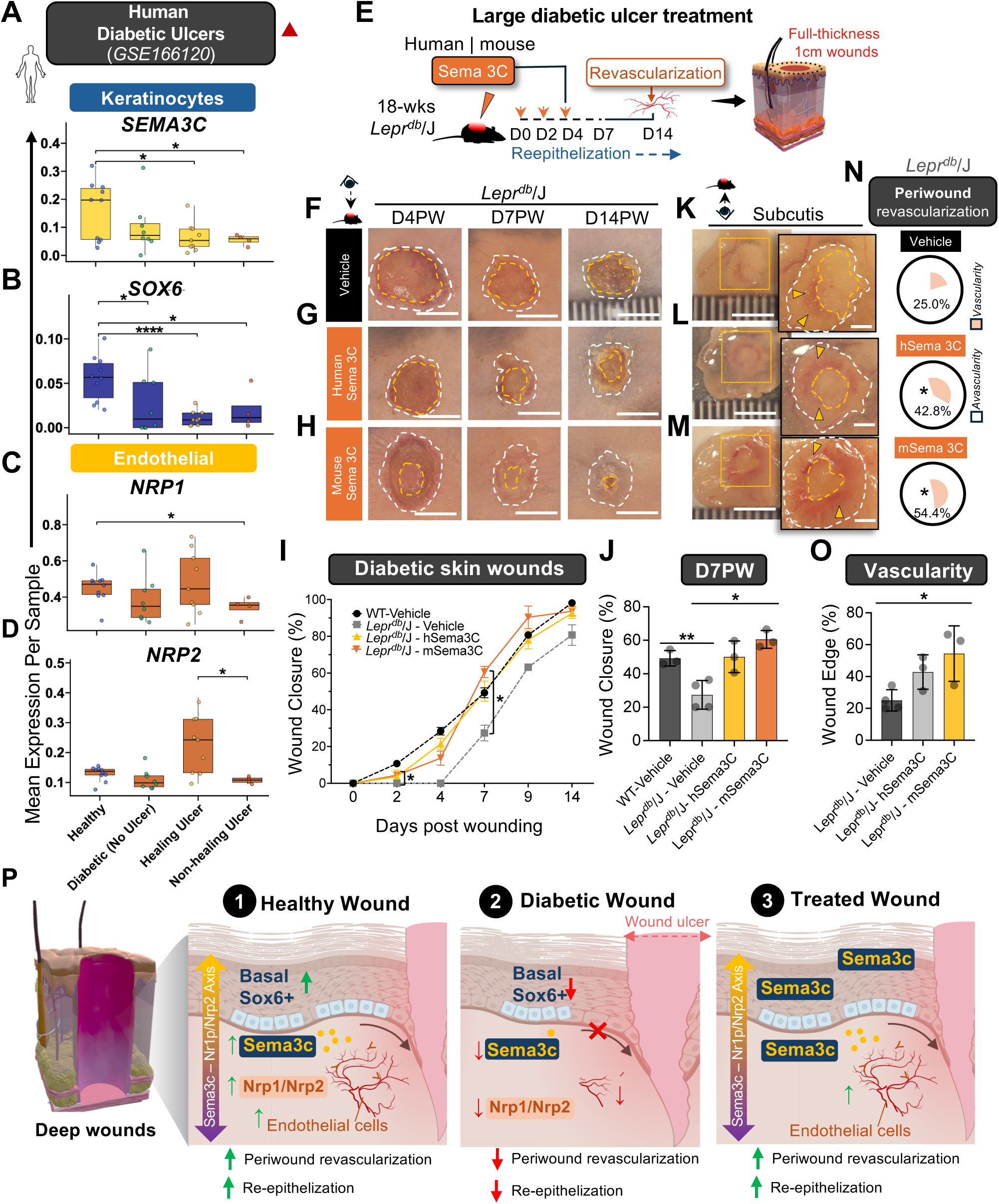
Epidermal *SEMA3C* is impaired in human diabetic wounds, but its topical reintroduction restores re-epithelization of diabetic skin ulcers *in vivo*. **(A–D)** SCT-normalized mean expression of (**A**) *SEMA3C* and (**B**) *SOX6* in human keratinocytes, and (**C**) *NRP1* and (**D**) *NRP2* in human endothelial cells (GSE166120). Data shown for healthy skin, diabetic skin without ulcers, healing diabetic foot ulcers, and non-healing diabetic foot ulcers. Each dot represents a sequencing run. Healthy control *n* = 11, diabetic non-ulcer *n* = 15, diabetic healing ulcer *n* = 9, diabetic non-healing ulcer *n* = 4. Statistical significance was determined using a Wilcoxon rank-sum test (p < 0.05 = *, p < 0.01 = **). (**E**) Schematic overview of the large diabetic wound ulcer model. 1-cm full-thickness wounds were generated in *Lepr^db^/J* diabetic mice before topical treatment with recombinant *SEMA3C* protein (human or mouse) or vehicle control at D0, D2, and D4 post-wounding. Wounds were harvested for closure and revascularization analysis at D14PW. **(F–H)** Representative dorsal wound images from **(F)** vehicle, **(G)** human *SEMA3C*, and **(H)** mouse *Sema3c* treatment groups at D4PW, D7PW, and D14PW. White dashed lines mark initial wound boundaries; yellow lines indicate wound diameter at imaging analysis. Images represent three independent biological replicates per condition (n=3). **(I)** Quantification of mean wound closure rate in WT and *Lepr^db^/J* mice treated with vehicle, human *SEMA3C*, or mouse *Sema3c*. Measurements taken at days D0, D2, D4, D7, D9, and D14PW. Data represents the mean of three independent biological replicates per condition (n=3), besides *Lepr^db^/J –* Vehicle which is two biological replicates. Statistical significance was determined by two-way ANOVA (*p* < 0.05 = *, *p* < 0.01 = **). Significance is marked between *Lepr^db^/J-*mSema3c treated vs *Lepr^db^/J –* Vehicle. **(J)** Quantification of D7PW wound closure rate in WT and *Lepr^db^/J* mice treated with vehicle, human *SEMA3C*, or mouse *Sema3c*. Data represents the mean of three independent biological replicates per condition (n=3), besides *Lepr^db^/J –* Vehicle which is two biological replicates. Statistical significance was determined by two-way ANOVA (*p* < 0.05 = *, *p* < 0.01 = **). **(K-M)** Representative ventral-view images of subcutaneous wounds in *Lepr^db^/J* mice at D14PW, treated with **(K)** vehicle, **(L)** human *SEMA3C*, or **(M)** mouse *Sema3c* for each condition. Boxed region highlights subcutaneous wound bed and periwound region. (*Right)* Enlarged view of the boxed region. White dashed lines mark the original wound boundaries, and yellow lines indicate the central wound bed. Yellow arrowheads indicate regions with active revascularization. Images represent three independent biological replicates per condition (n=3). **(N-O)** Periwound revascularization coverage in *Lepr^db^/J* mice at D14PW after treatment with vehicle, human *SEMA3C*, or mouse *Sema3c*. **(O)** Proportion of periwound revascularization at D14PW in each treatment group. Data represents the mean of three independent biological replicates per condition (n=3). Statistical significance was determined by two-way ANOVA (*p* < 0.05 = *, *p* < 0.01 = **). **(P)** Proposed model summarizing the role of the Sema3c*–*Nrp1/Nrp2 signaling axis in healthy versus diabetic skin wound healing. In healthy wounds, activation of *Sox6*⁺ Basal keratinocytes promote Sema3c secretion and paracrine signaling to *Nrp1/Nrp2*⁺ endothelial cells, facilitating re-epithelialization and revascularization of the wound bed. This circuit is disrupted in diabetic wounds, leading to reduced *Sox6*⁺ Basal keratinocyte activation and decreased Sema3c expression. This results in impaired re-epithelization and periwound vascular repair. However, topical reintroduction of Sema3C restores periwound revascularization and wound re-epithelization in diabetic skin ulcers.

### Discussion

In this study, we establish the first Organ-Scale Wound Healing Atlas (OWHA), a joint multimodal, spatiotemporal reference map of full-thickness murine skin repair that integrates scRNA-seq, snRNA-seq, CITE-seq, and high-resolution spatial transcriptomics across all major healing phases (**Figure 1A**). Profiling over 725,000 cells, this wound omnibus captures 107 skin subpopulations, overcoming long-standing limitations in temporal coverage, cell-type recovery, and spatial cell signaling. OWHA unveils wound healing as a coordinated, organ-level process governed by discrete cellular and molecular inflection points rather than a gradual continuum. These phase transitions occur synchronously across epithelial, stromal, vascular, perineural, and deep tissue compartments while intermediate healing phases remain comparatively stable. This inflection model refines classical linear paradigms of repair and provides a systems-level explanation for how temporally discrete biological programs coordinate transitions between inflammation, proliferation, and maturation to drive organ-scale restoration.

OWHA also reveals that wound healing is organized around conserved, wound-emergent cellular hierarchies and cross-lineage regulatory hubs we term Central Orchestrators (COs). High-resolution ST analysis demonstrates that wounding induces a global reorganization of tissue architecture into specialized, injury-responsive niches populated by COs. During the inflammatory-to-proliferative (D4PW-D7PW) transition, these niches assemble into discrete spatial transition interfaces at the wound edge where epithelial, stromal, vascular, and neural domains converge through coordinated ligand-receptor exchange. These organ-level analyses reframe wound healing not as a series of isolated lineage programs but as a tightly regulated relay of interconnected cell states acting across tissue layers and compartments. Among these CO populations, *Sox6*⁺ Basal IV keratinocytes emerged as a wound-responsive cell group that synchronizes epithelial stress responses with neurovascular remodeling by occupying a transient, anatomically restricted niche at the wound edge. Mechanistically, this process occurs through activation of a SEMA3C-dependent signaling axis with PECs that includes SEMA3C-NRP1/NRP2 and SEMA3C-PLXN (**Figures 5K-5P**), pathways classically implicated in developmental axon guidance and vascular patterning^77,80,81^ but acutely redeployed during adult skin repair (**Figure 5**). Furthermore, our findings demonstrate that during homeostasis, Basal IV keratinocytes follow canonical differentiation trajectories toward spinous and cornified states, supporting epidermal maintenance. Following injury, however, they exhibit injury-dependent plasticity, where a distinct wound-induced trajectory emerges that is enriched for neurogenic and vasculogenic programs and culminates in *Krt16*⁺ wound-edge keratinocytes (**Figures 5Q-5X**). This bifurcation is reminiscent of the reprogramming described for SOX6^+^ keratinocytes under UV stress^63^ and underscores how injury repurposes epithelial differentiation fates to meet regenerative demands. The multi-functional plasticity observed in basal stem cells may explain why defective re-epithelization and failure to restore these populations leads to such vast detrimental effects in chronic-ulcers and other wound complications.

A major finding was that several critical wound-associated populations–including Basal IV keratinocytes, adipocytes, Schwann cells, fascia, and vascular subsets–were underrepresented in scRNA-seq datasets yet robustly detected using snRNA-seq and spatial transcriptomics, revealing that their apparent absence largely reflects dissociation-driven technical bias rather than true biological divergence (**Figure 6**). In particular, Basal IV keratinocytes were selectively lost during enzymatic dissociation, masking their contribution to wound repair in conventional scRNA-seq atlases – an artifact corrected by spatial and snRNA-seq modalities. Cross-species integration validated the conservation of Sox6⁺ keratinocytes in human IFE and demonstrated their apparent absence from prior human scRNA-seq atlases arose from dissociation-driven dropout. These findings reveal that enzymatic dissociation inflates stress-responsive transcriptional programs in skin cells, thereby biasing cell-state representations in skin wound atlases. OWHA overcomes these constraints by integrating single-cell, single-nucleus, epitope-based proteomic, and spatial transcriptomic approaches in a phase-resolved manner, which provides a cohesive organ-level framework for resolving lineage programs, spatial niches, and inter-tissue communications during repair.

Importantly, probing human data demonstrated that the SEMA3C–NRP1/NRP2 axis is disrupted in diabetic ulcers, and *SOX6* expression is reduced even in non-ulcerated diabetic keratinocytes, indicating that Basal IV keratinocyte activity is disrupted in diabetic skin even prior to overt ulceration. This emphasizes recent evidence of a necessity for robust intercellular and cross-tissue communication for effective skin maintenance and wound response.^24,66,82^ Notably, topical recombinant SEMA3C accelerated wound closure, restored periwound angiogenesis, and promoted re-epithelialization in diabetic mouse models, validating this conserved epithelial-driven circuit as a therapeutically actionable target necessary for timely transition from inflammation to proliferation. Moreover, while the late maturation phase (D15PW–D30PW) exhibited minimal changes in cell-state composition, the spatial architecture of the integument remained distinctly remodeled, particularly within scar tissue. This divergence between transcriptional restoration and architectural recovery highlights the importance of the spatial wound context in defining long-term healing outcomes.

### Limitations

One limitation of our study is that very early injury timepoints (D1PW, D2PW) lacked sufficient biological sampling, which limits the statistical robustness of conclusions drawn from these stages. Furthermore, OWHA incorporates data from a mosaic of sequencing chemistries used to profile transcriptomes across batches. Although no single sequencing chemistry was disproportionately represented (**Figure S2F**), this heterogeneity may introduce batch-related variabilities that could be difficult to detect. Finally, while Sox6⁺ Basal IV cells strongly upregulate *Sema3C*, and its promoter contains a SOX-binding motif, direct transcriptional regulation remains to be experimentally validated in mouse and human skin^83–85^.

Overall, OWHA achieves a major advance in our understanding of wound physiology and provides an integrated 4D blueprint of mammalian wound repair that defines a core organizational logic of organ-scale tissue reconstruction. By uncovering conserved regulatory circuits with direct therapeutic relevance, this wound omnibus provides both a foundational resource for wound biology and a mechanistic framework for understanding and targeting impaired healing in mammalian systems. Our approach demonstrates the power of multimodal organ-level mapping to reveal previously inaccessible biology and furnishes a community resource for exploring lineage-specific programs, signaling circuits, and druggable targets across the wound healing continuum.

## MATERIALS AND METHODS

### Mouse wounding model

C57BL/6J wild-type mice were purchased from The Jackson Laboratory and maintained under specific pathogen-free conditions in the barrier facility at Columbia University Irving Medical Center. All mice were maintained under standard conditions with free access to food and water and a 12-hour light/dark cycle. Mice aged 7–9 weeks were used for all wound collection experiments. Prior to wounding, dorsal skin in the telogen phase of the hair cycle was shaved and treated with depilatory cream. Full-thickness 5 mm excisional wounds, extending through the panniculus carnosus, were created on the dorsal surface using a sterile biopsy punch. Wounds were bandaged with wound dressings consisting of non-adhesive gauze affixed with Tegaderm. Bandages were changed every 2 days until tissue harvest or 10 days post-injury. On days 4, 7, 15, and 30 post-wounding, skin samples encompassing the wound site and adjacent intact tissue were collected using a 10-mm biopsy punch for downstream single-cell sequencing, spatial transcriptomics, or histology.

### Single-nucleus isolation and sequencing from wounded mouse skin

Whole wound tissue sections (10mm) were flash-frozen in liquid nitrogen and stored until nuclei isolation. For tissue processing, samples were placed on glass petri dishes over dry ice and sectioned into rice-sized pieces using a chilled scalpel. Tissue fragments were transferred to pre-chilled cryovials and maintained on dry ice until digestion. Nuclei isolation was performed using the Singulator 100 (S2 Genomics) with NIC+ Isolation Cartridges using the extended nuclei isolation protocol. Both the instrument and cartridges were precooled at 4°C for 20 min prior to use. Tissue samples were loaded into NIC+ Isolation Cartridges with 75 μL of RNase inhibitor RNase Inhibitor V2 (Cat. #100-299-916, S2 Genomics) and Protector RNase Inhibitor (Cat. #3335402001, Roche)] and processed using the extended nuclei isolation protocol. Following digestion, samples were transferred to 15 mL conical tubes and centrifuged at 500 × g for 5 min at 4°C with low braking enabled. The nuclei pellet was subjected to debris removal using Nuclei Debris Removal Reagent diluted 1:5 with Nuclei Storage Reagent (Both S2 Genomics), followed by centrifugation at 500 × g for 15 min at 4°C. The resulting nuclei pellet was resuspended in 500 μL of nuclei resuspension buffer (1× PBS, 1% BSA, 0.2 U/μL RNase inhibitor (CAT. #M0314, NEB)) and filtered with a 40um filter (Cat. #431007040, pluriSelect).Nuclei were stained with DAPI (Cat. #D1306, Invitrogen) and counted using Millicellâ disposable hemocytometer (Cat. #MDH-2N1, Millipore) using a Nikon Ti2 fluorescence microscope to assess integrity and concentration prior to submission for sequencing. Cells were loaded onto the Chromium Controller (10X Genomics) for single-cell encapsulation and cDNA library generation using the Chromium Next GEM Single Cell 3ʹ Reagent Kits v3.1 (10× Genomics) or Chromium GEM-x Single Cell Gene Expression (3’) Kit. Libraries were sequenced on an Illumina NovaSeq6000 platform.

### Cellular Indexing of Transcriptomes and Epitopes by Sequencing (CITE-seq) isolation and sequencing

Whole wound (10 mm) tissues were harvested fresh and minced in 5 mL tubes using surgical scissors to a fine consistency. Samples were digested in buffer containing 0.1 mg/mL Liberase (Cat. #5401020001, Sigma) and 1 mg/mL DNase I (Cat. #10104159001, Sigma) in RPMI-1640 for 90 minutes at 37°C. Digested suspensions were filtered through a 70 μm filter (Cat. #352350, Corning), pelleted by centrifugation at 300 × g for 8 minutes at 4°C, washed once, and resuspended in 1 mL of cold FACS buffer (PBS w/ 1% BSA and 0.5 mM EDTA). Cells were stained with DAPI (Cat. #D1306, Invitrogen). Live, DAPI-negative cells were sorted into RPMI supplemented with 10% FBS. Sorted live cells were maintained on ice and blocked with anti-CD16/32 Fc blocking antibody (ThermoFisher Cat. #14-0161-82) at ∼1:100 dilution for 10 minutes at 4°C. Cells then were stained with reconstituted TotalSeq-B antibody cocktail (BioLegend, Cat. #199902) by adding 12.5–25 μL of antibody mix to a final staining volume of 50 μL (corresponding to a 1:4 to 1:2 dilution) and incubated for 30 minutes at 4°C. Following staining, 1 mL of RPMI with 10% FBS was added to each tube to wash unbound antibodies. Cells were pelleted, washed twice, and resuspended in 50 μL of RPMI with 10% FBS prior to sequencing. Samples were then loaded onto the Chromium Controller (10X Genomics) for single-cell encapsulation and cDNA library generation using the Chromium Next GEM Single Cell 3ʹ Reagent Kits v3.1 (10× Genomics).

### Spatial transcriptomic sequencing

Fresh mouse tissue wounds were prepared using formalin fixation and paraffin embedded (FFPE) and sectioned at 5-10 μm. Tissue sections were mounted on Apex Super Adhesive Slides (Cat. # 3800080E, Leica). Sections were submitted to the Columbia Human Immune Monitoring Core (HIMC). Spatial Transcriptomics data was generated using 10x Genomics Visium V2 CytAssist Spatial Gene Expression Mouse Transcriptome Assay (#1000445) for FFPE tissue as per user’s guide. The resulting libraries were sequenced on an Illumina NovaSeqX sequencer with pair-end 2×150 bp targeting 300 million reads per sample.

### Spatial transcriptomic processing

Manual image registration was performed in 10x Genomics Loupe Browser (v8, 10x Genomics) to align the H&E and CytAssist images prior to data processing. Data were processed using SpaceRanger (v3.1, 10x Genomics) for sequencing alignment with manual image alignment supplied. Aligned data were loaded into R using the *Load10X_Spatial function()* in Seurat (v4.3.0). Sequenced spots with fewer than 10 captured UMIs were filtered out, and the data were subsequently normalized and scaled using the *NormalizeData*(), *FindVariableFeatures(),* and *ScaleData()* functions in Seurat. For benchmarking clustering methods, we compared clusters generated by SpaceRanger’s graph clustering method (alignment_clusters), Seurat’s sketch clustering method (seurat_cluster.projected), and BANKSY^86^ spatially informed clustering Seurat sketch clustering was carried out using the SketchData() function with LeverageScore method on a subset of 50,000 spots, followed by principal component analysis (PCA) on the top 2,000 variable features. The top 30 principal components (PCs) were used for neighbor identification and uniform manifold approximation and projection (UMAP). BANKSY clustering was tested using multiple values of the spatial weighting parameter λ, which defines the influence of spatial coordinates on the clustering. A λ value of 0.5 was selected to equally weight cell type and spatial information in the resulting clusters, and a k_geom of 24 was used to capture all second-order spots. Quantification of clustering performance was assessed using normalized mutual information (NMI) relative to the cell type annotations generated from RCTD analysis^61^. For region-specific analyses, Loupe Browser was used to annotate regions of interest, and cluster coordinates were exported as .csv files for downstream analysis in R.

### Single cell dataset realignment and processing

Raw sequencing FASTQ and BAM files for publicly available single-cell RNA-seq runs were downloaded from the Gene Expression Omnibus (GEO) and Sequence Read Archive (SRA). When FASTQs were not available, BAM files were downloaded from SRA and converted to FASTQs using 10X Genomics *bamtofastq* function. Reads were aligned to the *Mus musculus* mm39 reference genome^30^ using the 10x Genomics Cloud Analysis pipeline (Cell Ranger v8.0.1). To assess potential discrepancies in gene representation, we compared the gene feature sets by extracting the row names of Seurat objects aligned to either mm10 or mm39 reference genomes. All data from the OWHA dataset were aligned exclusively to mm39.

The single-cell sequence data were subsequently mapped with CellRanger (version 7.2.0) to the GRCm39 mouse reference genome. Aligned datasets were loaded in R and merged into Seurat (v4)^87^ with metadata columns added for timepoint, modality, sex, and sample number. Only features detected in at least three cells were retained. Mitochondrial and ribosomal gene percentages were calculated using the *PercentageFeatureSet()* function, and the dataset was normalized using *NormalizeData()*. Quality control filtering was applied to retain cells with between 200-5,000 features, mitochondrial gene percentages <15%, and ribosomal gene percentages <50%. Potential doublets were identified using DoubletFinder^88^ and later confirmed and removed from subclusters. In total, 193,524 cells and 27,107 genes from 28 samples of unwounded and acutely wounded murine skin were retained for downstream analysis. The dataset was then normalized and scaled using *SCTransform()*^89^ while regressing out mitochondrial gene percentage. The top 3000 variable genes were used for principal component analysis (PCA). We next ran *FindNeighbors()* with 41 PCs, which was determined to be the minimum number of PCs required to capture 90% of the total variance. Clustering was performed using *FindClusters()* at a resolution of 1.5, and dimensionality reduction was performed with *RunUMAP()* using the same 41 PCs. Integration was performed using the reciprocal PCA (RPCA) method, maintaining the same dimensionality and resolution parameters for clustering and UMAP. Additional clusters enriched for ribosomal or mitochondrial genes were removed before rerunning *FindNeighbors()*, clustering, and UMAP to obtain the final OWHA dataset UMAP.

### Single cell integration testing

We benchmarked three integration approaches: Harmony (θ = 3)^90^, canonical correlation analysis (CCA)^91^, and reciprocal PCA (RPCA), using PCA without integration as a control. Integration performance was evaluated by calculating Local Inverse Simpson’s Index (LISI) scores for each method. We selected the integration strategy that most effectively minimized batch effects across modalities, datasets, and timepoints while preserving cell type diversity, as indicated by high and low LISI values, respectively.

### Single cell transcription factor scoring and stress scoring

To quantify transcription factor expression in each cell, we applied the *PercentFeatureSet()* function in Seurat using a curated list of transcription factors obtained from NCBI (**Table S5**). To calculate the stress score, we used a gene set defined by a past stress-dissociation sequencing study^92^. We then applied the *AddModuleScore()* function in Seurat to compute a weighted score for stress-associated genes in each cell.

During analysis, we observed that our single-nuclei modality exhibited lower stress scores compared to whole-cell modalities (single-cell and CITE-seq). To investigate these differences, we calculated the percent detection rate of each gene included in the stress score. To correct for single-nuclei biases, we selected transcription factors and stress-related genes that were detected in ≥10% of snRNA-seq nuclei to assemble a Stress Regulon Score. This score was then used in downstream analyses to determine stress responses across the OWHA atlas in cross modality comparisons.

### Single cell sex scoring

To deconvolve sex from mixed samples, we applied the *AddModuleScore()* function in Seurat using sex-specific gene sets for male and female mice^93–95^ (**Table S5**). Cells from sex-mixed samples were assigned a sex based on signature scores, with a threshold of >0.5 for classification as either male or female.

### Mouse skin-specific cell type annotation

After selection of the RPCA integration method, we performed k-nearest neighbor clustering at a resolution of 1.5 and identified 48 distinct clusters. To annotate these clusters in an unbiased and unsupervised manner, we benchmarked three single-cell annotation methods: ScType^36^, GPTcelltype^37^, and SingleR^96^. The development of our custom annotation framework, Skin-ScType, required a systematic and rigorous process. First, we performed a literature review to curate known marker genes for mouse skin cell types from multiple high-quality single-cell transcriptomics datasets^5–8,10,56,57,97–100^ (**Table S1**). Next, we evaluated gene expression by feature plot analysis to quantify inter-cluster expression across cell types. This was followed by iterative *in silico* validation within the OWHA dataset to refine these marker genes and to establish a threshold for marker gene specificity with minimal intercluster overlap. This iterative refinement enabled the annotation of both well-characterized major cell types and previously unclassified subpopulations, including keratinocyte subgroups, immune subsets, fibroblast subtypes, endothelial cells, and other specialized populations. Each cell type was subsequently mapped to marker genes with high discriminatory power at both the individual cluster and metacluster levels. Marker genes that passed specificity thresholds were assigned high-confidence labels and used for downstream analyses. For example, *Krt14* in combination with *Krt27* identified inner root sheath (IRS) keratinocyte subpopulations, while *Ptprc* together with *Csf1r* and *C3ar1* distinguished inflammatory macrophage subsets. These markers were validated across multiple wound single-cell databases to ensure specificity, minimal intercluster leakage, and reproducible expression patterns. The final Skin-ScType marker database was formatted according to *ScType* specifications to allow accurate marker-based cell type prediction in both R and Python environments. This curated resource provides a higher degree of precision than default annotation databases and enables robust annotation of mouse skin single-cell data.

### Subclustering and annotation of metaclusters populations

Each metacluster group was subset and integrated using Harmony (θ = 3). Harmony integration was determined to be optimal based on LISI scoring, which indicated improved preservation of biological variation and reduced batch effects. Dimensionality reduction was performed using the minimum number of PCs which capture 90% of the total variance across the dataset, followed by neighbor graph construction with *FindNeighbors(*), clustering with *FindClusters()* at a resolution of 1, and visualization with *RunUMAP()*. Clusters were then annotated using canonical marker genes in combination with the *FindMarkers()* function in Seurat. Clusters exhibiting ambiguous cell type identity, defined by the co-expression of markers from two distinct lineages (e.g., immune and keratinocyte markers or fibroblast and immune markers), were classified as putative doublets and removed. In addition, cells with extreme UMI counts were excluded. Specifically, cells with UMI counts below the 10th percentile (849 UMIs) or above the 90th percentile (11,410 UMIs) were removed prior to final subcluster analysis. Each refined metacluster dataset was then re-integrated to yield a high-quality set of cells for subclustering. These subcluster-level UMAP embeddings were used for subsequent downstream analyses.

Given the transcriptional heterogeneity of fibroblasts, we applied a fibroblast-specific annotation strategy adapted from prior work^57^. This approach leverages curated gene modules and Seurat’s *AddModuleScore()* function to assign fibroblast subtypes, including myofibroblast, reticular, papillary, and fascia-associated populations.

### Dendrogram Construction

Hierarchical relationships among cell clusters were visualized using dendrograms constructed with the *BuildClusterTree()* function in Seurat. Dendrograms were generated based on 44 principal components, which together captured 90% of the total transcriptional variance in the dataset.

### Differential expression analysis and Gene Ontology testing

Differential expression analysis was performed using the DElegate package^101^ (v1.2.1). Counts were grouped by sequencing batch and pseudobulked to enable differential expression comparisons across timepoints. Genes were considered significantly upregulated if they exhibited a log₂ fold change > 1 and an adjusted *P* value ≤ 0.01.

Gene ontology enrichment was performed using the clusterProfiler package^102^ (v4.0.5). Gene symbols were first converted to Entrez IDs using the *bitr()* function. Data were then grouped, and the *compareCluster()* function was applied with the org.Mm.eg.db annotation database, using a *P* value cutoff of 0.05. Unless otherwise specified, enrichment was restricted to the Biological Process (BP) ontology. Results were exported and visualized in R.

### Differential abundance testing

We tested for differential cell-state abundances of subpopulations across wound healing using the MiloR package (v1.6.0)^39^. A K-nearest neighbors (KNN) graph was built using the RPCA integrated slot from the adjacency matrix of the processed Seurat object with the parameters: k = 40 and d=30. Cells were assigned to the neighborhoods based on the KNN graph using the ‘makeNhoods’ function (prop=0.1). Differential neighborhood abundance testing was performed using a generalized linear model (GLM). Differentially abundant cell neighborhoods with SpatialFDR ≤ 0.1 were plotted using the *plotNhoodGraphDA()* function.

### CellChat cell-cell communication analysis

Cell–cell communication analysis was performed using CellChat (v2.2.0)^54^. Seurat objects were imported into CellChat, and the standard CellChat pipeline was applied. First, overexpressed genes and potential ligand–receptor interactions were identified using *identifyOverExpressedGenes()* and *identifyOverExpressedInteractions(),* respectively. Data were then smoothed using the smoothData() function, and subsequent analyses were performed following the CellChat vignette. CellChat signaling strengths were determined using the compute *CommProb()* function. Circle plots were then generated with *netVisual_circle(),* while differential interaction analysis between conditions or timepoints was performed using the *netVisual_diffInteraction()* and *rankNet()* functions. Dot plots of predicted ligands were produced using *netVisual_bubble()*.

### Pseudotime monocle 3 analysis

Pseudotime analysis was performed using Monocle3 (v 1.3.7)^103^. IFE keratinocytes from snRNAseq batches were isolated and subsequently subset by timepoint, with groups defined as Unwounded (UW) and Wounded (D4PW/D7PW). The root of the trajectory was set to the cell group with the highest *Krt14* expression. Differential genes along each pseudotime trajectory of interest were identified using the *graph_test()* function, filtered for a q-value < 0.05, and arranged by Moran’s I value.

### Spatial transcriptomic robust cell type deconvolution (RCTD) annotation

RCTD was performed following the published guidelines and Seurat vignette for spatial deconvolution (https://satijalab.org/seurat/articles/spatial_vignette.html). Briefly, clusters were determined using Seurat’s sketch clustering method with return.model = TRUE in the *RunUMAP()* function. The annotated OWHA dataset was then used to generate the RCTD reference object, which was supplied to the *create.RCTD* function with a minimum UMI threshold of 10 and a minimum cell instance threshold of 20. RCTD was run in doublet mode to identify spatial barcodes representing mixtures of multiple cell types. The method was implemented as previously described in Cable et al. (*Nat. Biotechnol.* 40, 1025–1034, 2022).

### Spatial transcriptomic major anatomical site annotation

Tissue sections were annotated for major anatomical sites, including the epidermis, dermis, and adipose layers, as well as cell groups such as fascia, hair follicle, muscle, and endothelial cells. Annotation was performed using both hematoxylin and eosin (H&E) staining and cellular transcriptomic markers (*Krt14, Krt15, Krt79, Krt17, Krt77, Pecam1, Acta2, Adipoq1, Mylpf, Col1a2*). Wound sites were identified based on the loss of adipose tissue, muscle, and hair follicle structures within the tissue sections.

### Spatial transcriptomic Basal IV and proliferative endothelial scoring

Spatial transcriptomic data was scored using the top 10 marker genes from the OWHA data identified by the Seurat *FindMarkers()* function. Signature scores were calculated using the *AddModuleScore()* function for both Basal IV and Proliferative Endothelial cell populations. Spots were classified as Basal IV or Proliferative Endothelial if they had a signature score >0.2 and an average expression level >1 for *Krt14* (epithelial) or *Pecam1* (endothelial), respectively.

### Variance-based spatial localization test

Quantification of cluster and cell type localization in Visium HD data was carried out using a variance-based localization test. First, PC analysis was used on the tissue coordinates to generate component vectors associated with tissue length and depth axes. Annotated spots were then projected onto this 1D axis to yield distributions. To determine significance of localization for each cluster, we computed the variance of its normalized projections along the normalized axis of tissue length (values scaled between 0 and 1). To assess whether the observed variance differed significantly from the null expectation under a uniform spatial distribution, we used two complementary approaches depending on cluster size. For large clusters (≥ 500 cells by default): an analytic chi-square test was applied, comparing the observed variance to the theoretical variance of a Uniform [0,1] distribution (variance = 1/12). Test statistics were converted to two-sided p-values. For small clusters (< 500 cells): A bootstrap approach was used, repeatedly sampling the same number of points from a Uniform [0,1] distribution to generate a null distribution of variances. The empirical p-value was defined as the proportion of bootstrap variances at least as extreme as the observed value. To provide a standardized effect size across clusters, we also calculated a z-score: the deviation of the observed variance from the null variance (generated from a hypothetical null cluster of uniform distribution), scaled by the standard deviation of the null distribution. This z-score indicates whether a cluster is more localized (low variance, negative z) or more dispersed (high variance, positive z) than expected under spatial uniformity. Multiple testing correction was applied to p-values using the Benjamini–Hochberg procedure, and clusters were annotated with significance levels (p < 0.05 *, p < 0.01 **, p < 0.001 ***) where appropriate.

### Fluorescent *in situ* hybridization (FISH)

Fluorescent *in situ* hybridization (FISH) was performed using RNAscope technology (Advanced Cell Diagnostics, ACD). Probes for *Sema3c* and *Sox6* (CAT# 441441, RNAscope Probe – Mm-Sema3c-C3; CAT# 472061-C2, RNAscope Probe RNAscope Probe – Mm-Sox6-C2) were obtained from ACD. Mouse wound sections were prepared according to the manufacturer’s instructions. Slides were fixed with 4% paraformaldehyde (PFA) for 1 hour at 4 °C and then dehydrated through an ethanol series (50%, 70%, and 100%). Following dehydration, slides were treated with Protease IV (CAT# 322336, ACD) for 30 minutes at room temperature. Hybridization was carried out by incubating the sections with probes for 2 hours at 40 °C using the HybEZ™ II Hybridization System and the RNAscope® Multiplex Fluorescent Reagent Kit v2 (ACD). Signal amplification was performed according to the manufacturer’s protocol, and images were captured using a Nikon Ti2 microscope.

### Cross-species atlas generation

Orthologous gene mapping was performed to harmonize human and mouse single-cell datasets. A custom ortholog table was generated containing human–mouse gene symbol conversions using nichenetR’s^104^ (v 2.2.0) *convert_mouse_to_human_symbols*() function. For mouse data, count matrices were extracted from Seurat objects and gene symbols were mapped to their corresponding human orthologs. Genes without human counterparts were removed, and duplicated gene entries were collapsed. The human wound healing single-cell dataset was downloaded from GEO (GSE241132). Count matrices were extracted and mapped to mouse orthologs. Rows without a mouse ortholog were removed, and a parallel Seurat object with mouse gene symbols was generated. Both the human and mouse Seurat objects were then subset to include only intersecting orthologous genes. To generate a joint cross-species atlas, Seurat objects were labeled by species (“Human” or “Mouse”), assays were restricted to RNA, and SCT assays were removed for consistency. The two objects were then merged into a single Seurat object. After merging, data were normalized and variance-stabilized using SCTransform() while regressing out mitochondrial percentage. Dimensionality reduction was performed using PCA, followed by nearest-neighbor graph construction, clustering (resolution = 0.5), and UMAP embedding. For cross-species integration, Harmony (θ = 3) was applied using SCT-normalized values with the PCA reduction as input. The integrated dataset was reclustered based on the Harmony embeddings, and a UMAP was generated for visualization of the cross-species atlas.

Human wound healing spatial transcriptomics data were also downloaded from GEO (accession number GSE241124) and processed using Seurat. Raw data were imported as Seurat objects, and the default assay was set to RNA for downstream analysis. To examine basal keratinocyte spatial dynamics, we focused on the Basal IV keratinocyte population. Module scores were calculated using *AddModuleScore()* with the same gene sets identified in the mouse data sets. Spots with *KRT5* expression > 2 were classified as epidermis-enriched while spots with PECAM1 expression > 2 were classified as endothelial-enriched and used for downstream analysis and graphing to classify both keratinocyte and endothelial spatial areas.

### Recombinant Sema3C treatment

Functional validation of the SEMA3C axis on chronic wound pathophysiology was conducted in 18-week-old *Lepr^db^*/J diabetic mice (Jackson Laboratory). For wounding, dorsal skin was shaved and treated with depilatory cream. Full-thickness 1-cm full-thickness excisional wounds were generated on the dorsal skin to create a large diabetic ulcer. Mice received 25 µL of recombinant mouse (mSema3c; Bio-Techne, Catalog #: 1728-S3) or human (hSEMA3C; Abcam, Catalog #: ab153780) protein at 20 µg/mL. Treatments were applied topically every other day for four days (D0, D2, and D4). Wound closure rates were monitored digitally using a TOMLOV DM10 10.1” Digital Microscope at D0, D2, D4, D7, D9, and D14PW to compare the recovery speed of treated *Lepr^db^*/J mice against PBS vehicle controls and wild-type (WT) mice. Subcutis revascularization and periwound angiogenic sprouting were assessed via digital imaging at D14PW to evaluate the proportion of revascularization coverage in the wound bed.

## DECLARATION OF INTERESTS

All other authors declare that they have no competing interest.

## AUTHOR CONTRIBUTIONS

Y.W conceptualized and designed the study. J.C and S.V performed the experiments. J.C, S.V, D.A conducted the computational analyses. D.A, SH, S.V and Y.W conducted mouse studies. J.C, S.V, and Y.W. analyzed and interpreted the data. J.C, S.V, and Y.W. wrote the manuscript.

## ACKNOWLEDGEMENTS

We would like to thank Erin Bush, the Single Cell Core, and the Columbia Genome Center core facility. Parin Shah, and the Human Immune Monitoring Core. The Molecular Pathology Shared Resource macromolecule and histology services. Ronan Chaligne and the Single-cell Analysis Innovation Lab at Sloan Kettering Institute for assistance with technical troubleshooting, helpful comments, and consultation on this project. This work was supported by grants R00GM140262, K99GM140262; Columbia Data Science Institute Seeds Funds; CUIMC Healthy Aging Initiative Award. J.C was supported by NIH 5T32HL120826.

## TABLES

**Table 1: OWHA top fine cell type markers**

**Table 2: Functional hierarchy of wound-activated cell states organized by metacluster**

## SUPPLEMENTAL TABLES

**Supplemental Table 1: Summary of publicly available scRNA-seq datasets of mouse skin wounds**

**Supplemental Table 2: Summary of Single Cell Sequencing Batches**

**Supplemental Table 3: Skin ScType database**

**Supplemental Table 4: COWA metacluster cluster correspondence**

**Supplemental Table 5: Genes used for scoring stress, sex, and transcription factors**

## SUPPLEMENTAL FIGURES

**Supplemental Figure 1.**
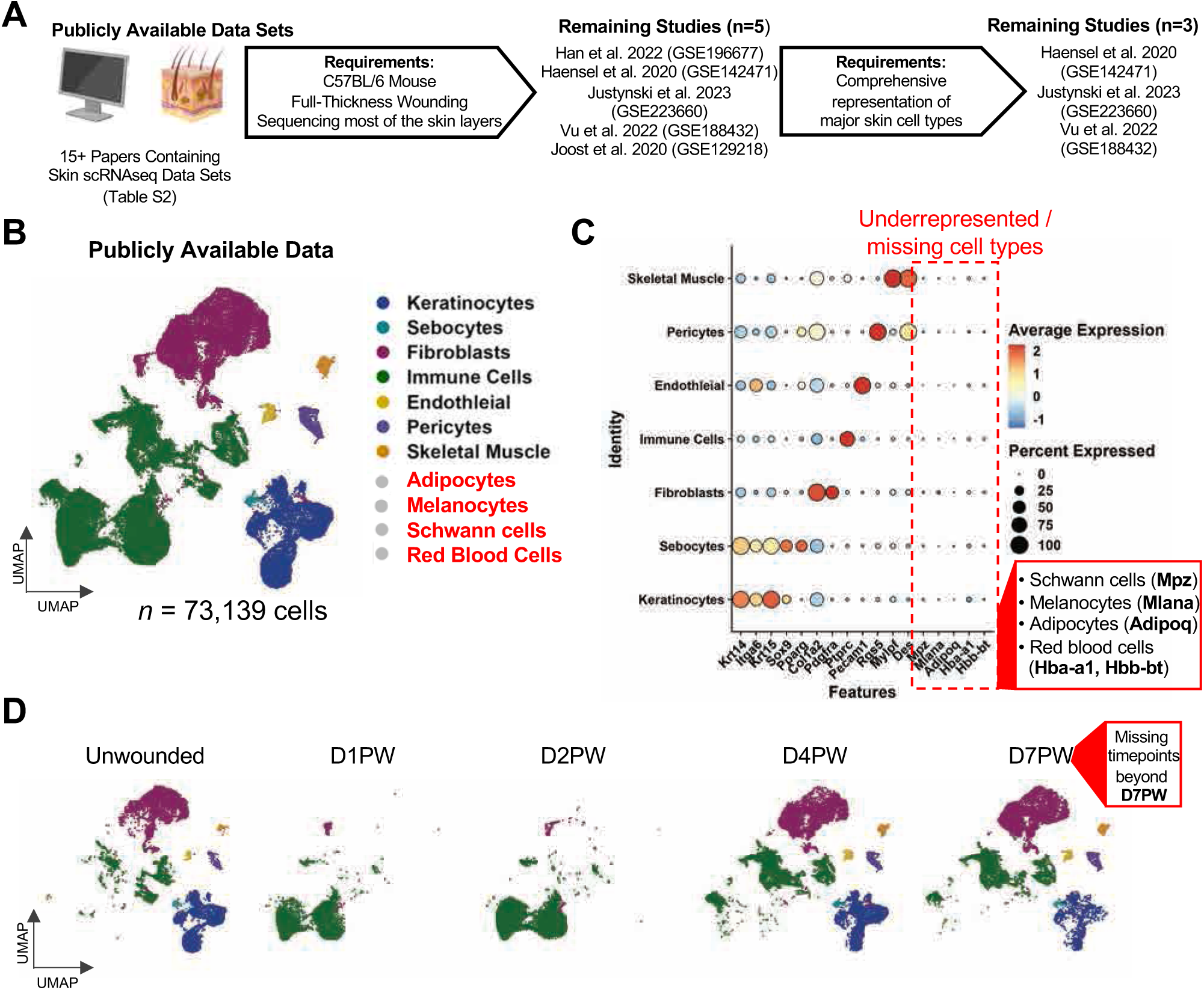
Identifying knowledge gaps in publicly available skin wound healing datasets. **(A)** Schematic illustrating the inclusion and exclusion criteria used to select rigorous publicly available scRNAseq wound datasets for our analysis. **(B)** Uniform Manifold Approximation and Projection (UMAP) of the three publicly available datasets that met the inclusion criteria. Data was integrated by Harmony and colored by major metaclusters representing major cell type classes. Clusters highlighted in red indicate critically underrepresented or missing cell types. *n* = 12. **(C)** Dot plot heatmap displaying marker gene expression for major metaclusters identified across the integrated publicly available datasets. Boxed genes indicate cell type markers underrepresented or missing from integrated public datasets, including melanocytes (*Mlana*), adipocytes (*Adipoq*), and red blood cells (*Hba-a1, Hbb-bt*). **(D)** UMAP projection of the integrated publicly available datasets, colored by major cell clusters and split by timepoint to visualize temporal dynamics in skin wound healing. Notably missing are late stage wound healing time points beyond D7PW.

**Supplemental Figure 2.**
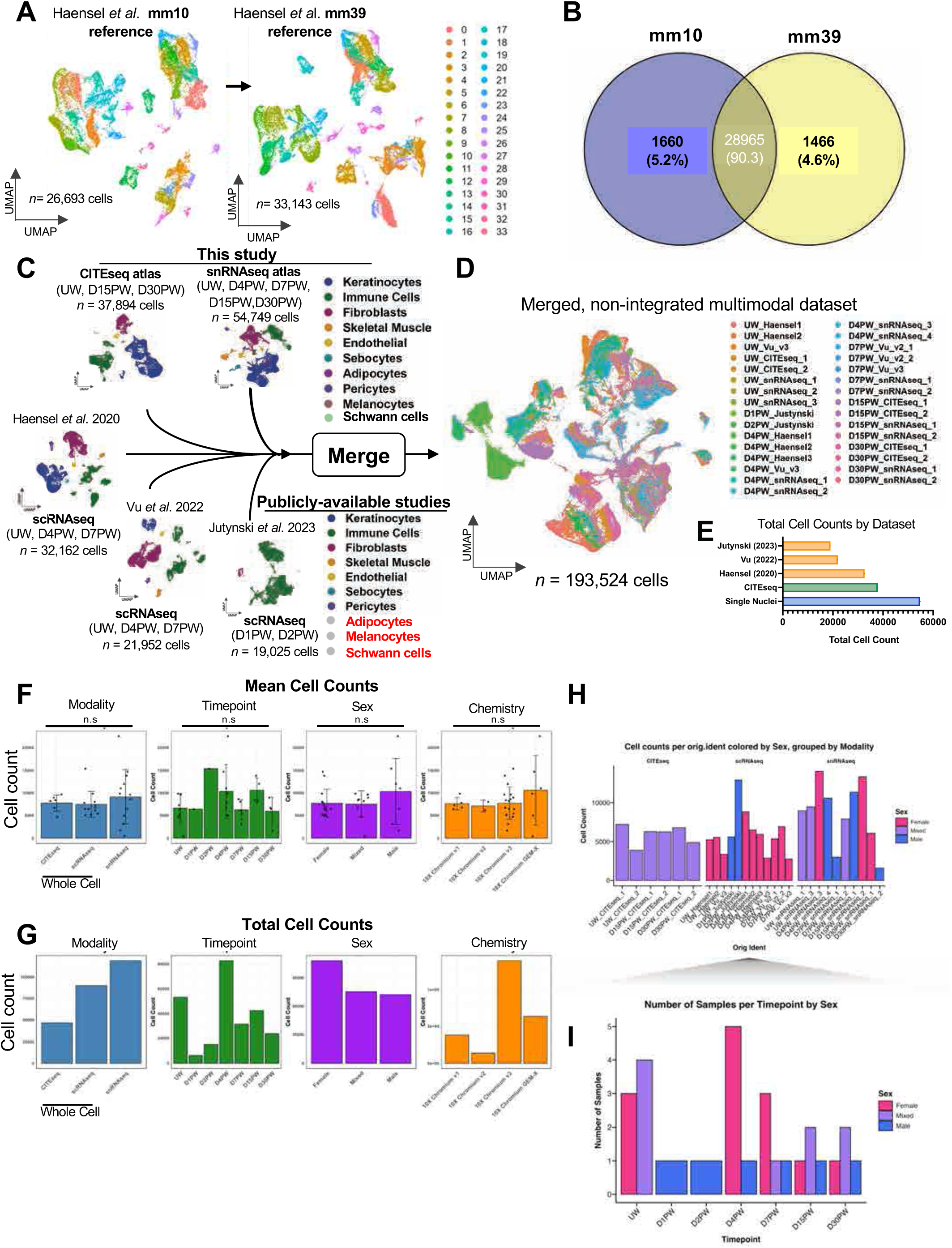
Multimodal whole-cell sequencing and snRNA-seq datasets are matched for cell number and sex across mouse skin wound samples. **(A)** Uniform Manifold Approximation and Projection (UMAP) plots of data from Haensel *et al*. 2020 (GSE142471) before (left) and after (right) realignment from the mm10 (*n*=26,693 cells) to the mm39 (n= 33,143 cells) reference genome. This approach was applied to all samples in the study. **(B)** Venn diagram depicting number and proportion of genes uniquely found in the mm10 (left) or mm39 (right) reference genome, or shared by both, for all datasets included in the study. **(C)** UMAP projections of source data from different studies merged to generate a multimodal dataset of mouse skin wound healing. Populations named in red indicate missing or underrepresented cell types from publicly-available studies. **(D)** Unintegrated UMAP grouped by sample after data merging, prior to integration. *n* **=** 28 batches. **(E)** Quantification of total cell counts for the final 3 external public datasets curated (Jutynski *et al*. 2023, Vu *et al*. 2022, Haensel *et al*. 2020), along with data collected in this study using CITE-seq and snRNA-seq. **(F-G)** Average (top) and total (bottom) quantification of cell counts in sequenced batches after QC filtering. Left to right, cell counts by modalities, timepoints, sex, and sequencing chemistries. Bar graphs represent the mean; error bars represent standard deviation (s.d). Significances were determined using a one-way ANOVA plus Tukey’s MC test (*: p < 0.05, n.s: no significance). **(H)** Bar plot of total cells counts sequenced across batches and modalities. Blue indicates male samples, pink indicates female samples, and purple indicates sex-mixed samples. **(I)** Bar Plot of total number of batches sequences for each wound healing timepoint. Blue indicates male samples, pink indicates female samples, and purple indicates sex-mixed samples.

**Supplemental Figure 3.**
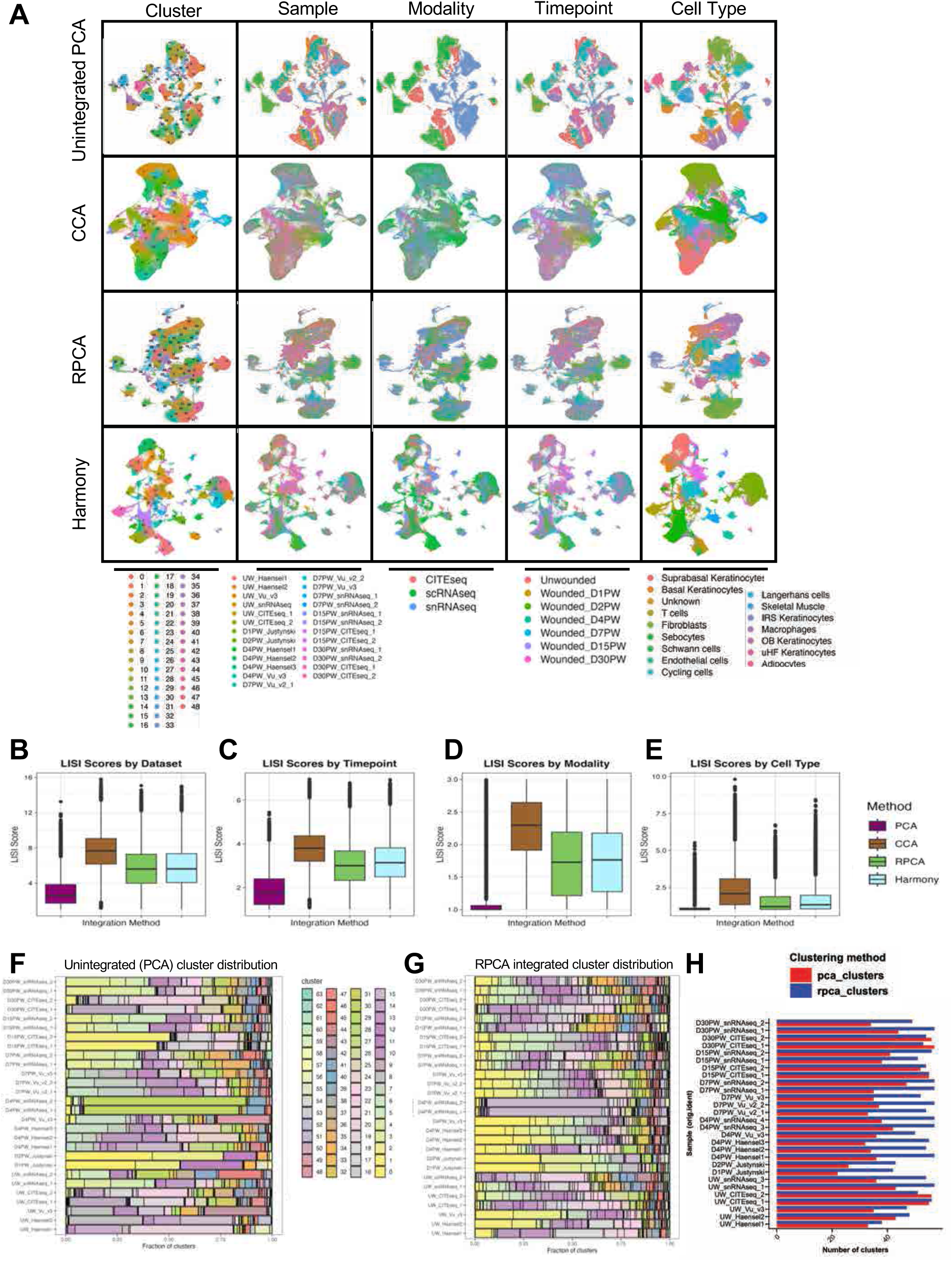
Integration benchmarking of multimodal wound sequencing data. **(A)** UMAP projections of cells processed using unintegrated PCA and three integration strategies (CCA integration, and RPCA integration). UMAPs are colored by cluster, sample, modality, and timepoint. Legends corresponding to each metadata category are shown beneath the respective panels. (*n* = 193,524 cells). **(B–E)** Box plots of Local Inverse Simpson’s Index (LISI) scores generated for each integration method (unintegrated PCA, CCA, RPCA). LISI was calculated after labeling cells by **(B)** dataset **(C)** timepoint, **(D)** modality, and **(E)** cell type. Box plots display the median, interquartile range (IQR), whiskers representing 1.5 × IQR. **(F-G)** Stacked bar plots quntifying the distribution of individual samples across **(F)** unintegrated PCA clusters and **(G)** RPCA-integrated clusters. RPCA demonstrates more even representation of samples across clusters, indicating improved batch harmonization relative to PCA. **(H)** Bar plot comparing the number of clusters detected within each sequencing batch before integration (red) and after RPCA integration (blue). RPCA substantially reduces batch-specific clustering artifacts.

**Supplemental Figure 4.**
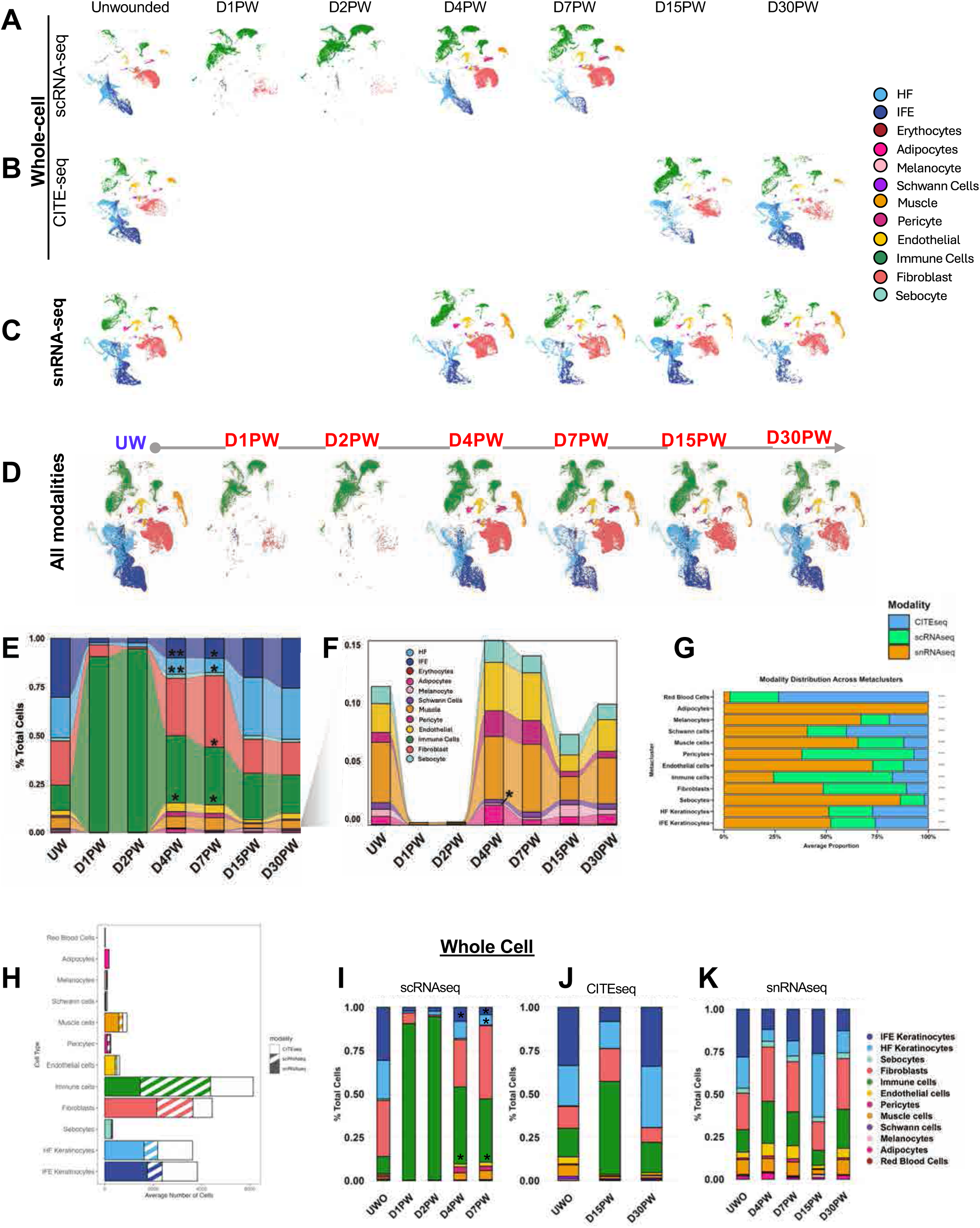
Characterization of modality differences between wound healing atlas. **(A-C)** RPCA-integrated UMAP projection of scRNAseq (A), CITEseq (B), and snRNAseq (C) datasets split by timepoint. Colored by major metaclusters labeled in wound healing atlas. **(D)** UMAP projection of the OWHA atlas separated by timepoint, colored by major metaclusters. n =193,524 cells. **(E)** Stacked bar plots showing the proportional representation of metaclusters across timepoints. Significance was assessed using the Wilcoxon rank-sum test relative to UW (*p* < 0.05 = *, *p* < 0.01 = **). **(F)** Stacked bar plots showing the proportional representation of rare cell types with representation <0.15% for visualization. Significance was assessed using the Wilcoxon rank-sum test relative to UW (*p* < 0.05 = *, *p* < 0.01 = **). **(G)** Stacked bar plot showing proportion of cell type metacluster captured by each modality across all timepoints. Significance was assessed using a Chi-Square test (*p* < 0.05 = *, *p* < 0.01 = ***). **(H)** Bar Plot showing average number of cells from each modality [scRNAseq (stripes), CITEseq (blank), snRNAseq (solid)] for each of the major cell Metaclusters. **(I-J)** Stacked bar plots depicting the proportions of major metaclusters over time separated by modality.

**Supplemental Figure 5.**
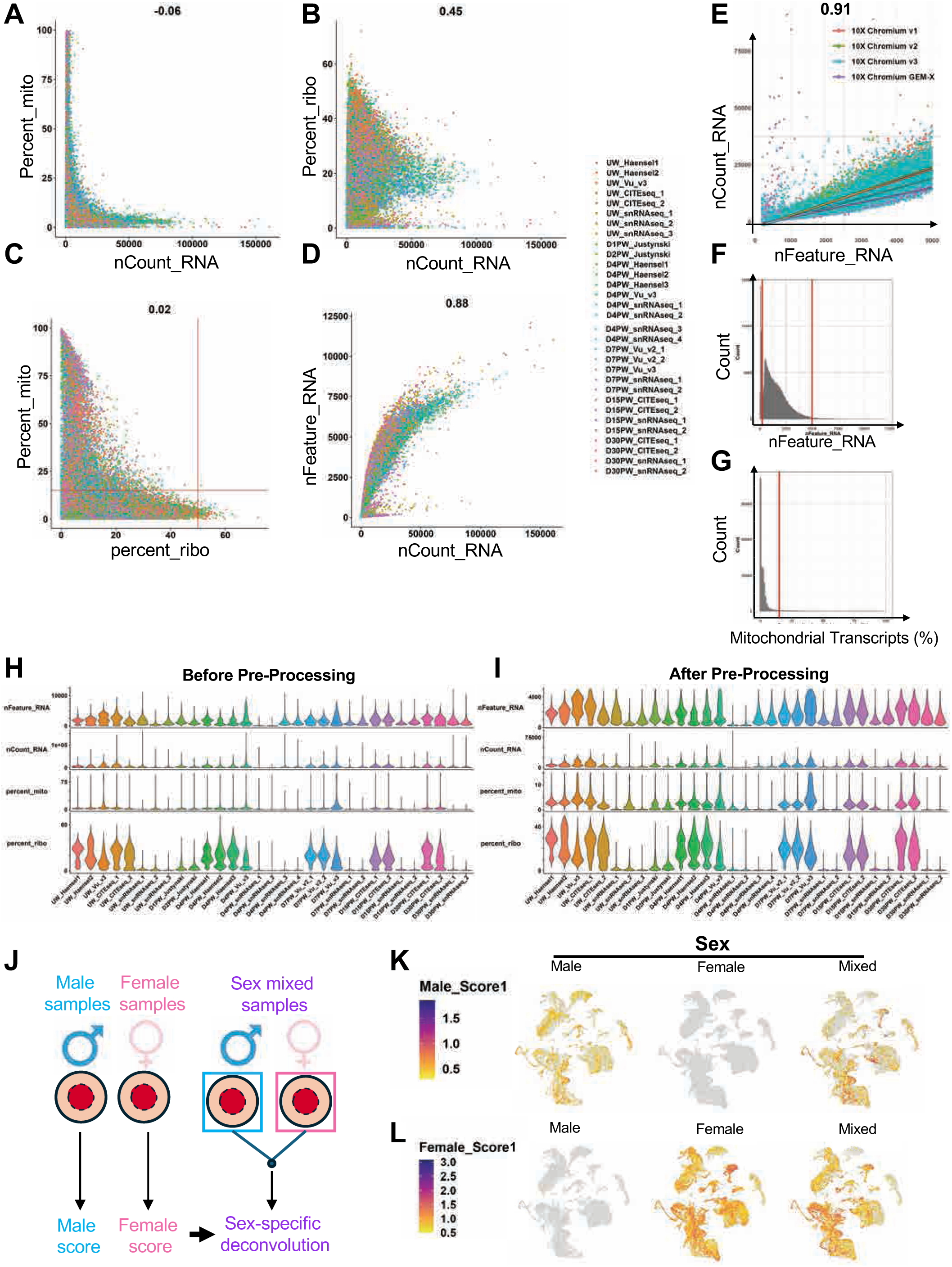
Sample benchmarking across sex and transcript capture. **(A-D)** FeatureScatter plot showing the Pearson correlation between percentage of mitochondrial transcripts (percent_mito) and total RNA counts (nCount_RNA) per cell. (A) percentage of ribosomal transcripts (percent_ribo) and nCount_RNA per cell (B), percent_ribo and percent_mito per cell (C), and unique genes detected (nFeature_RNA) and nCount_RNA (D). Points are colored by sequencing batch (n = 28 sequencing runs). **(E)** FeatureScatter plot showing the Pearson correlation between nFeature_RNA and nCount_RNA, colored by sequencing chemistry. R and R² values are shown. All 28 sequencing batches are included. **(F)** Histogram of nFeature_RNA values per cell across all sequencing batches. Red vertical lines indicate the filtering thresholds applied during preprocessing. **(G)** Histogram of percent_mito values per cell. Red vertical lines indicate cutoffs used for filtering during preprocessing. **(H-I)** Violin plots showing QC metrics **(H)** before and **(I)** after QC filtering and preprocessing for each sequencing batch, including nFeature_RNA, nCount_RNA, percent_mito, and percent_ribo. **(J)** Schematic illustrating the development of sex-based transcriptomics signatures. Samples from each sex were sequenced individually, and key genes exclusively expressed by each sex were identified. These sex-specific genes were then used to assign individual cells to their respective sex within sex-mixed samples, relative to groundtruth controls. **(K-L)** Feature expression plots showing male (K) and female (L) scoring of aggregated gene expression of sex-specific genes (Table S3), split by sex.

**Supplemental Figure 6.**
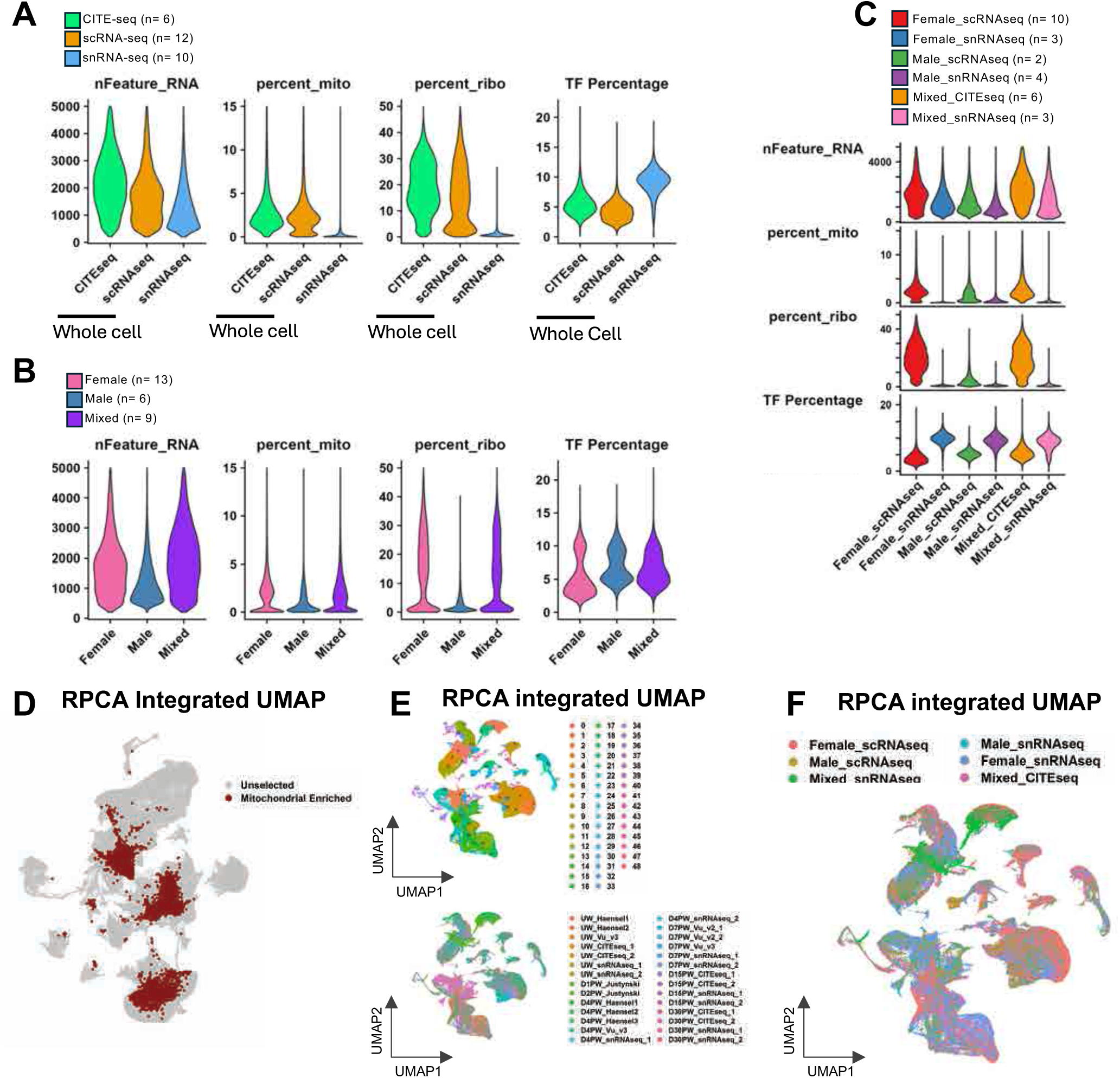
Quality control assessment of integrated wound data by sex and sequencing modality. (**A**) Violin plots of quality control (QC) metrics quantifying number of unique genes (nFeature_RNA), percentage of mitochondrial genes, percentage of ribosomal genes, percentage of transcription factor (TF) genes detected in whole-cell and snRNA-seq modalities. CITE-seq, *n* =6; scRNA-seq, *n* = 12; snRNA-seq, *n* = 10. (**B**) Violin plots of QC metrics (nFeature_RNA, percent_mito, percent_ribo, TF percentage) in samples from male or female batches, as well as batches containing a mix of both female and male-derived cells. Female, *n* = 13; Male *n =* 6; Mixed *n = 9*. (**C**) Violin plots of QC metrics (nFeature_RNA, percent_mito, percent_ribo, TF percentage) in samples from male or female batches, as well as batches containing a mix of both female and male-derived cells separated also by modality as well Female *n* = 13, Male *n =* 6, Mixed *n = 9*. (**D**) UMAP projection of RPCA integrated data highlighting cell populations highly enriched for mitochondrial transcripts (>30%, red). These populations were removed prior to downstream analysis. (**E**) UMAP projection of the RPCA integrated skin wound healing atlas after subsetting, colored by integrated clusters (top) and sequencing batch (bottom). (**F**) UMAP projection of the RPCA integrated skin wound healing atlas colored by sex and modality combined.

**Supplemental Figure 7.**
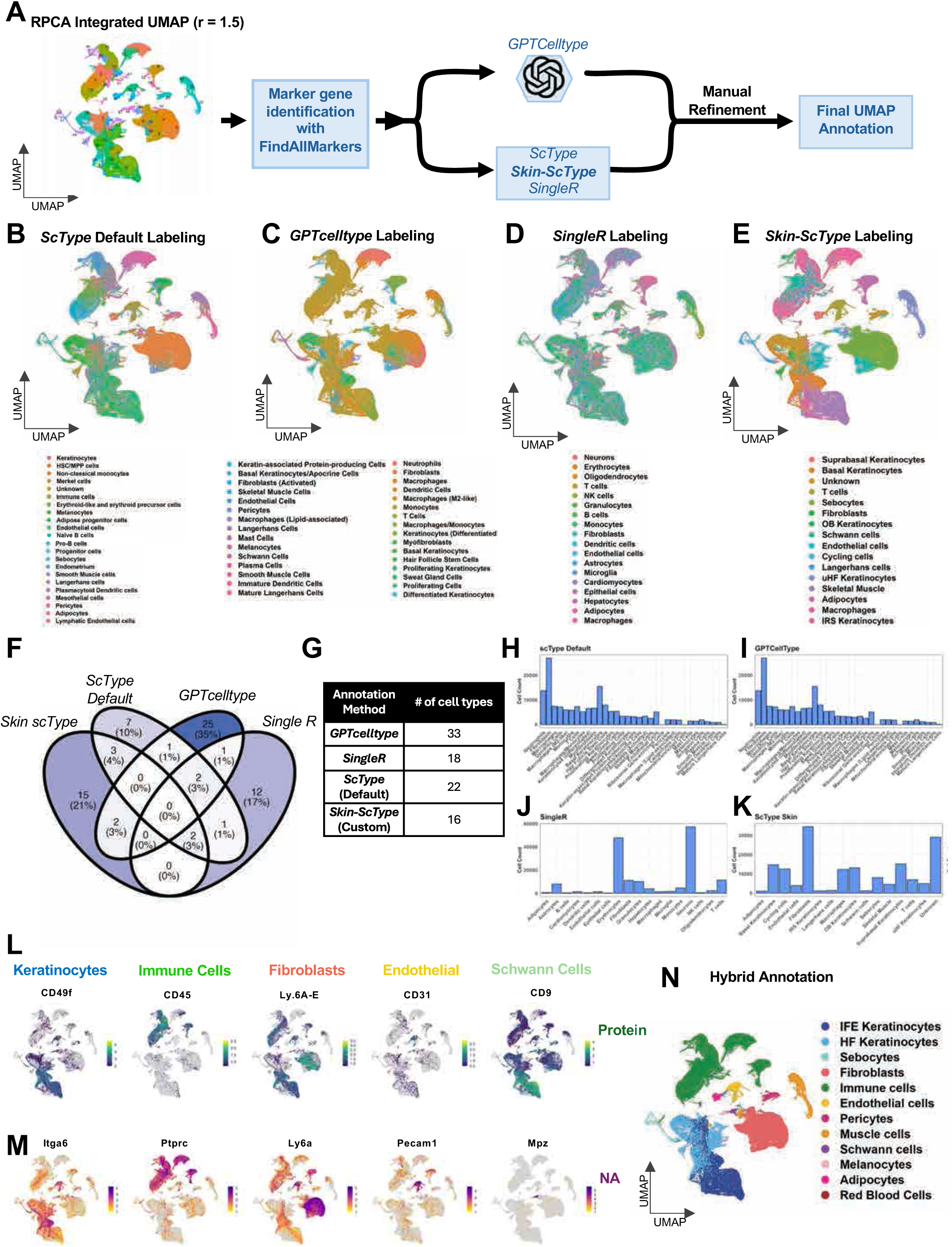
Development of mouse skin-specific cell type annotation methods. (**A**) Schematic of workflow pipeline in labeling RPCA integrated clusters using two automated strategies (GPTCelltype, ScType) to achieve our final annotation with a manual refinement. (**B**) UMAP projection of the RPCA-integrated dataset, colored by annotation generated using the default ScType database. (**C**) UMAP projection of the RPCA-integrated dataset colored by annotation generated using GPTcelltype. (**D**) UMAP projection of the RPCA-integrated dataset, colored by annotation generated using singleR (**E**) UMAP projection of the RPCA-integrated dataset, colored by annotation generated using a custom skin-specific ScType database. (**F**) Venn diagram showing overlapping annotation names among all four annotation strategies. (**G**) Table showing annotation methods and resulting number of cell types identified. **(H-K)** Bar plots showing annotation names and resulting cell counts identified by each annotation method: Default ScType (**H**), GPTCellType (**I**), SingleR (**J**), and Skin-ScType (**K**). **(L)** Feature expression plots of the RPCA-integrated dataset highlighting expression of CITE-seq proteins used to verify major meta-cluster populations (Keratinocytes: CD49f, Immune cells: CD45, Fibroblasts: Ly6A/E (Sca1), Endothelial cells: CD31 (Pecam1), Schwann cells: CD9). **(M)** UMAP projection of the RPCA integrated colored by annotated metaclusters determined through combining both automated cell labeling and manual refinement. **(N**) RPCA Integrated UMAP colored by hybrid annotation derived from Skin-ScType (n = 193,524 cells).

**Supplemental Figure 8:**
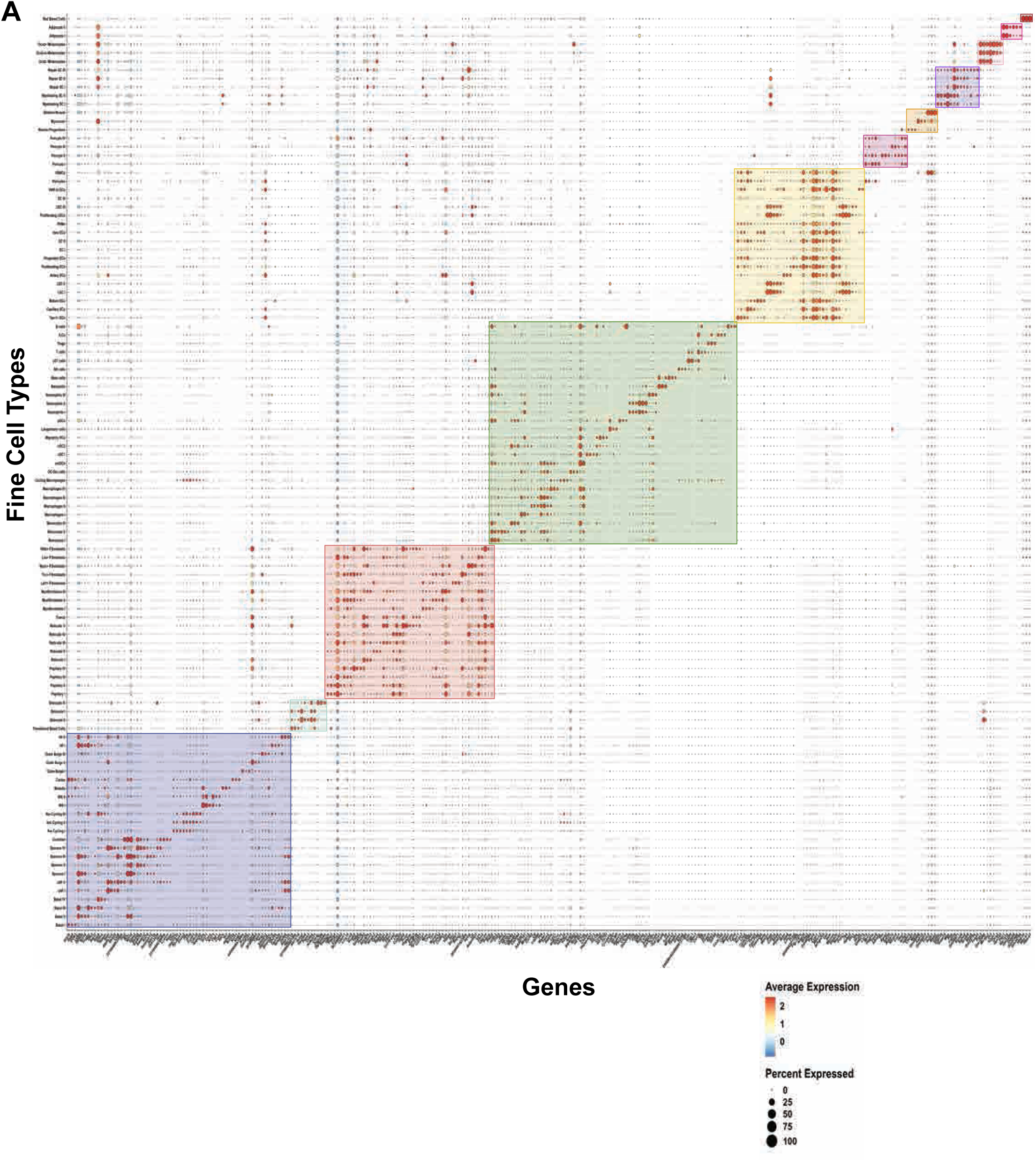
OWHA Sub clustering marker expression. **(A)** Dot plot of gene expression of top marker genes for each subcluster identified in OWHA. Size of dots depict percent of subcluster that is expressing marker genes. Genes are grouped together by major meta clusters.

**Supplemental Figure 9:**
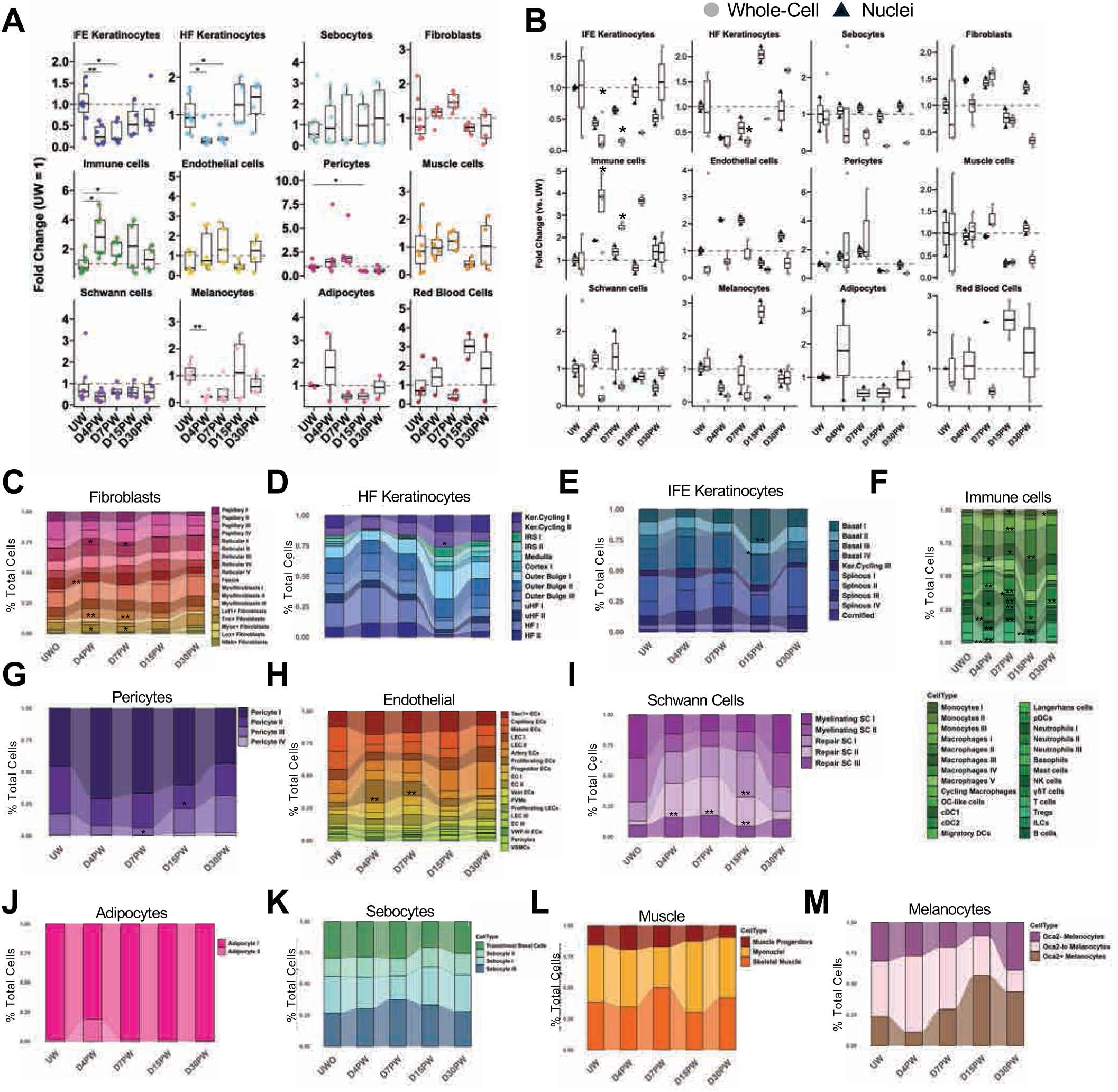
Temporal dynamics of OWHA subclusters throughout healing response. **(A)** Box and whisker plots quantifying fold-change in proportion of major metaclusters across all modalities. Each dot represents an individual sequencing run: UW = 7, D4PW = 6, D7PW = 5, D15PW = 4, D30PW = 4. **(B)** Box and whisker plots showing fold-change in the proportion of major metaclusters separated by modality: whole-cell (grey circles) and single-nucleus (black triangles). (A and B) Significance was assessed using the Wilcoxon rank-sum test relative to UW (*p* < 0.05 = *, *p* < 0.01 = **). UW = 7, D4PW = 6, D7PW = 5, D15PW = 4, D30PW = 4. **(C-M)** Stacked bar plots of subcluster proportions over healing time in **(C)** Fibroblasts, **(D)** HF Keratinocytes, **(E)** IFE Keratinocytes, **(F)** Immune Cells, **(G)** Pericytes, **(H)** Endothelial, **(I)** Schwann Cells, **(J)** Adipocytes, **(K)** Sebocytes, **(L)** Muscle, **(M)** Melanocytes.

**Supplemental Figure 10:**
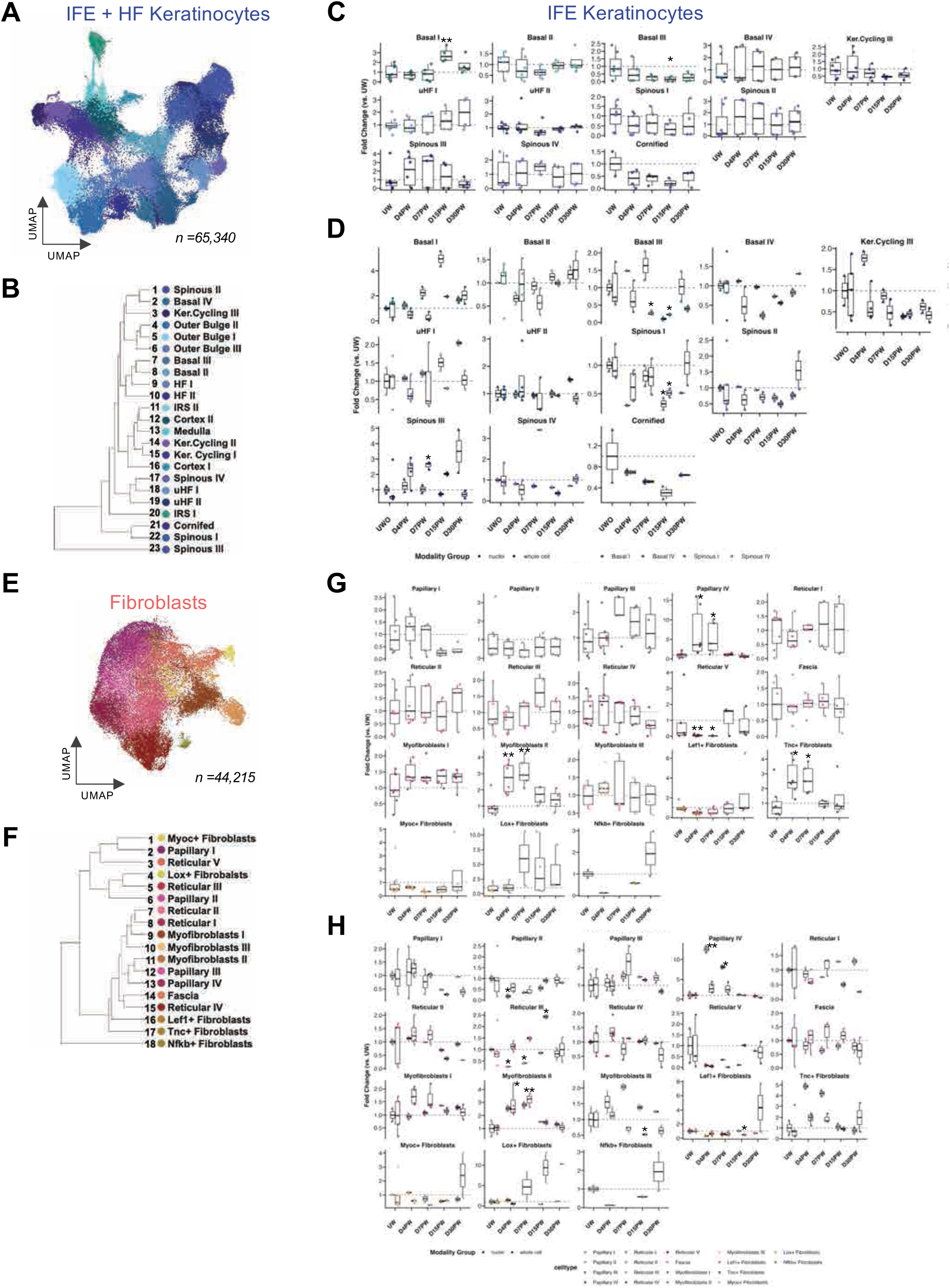
Keratinocyte and fibroblast subsets are heterogeneous throughout wound healing. (**A**) UMAP embedding of IFE and HF keratinocyte subclustering from all timepoints, colored by fine cell type annotation. n = 65,340 cells. (**B**) Dendrogram depicting relationships between keratinocyte subsets, generated from PCA. (**C**) Boxplots depicting fold change in IFE keratinocyte subset proportions relative to UW across all modalities. (**D**) Boxplots depicting fold change in IFE keratinocyte subset proportions relative to UW, separated by whole cell (circles) and single nuclei (triangles) modalities. (**E**) UMAP embedding of Fibroblast subclustering from all timepoints, colored by fine cell type annotation. n = 44,215 cells. (**F**) Dendrogram depicting relationships between fibroblast subsets, generated from PCA. (**G**) Boxplots depicting fold change in fibroblast subset proportions relative to UW across all modalities. (**H**) Boxplots depicting fold change in fibroblast subset proportions relative to UW, separated by whole cell (circles) and single nuclei (triangles) modalities. (**C, D, G, H**) Significance was assessed using the Wilcoxon rank-sum test relative to UW (*p* < 0.05 = *, *p* < 0.01 = **). UW = 7, D4PW = 6, D7PW = 5, D15PW = 4, D30PW = 4.

**Supplemental Figure 11:**
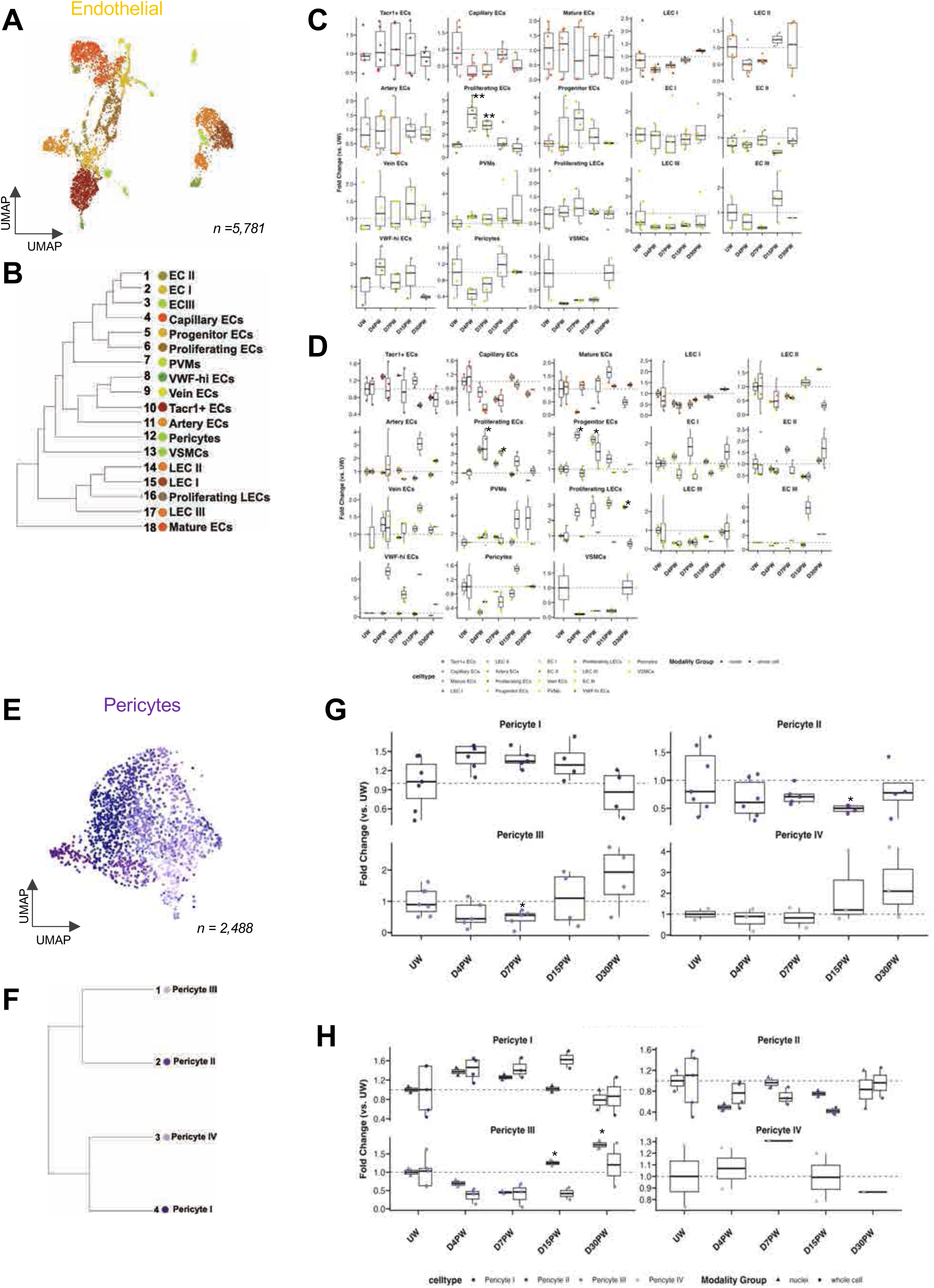
Endothelial and pericyte subsets are heterogeneous throughout wound healing. (**A**) UMAP embedding of endothelial cell subclustering from all timepoints, colored by fine cell type annotation. n = 5,781 cells. (**B**) Dendrogram depicting relationships between endothelial subsets, generated from PCA. (**C**) Boxplots depicting fold change in endothelial subset proportions relative to UW across all modalities. (**D**) Boxplots depicting fold change in endothelial subset proportions relative to UW, separated by whole cell (circles) and single nuclei (triangles) modalities. (**E**) UMAP embedding of pericyte subclustering from all timepoints, colored by fine cell type annotation. n = 2,488 cells. (**F**) Dendrogram depicting relationships between pericyte subsets, generated from PCA. (**G**) Boxplots depicting fold change in pericyte subset proportions relative to UW across all modalities. (**H**) Boxplots depicting fold change in pericyte subset proportions relative to UW, separated by whole cell (circles) and single nuclei (triangles) modalities. (**C, D, G, H**) Significance was assessed using the Wilcoxon rank-sum test relative to UW (*p* < 0.05 = *, *p* < 0.01 = **). UW = 7, D4PW = 6, D7PW = 5, D15PW = 4, D30PW = 4.

**Supplemental Figure 12:**
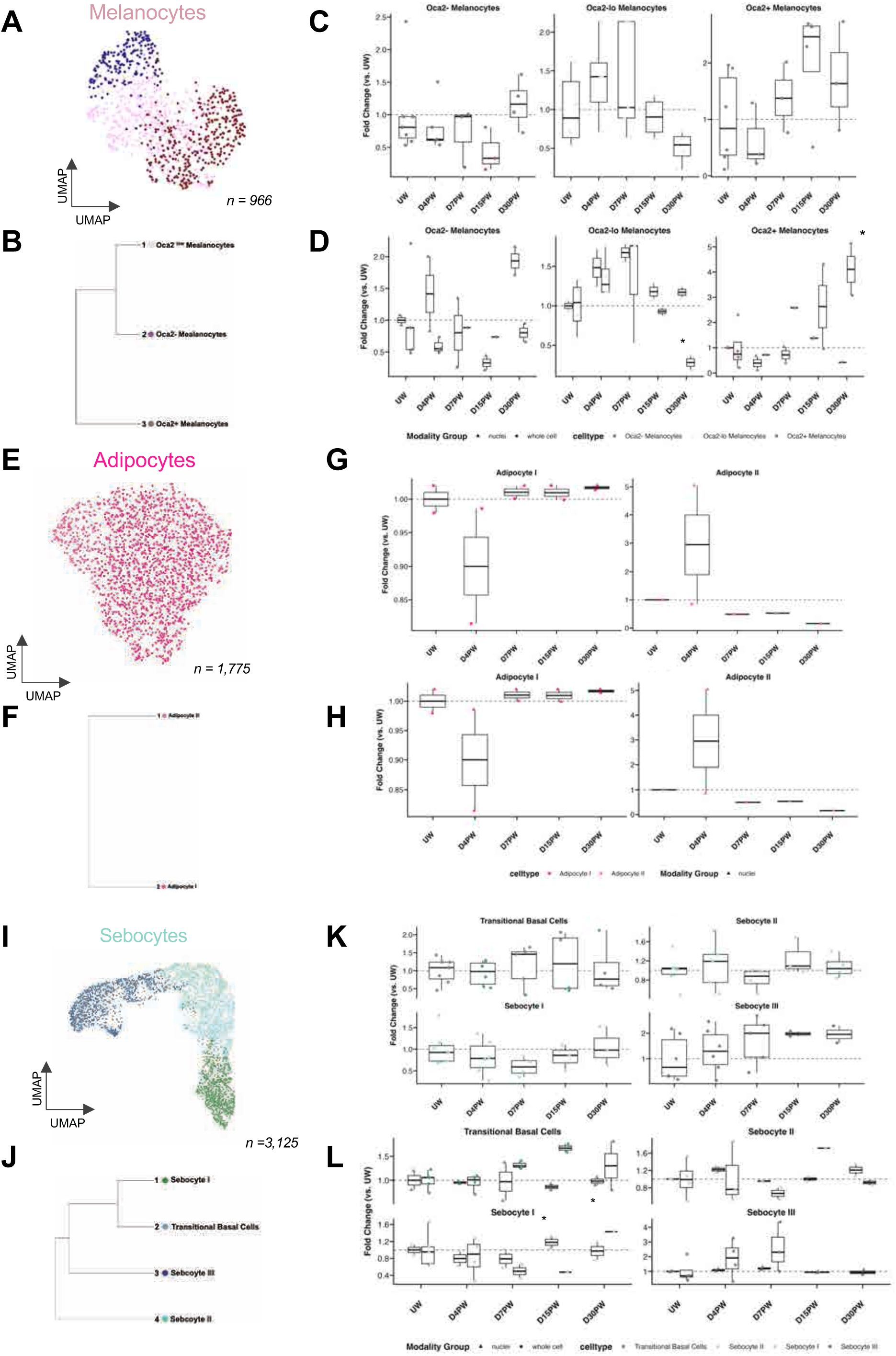
Melanocyte, adipocyte, and pericyte subsets are heterogeneous throughout wound healing. (**A**) UMAP embedding of melanocyte subclustering from all timepoints, colored by fine cell type annotation. n = 966 cells. (**B**) Dendrogram depicting relationships between melanocyte subsets, generated from PCA. (**C**) Boxplots depicting fold change in melanocyte subset proportions relative to UW across all modalities. (**D**) Boxplots depicting fold change in melanocyte subset proportions relative to UW, separated by whole cell (circles) and single nuclei (triangles) modalities. (**E**) UMAP embedding of adipocyte subclustering from all timepoints, colored by fine cell type annotation. n = 1,775 cells. (**F**) Dendrogram depicting relationships between adipocyte subsets, generated from PCA. (**G**) Boxplots depicting fold change in adipocyte subset proportions relative to UW across all modalities. (**H**) Boxplots depicting fold change in adipocyte subset proportions relative to UW, separated by whole cell (circles) and single nuclei (triangles) modalities. (**I**) UMAP embedding of sebocyte sub clustering from all timepoints, colored by fine cell type annotation. n = 3,125 cells. (**J**) Dendrogram depicting relationships between sebocyte subsets, generated from PCA. (**K**) Boxplots depicting fold change in sebocyte subset proportions relative to UW across all modalities. (**L**) Boxplots depicting fold change in sebocyte subset proportions relative to UW, separated by whole cell (circles) and single nuclei (triangles) modalities. (**C, D, G, H, K, L**) Significance was assessed using the Wilcoxon rank-sum test relative to UW (*p* < 0.05 = *, *p* < 0.01 = **). UW = 7, D4PW = 6, D7PW = 5, D15PW = 4, D30PW = 4.

**Supplemental Figure 13:**
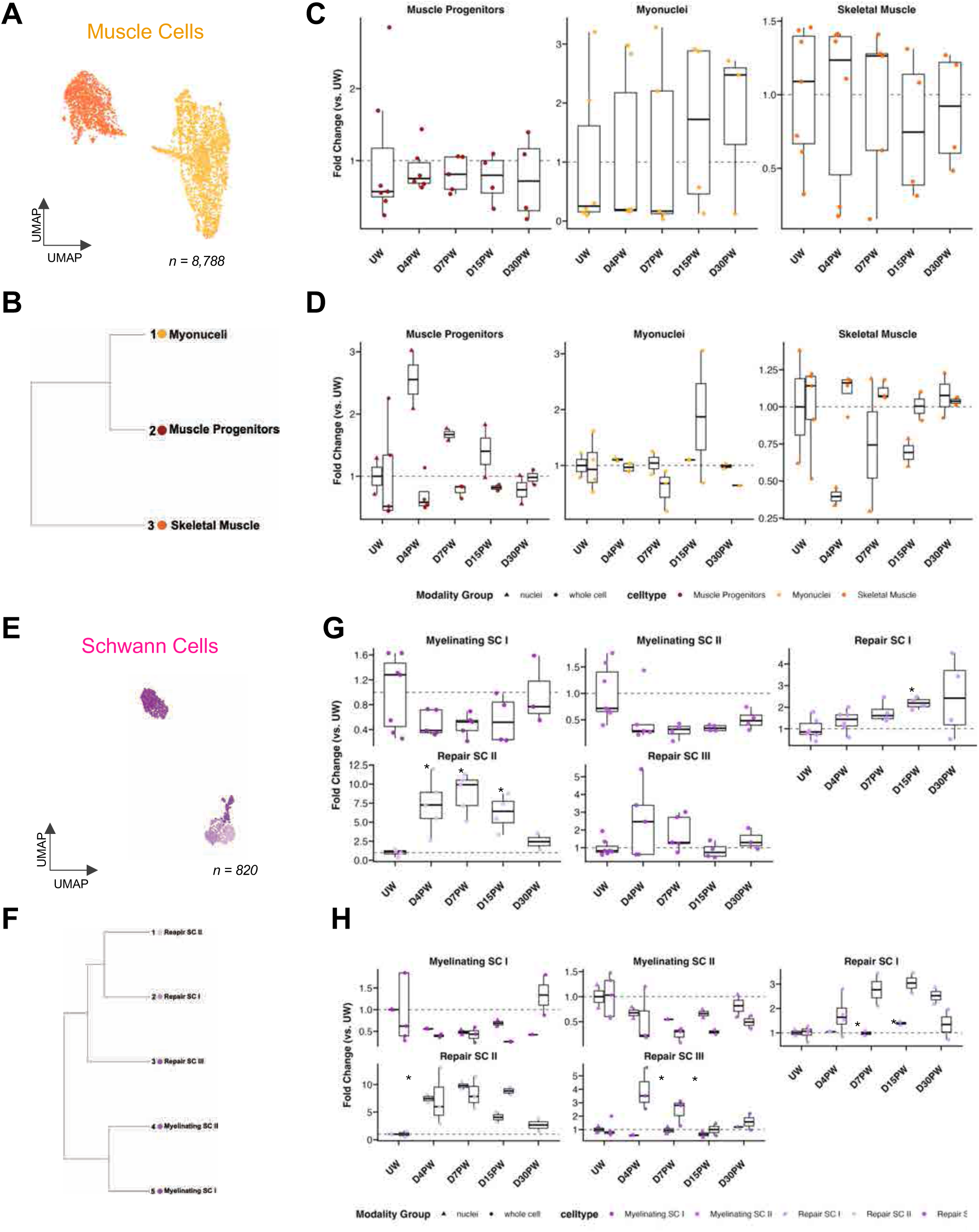
Endothelial, pericyte, Schwann Cell subsets are heterogeneous throughout wound healing. (**A**) UMAP embedding of muscle cell sub clustering from all timepoints, colored by fine cell type annotation. n = 5,781 cells. (**B**) Dendrogram depicting relationships between muscle subsets, generated from PCA. (**C**) Boxplots depicting fold change in muscle subset proportions relative to UW across all modalities. (**D**) Boxplots depicting fold change in muscle subset proportions relative to UW, separated by whole cell (circles) and single nuclei (triangles) modalities. (**E**) UMAP embedding of Schwann cell subclustering from all timepoints, colored by fine cell type annotation. n = 820 cells. (**F**) Dendrogram depicting relationships between Schwann cell subsets, generated from PCA. (**G**) Boxplots depicting fold change in Schwann cell subset proportions relative to UW across all modalities. (**H**) Boxplots depicting fold change in Schwann cell subset proportions relative to UW, separated by whole cell (circles) and single nuclei (triangles) modalities. (**C, D, G, H**) Significance was assessed using the Wilcoxon rank-sum test relative to UW (*p* < 0.05 = *, *p* < 0.01 = **). UW = 7, D4PW = 6, D7PW = 5, D15PW = 4, D30PW = 4.

**Supplemental Figure 14:**
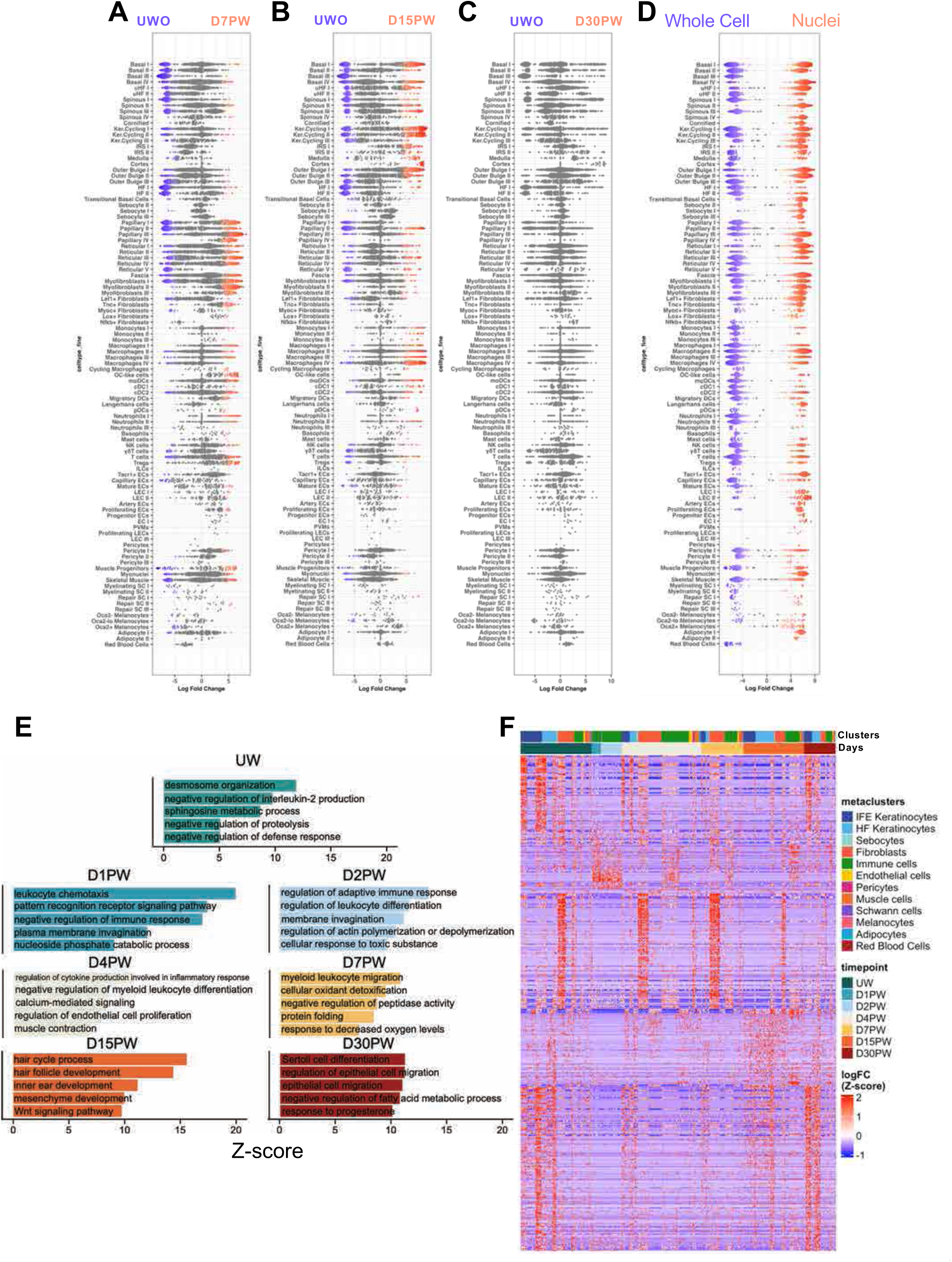
Defining a transcriptomic signature of wound healing by differential abundance and differential gene expression. **(A-C)** Beeswarm plots of Milo differential abundance analysis across wound conditions compared to UW (UW vs. D7PW, UW vs. D15PW, and UW vs. D30PW). Colored points indicate FDR ≤ 0.10. **(D)** Beeswarm plots of Milo differential abundance analysis comparing whole cell and nuclei modalities across all wounding timepoints. Colored points indicate FDR ≤ 0.10. **(E)** Bar plots showing z-scores of the top 5 Gene Ontology Biological Process (BP) pathways at each respective timepoint relative to all other timepoints. z-scores calculated using clusterProfiler (cutoff *p* < 0.05). **(F)** Heatmap displaying top 200 upregulated and downregulated genes for each phase transitions across metaclusters i.e. UWO vs D4PW, D4PW vs D7PW, D7PW vs D15PW, D15PW vs D30PW.

**Supplemental Figure 15:**
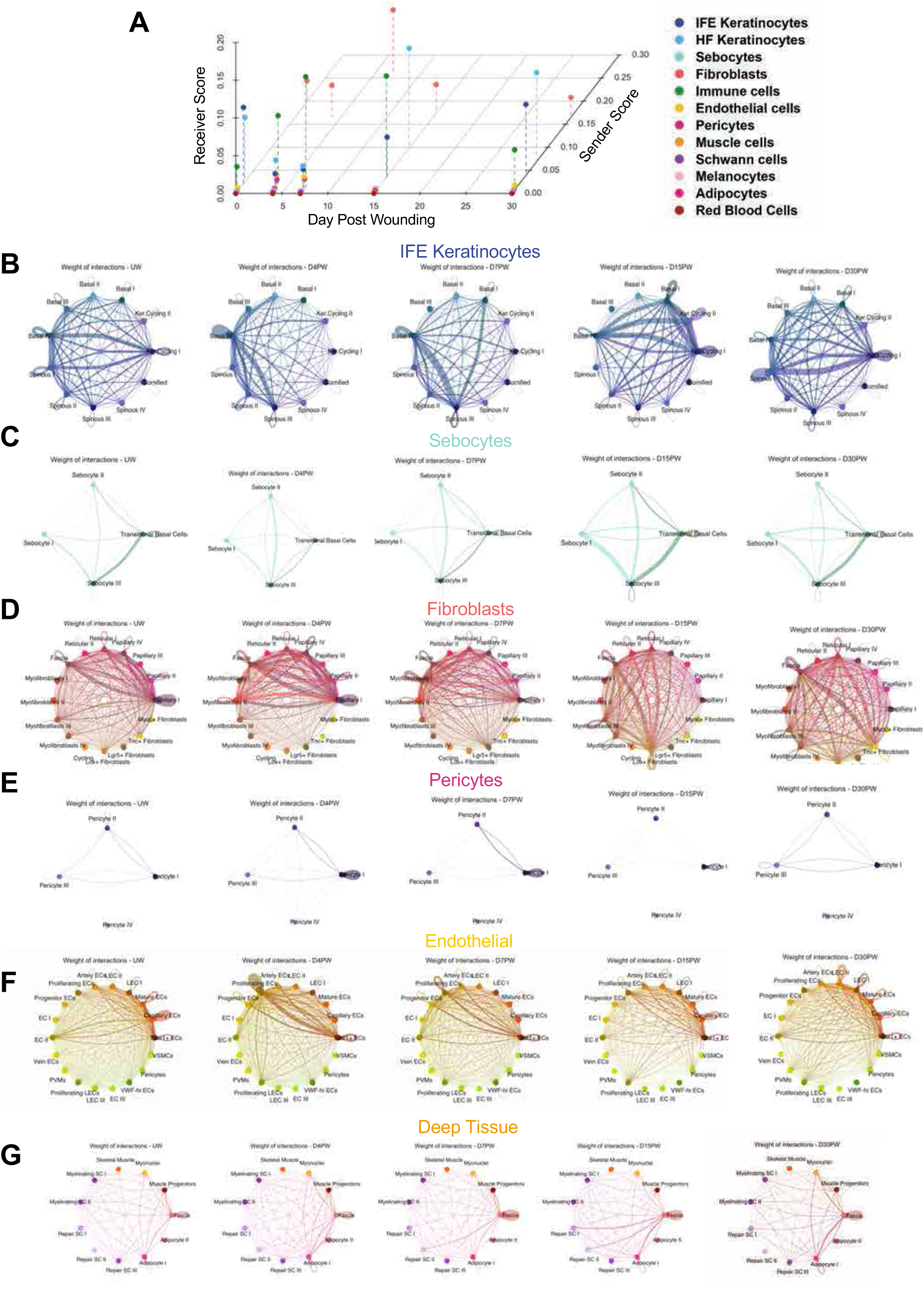
Subclusters contain dominant signaling nodes for each major metacluster. (**A**) 3D scatter plot summarizing signaling sender and receiver scores over time for each metacluster. **(B–G)** CellChat circle plots displaying the weighted ligand–receptor communication between subclusters within each major metacluster population. Shown are: (**B**) Keratinocytes, (**C**) Sebocytes, (**D**) Fibroblasts, (**E**) Pericytes, (**F**) Endothelial cells, and **(G)** Deep tissue structures. Width of edges between groups denote strength of signaling interactions.

**Supplemental Figure 16:**
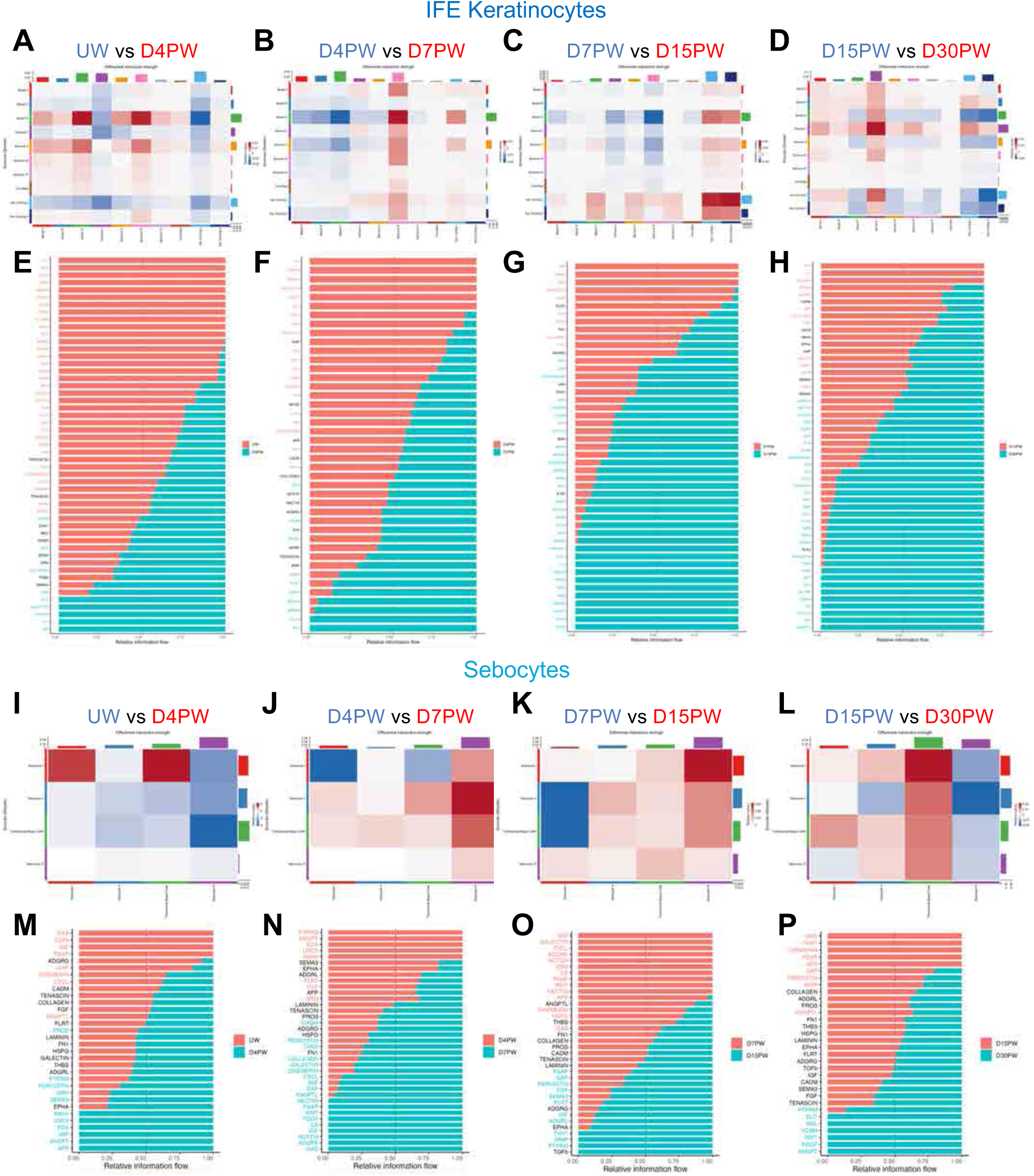
Dynamic signaling programs in IFE keratinocyte and sebocyte subclusters during wound healing. (**A-D**) Differential CellChat heatmaps displaying changes in interaction strength between IFE keratinocyte subsets over sequential healing timepoints: (**A**) UW vs. D4PW, (**B**) D4PW vs. D7PW, (**C**) D7PW vs. D15PW, (**D**) D15PW vs. D30PW. Color indicates relative increase or decrease in interaction strength between populations as indicated by colored text. (**E-H**) Stacked barplots depicting relative strength of signaling pathway between all IFE keratinocytes subsets over sequential healing timepoints: (**E**) UW vs. D4PW, (**F**) D4PW vs. D7PW, (**G**) D7PW vs. D15PW, (**H**) D15PW vs. D30PW. Colored text indicates statistically significant enrichment of a given pathway at the indicated timepoint (Wilcoxon test, *p* < 0.05). (**I-L**) Differential CellChat heatmaps displaying changes in interaction strength between sebocyte subsets over sequential healing timepoints: (**I**) UW vs. D4PW, (**J**) D4PW vs. D7PW, (**K**) D7PW vs. D15PW, (**L**) D15PW vs. D30PW. Color indicates relative increase or decrease in interaction strength between populations as indicated by colored text. (**M-P**) Stacked barplots depicting relative strength of signaling pathway between all sebocyte subsets over sequential healing timepoints: (**M**) UW vs. D4PW, (**N**) D4PW vs. D7PW, (**O**) D7PW vs. D15PW, (**P**) D15PW vs. D30PW. Colored text indicates statistically significant enrichment of a given pathway at the indicated timepoint (Wilcoxon test, *p* < 0.05).

**Supplemental Figure 17:**
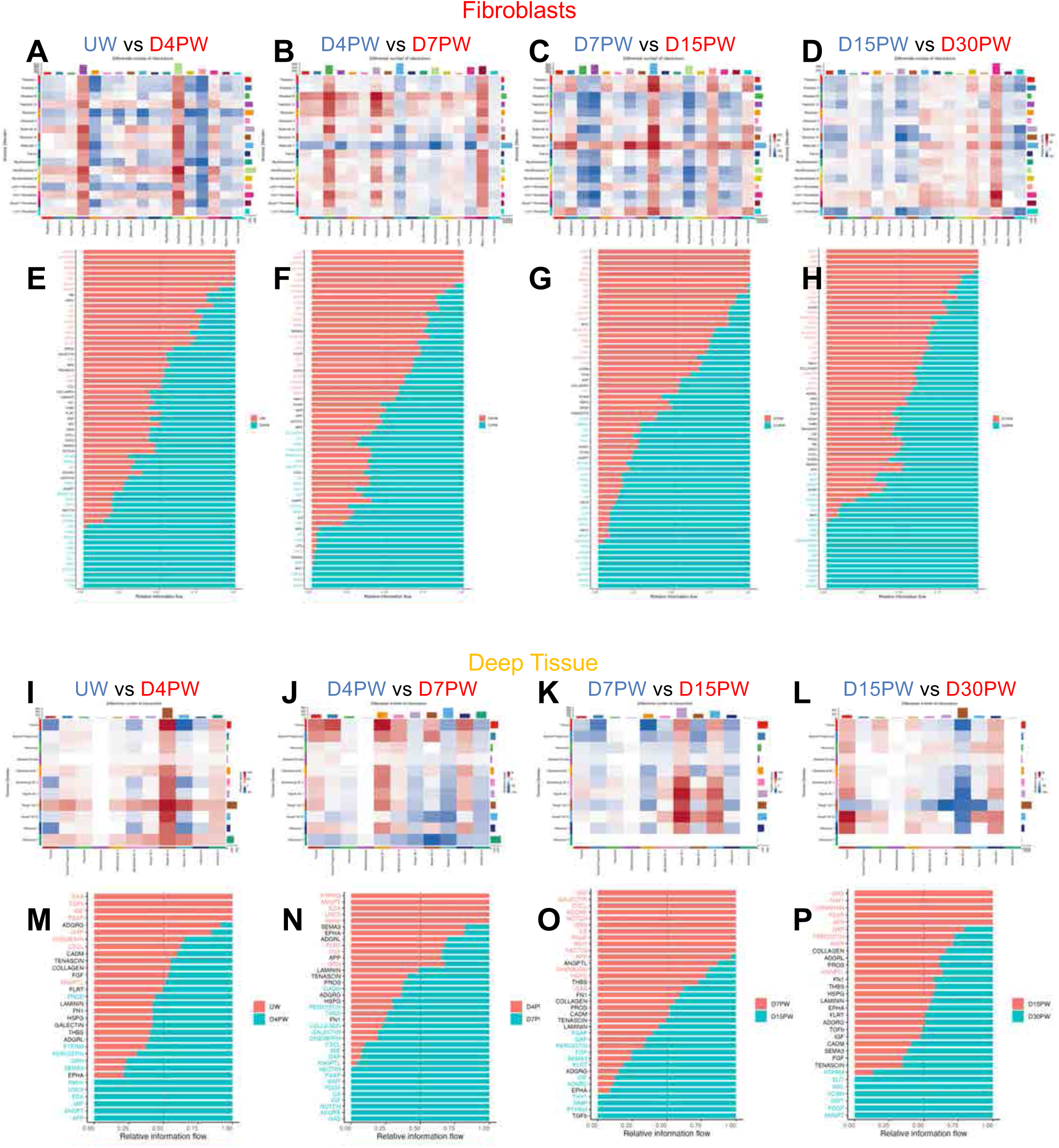
Dynamic signaling programs in fibroblasts and deep tissue subclusters during wound healing. (**A-D**) Differential CellChat heatmaps displaying changes in interaction strength between fibroblast subsets over sequential healing timepoints: (**A**) UW vs. D4PW, (**B**) D4PW vs. D7PW, (**C**) D7PW vs. D15PW, (**D**) D15PW vs. D30PW. Color indicates relative increase or decrease in interaction strength between populations as indicated by colored text. (**E-H**) Stacked barplots depicting relative strength of signaling pathway between all fibroblast subsets over sequential healing timepoints: (**E**) UW vs. D4PW, (**F**) D4PW vs. D7PW, (**G**) D7PW vs. D15PW, (**H**) D15PW vs. D30PW. Colored text indicates statistically significant enrichment of a given pathway at the indicated timepoint (Wilcoxon test, *p* < 0.05). (**I-L**) Differential CellChat heatmaps displaying changes in interaction strength between Deep Tissue subsets over sequential healing timepoints: (**I**) UW vs. D4PW, (**J**) D4PW vs. D7PW, (**K**) D7PW vs. D15PW, (**L**) D15PW vs. D30PW. Color indicates relative increase or decrease in interaction strength between populations as indicated by colored text. (**M-P**) Stacked barplots depicting relative strength of signaling pathway between all Deep Tissue subsets over sequential healing timepoints: (**M**) UW vs. D4PW, (**N**) D4PW vs. D7PW, (**O**) D7PW vs. D15PW, (**P**) D15PW vs. D30PW. Colored text indicates statistically significant enrichment of a given pathway at the indicated timepoint (Wilcoxon test, *p* < 0.05).

**Supplemental Figure 18:**
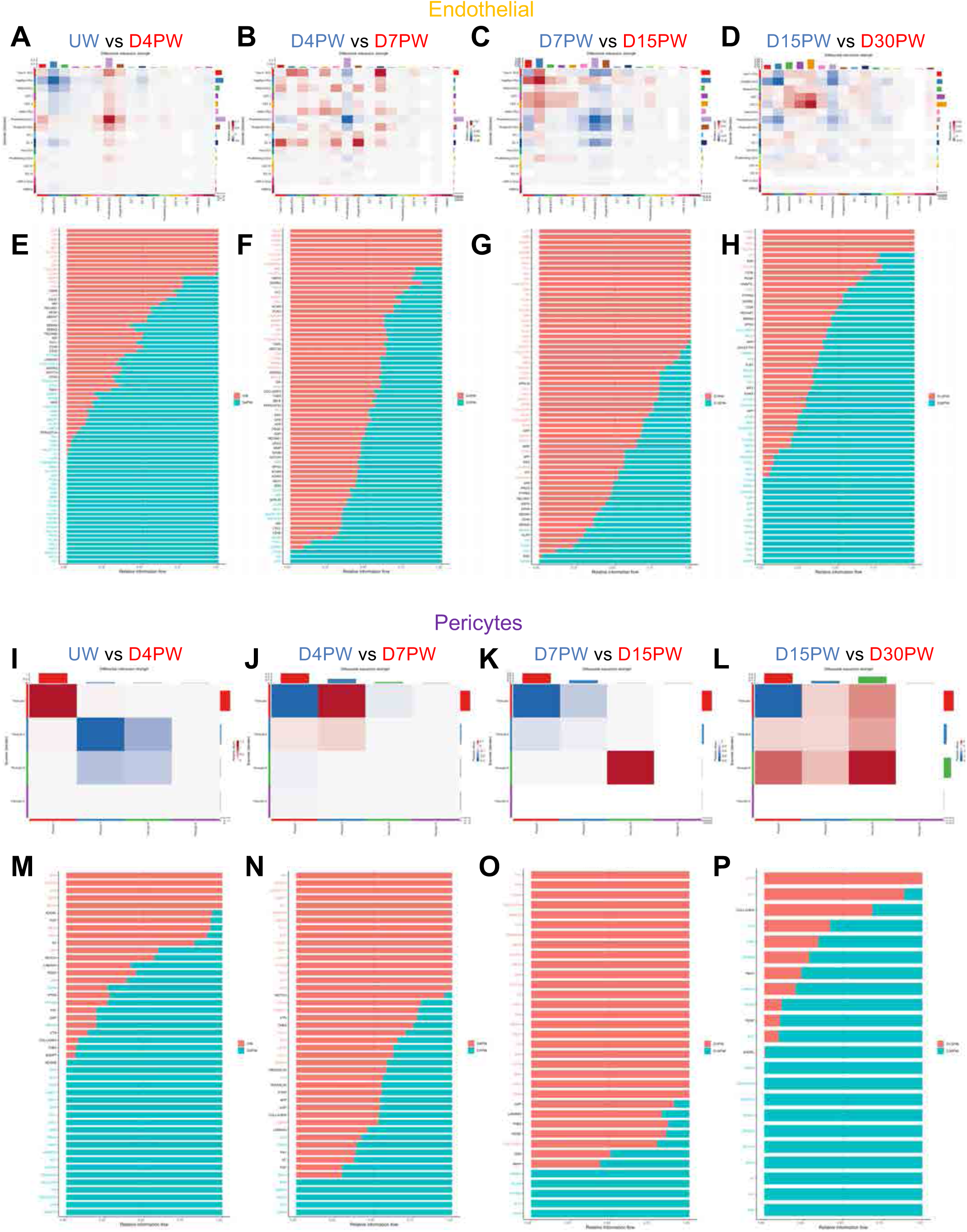
Dynamic signaling programs in endothelial and pericytes subclusters during wound healing. (**A-D**) Differential CellChat heatmaps displaying changes in interaction strength between endothelial cell subsets over sequential healing timepoints: (**A**) UW vs. D4PW, (**B**) D4PW vs. D7PW, (**C**) D7PW vs. D15PW, (**D**) D15PW vs. D30PW. Color indicates relative increase or decrease in interaction strength between populations as indicated by colored text. (**E-H**) Stacked barplots depicting relative strength of signaling pathway between all endothelial subsets over sequential healing timepoints: (**E**) UW vs. D4PW, (**F**) D4PW vs. D7PW, (**G**) D7PW vs. D15PW, (**H**) D15PW vs. D30PW. Colored text indicates statistically significant enrichment of a given pathway at the indicated timepoint (Wilcoxon test, *p* < 0.05). (**I-L**) Differential CellChat heatmaps displaying changes in interaction strength between pericyte subsets over sequential healing timepoints: (**I**) UW vs. D4PW, (**J**) D4PW vs. D7PW, (**K**) D7PW vs. D15PW, (**L**) D15PW vs. D30PW. Color indicates relative increase or decrease in interaction strength between populations as indicated by colored text. (**M-P**) Stacked barplots depicting relative strength of signaling pathway between all pericyte subsets over sequential healing timepoints: (**M**) UW vs. D4PW, (**N**) D4PW vs. D7PW, (**O**) D7PW vs. D15PW, (**P**) D15PW vs. D30PW. Colored text indicates statistically significant enrichment of a given pathway at the indicated timepoint (Wilcoxon test, *p* < 0.05).

**Supplemental Figure 19.**
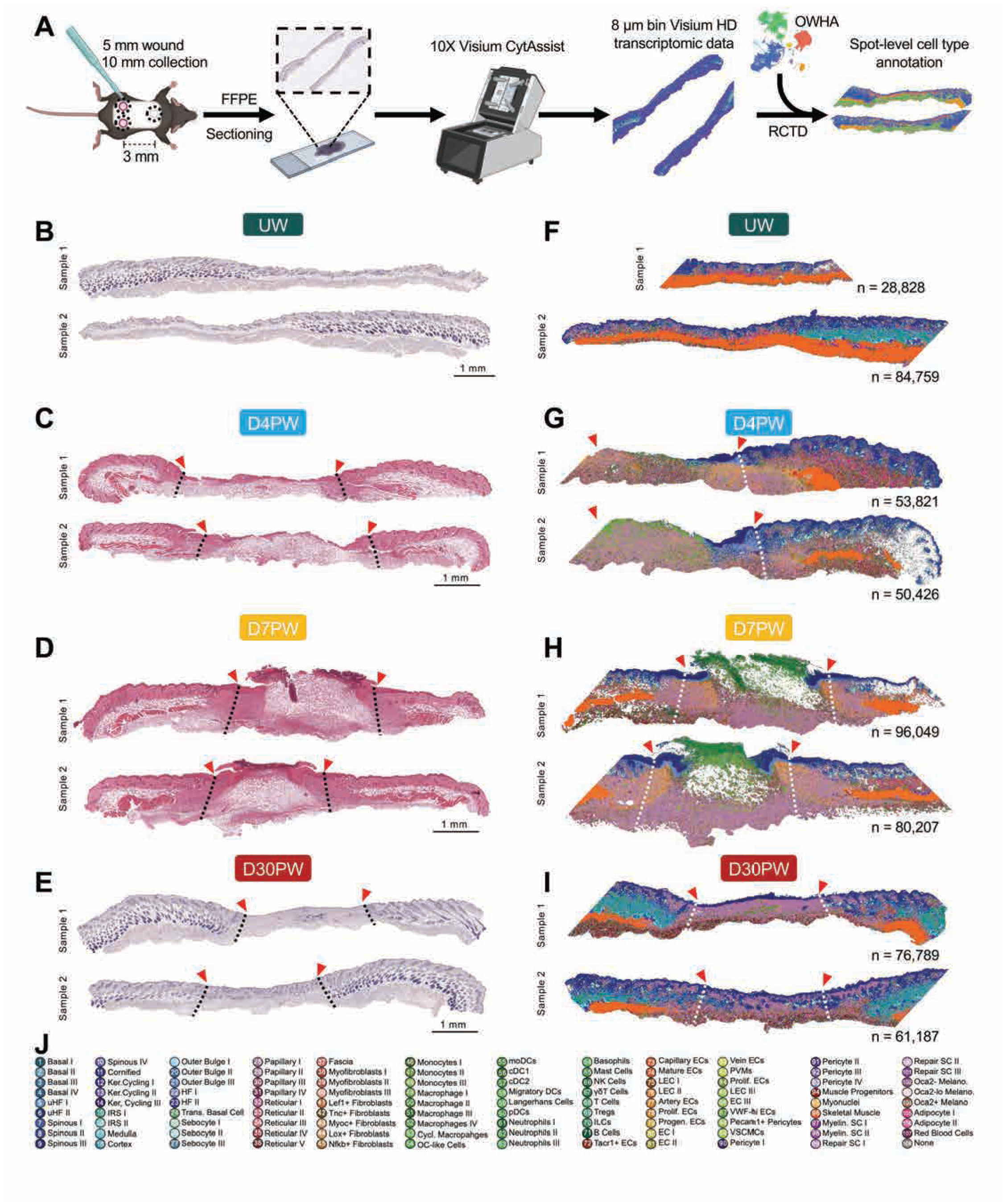
Visium HD provides high resolution spatial transcriptomics information across wound healing timeline. **(A)** Schematic overview of Visium HD preparation from mouse wounds. Robust cell type deconvolution (RCTD) was used with the OWHA dataset as a reference to annotate spots. **(B-E)** Hematoxylin & Eosin (H&E) stains of sections used for Visium HD from unwounded (UW) (B), D4PW (C), D7PW (D), and D30PW (E) skin sections. Sections represent paired tissues from same animal and wound site. Red triangles and dashed lines denote suprabasal and dermal wound edges, respectively. **(F-I)** Fine cell type annotations from RCTD, corresponding to cell types captured in OWHA from UW (F), D4PW (G), D7PW (D), and D30PW (E) skin sections. n = Number of 8 μm spots captured in each tissue section shown. Red triangles and dashed lines denote suprabasal and dermal wound edges, respectively. (**J**) Numbering and classification legend of fine cell types corresponding to panels F-I.

**Supplemental Figure 20:**
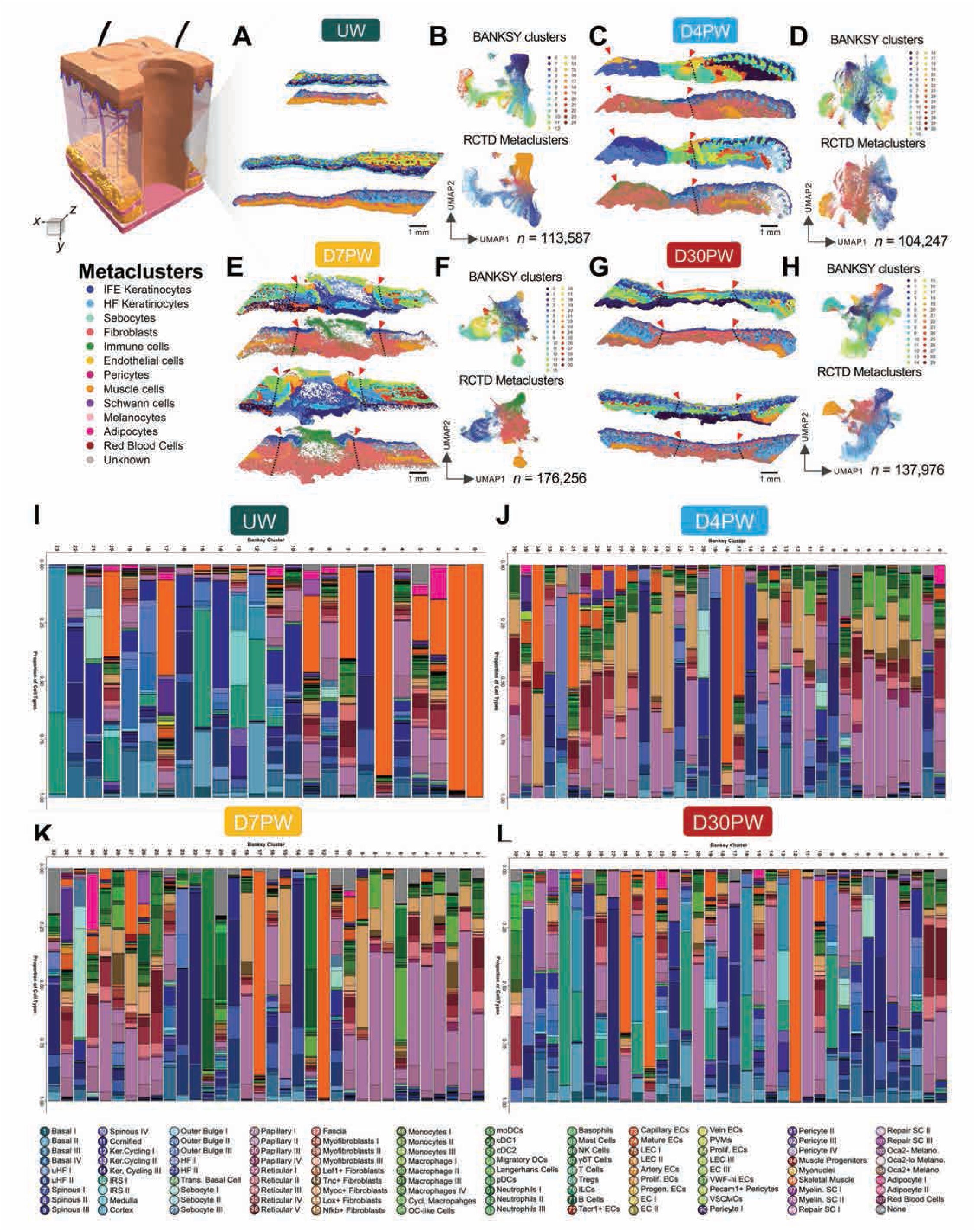
BANKSY spatially informed clustering identifies localized spatial niches across wound healing timeline. **(A, C, E, and G)** Spatial plots of BANKSY clusters and RCTD Metaclusters in replicate tissue sections from unwounded (**A**), D4PW (**C**), D7PW (**E**), and D30PW (**G**). **(B, D, F, and H)** UMAP plots of BANKSY clusters (top) and RCTD Metaclusters (bottom) from unwounded (**B**), D4PW (**D**), D7PW (**F**), and D30PW (**H**). **n** = total number of spots captured by both tissue sections. **(I-L)** Bar plots showing proportional representation of fine cell types in BANKSY clusters for UW (**I**), D4PW (**J**), D7PW (**K**), and D30PW (**L**) Visium HD samples. Plots shown as total cell type proportions across 2 technical replicate tissue sections.

**Supplemental Figure 21.**
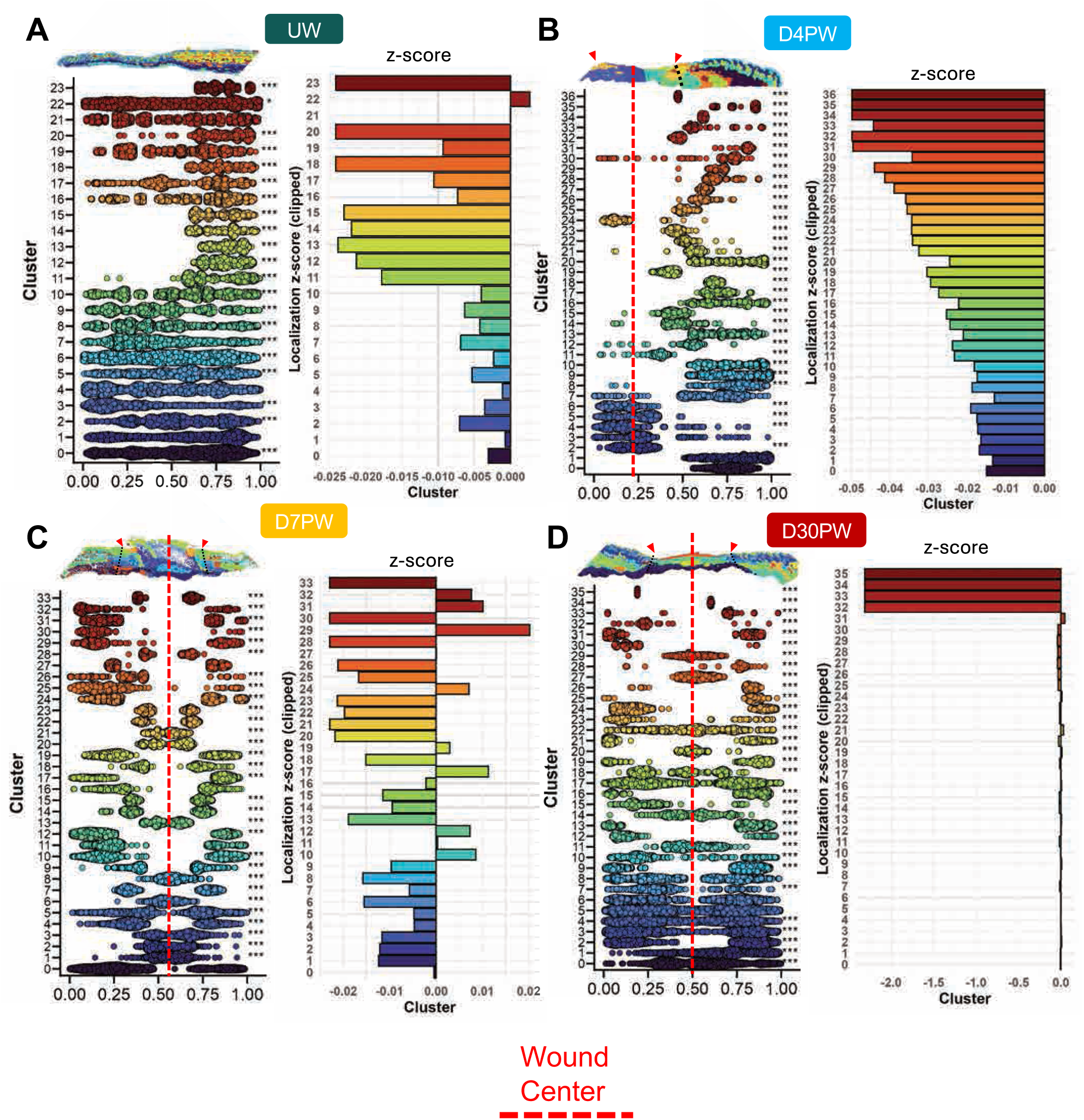
BANKSY spatial niches localize to specific regions of the wound site. (**A-D**) Beeswarm plots of BANKSY cluster localization on normalized tissue length axis (left), and z-scores of localizations relative to null-distribution cluster (right) for UW (**A**), D4PW (**B**), D7PW (**C**), D30PW **(D**) samples. Red dashed lines denote relative location of wound center in tissue sections. Significance was evaluated using a bootstrap test for small clusters and a chi-square test for large clusters, comparing observed variance in cluster localization to a hypothetical uniformly distributed cluster. Multiple testing correction was applied to p-values using the Benjamini–Hochberg procedure, p < 0.05 *, p < 0.01 **, p < 0.001 ***.

**Supplemental Figure 22:**
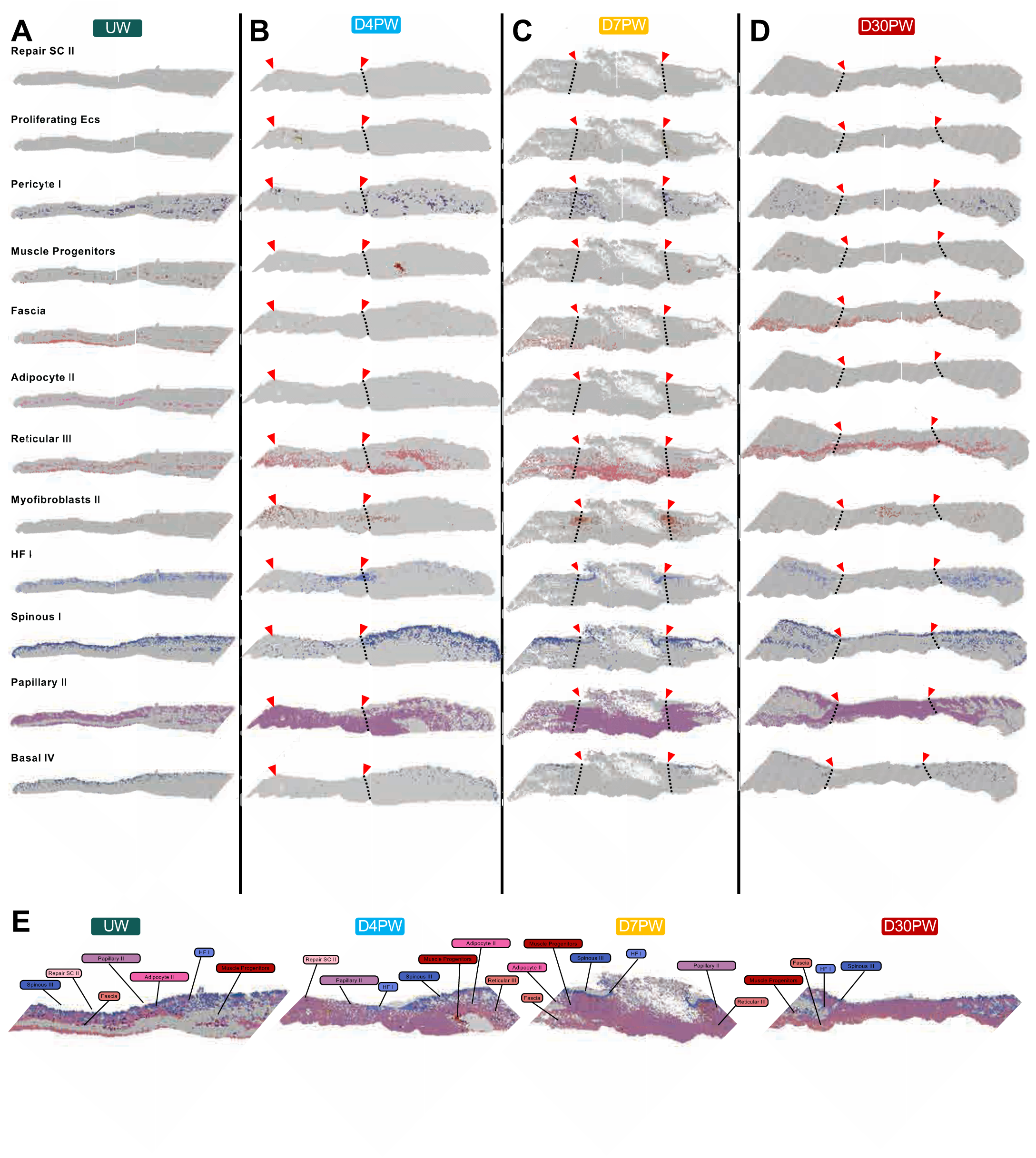
Emergent cell types localize to distinct regions of healing skin in Visium HD data. **(A-D)** Spatial plots highlighting individual emergent cell types in Visium HD data for UW (A), D4PW (B), D7PW (C), and D30PW (D) samples. Plots show representative single tissue section. Red triangles and dashed lines denote suprabasal and dermal wound edges, respectively. **(E)** Spatial plots highlighting combined emergent cell types in Visium HD data for UW (A), D4PW (B), D7PW (C), and D30PW (D) samples. Plots highlight colocalization of emergent cell types in specific wound zones.

**Supplemental Figure 23.**
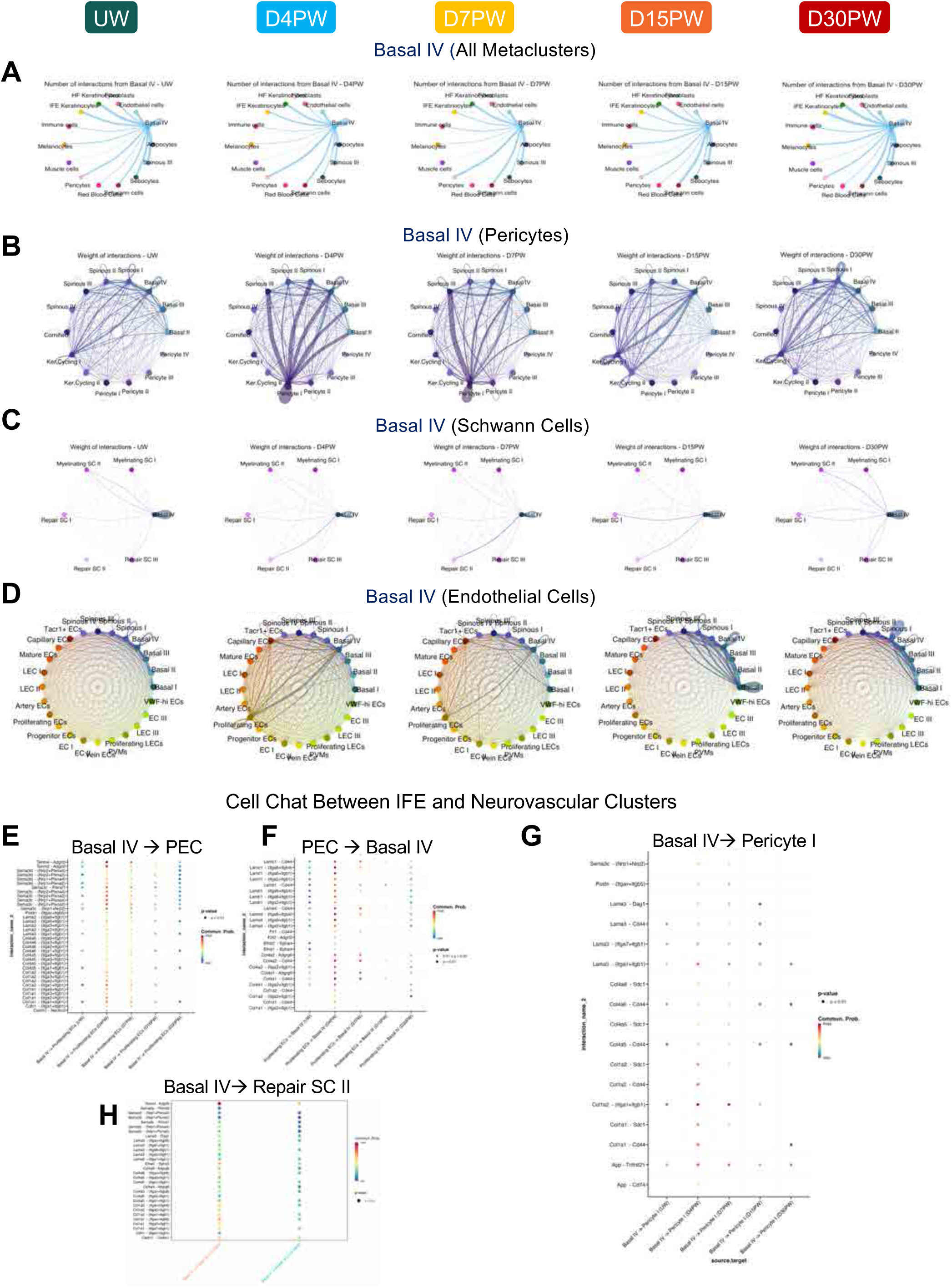
Basal IV keratinocytes interact with distinct populations throughout wound healing. **(A)** CellChat circle plot showing the number of ligand–receptor interactions sent by Basal IV keratinocytes other major Metaclusters over time. **(B)** CellChat circle plot showing the interaction weights between IFE subclusters and pericytes over time. **(C)** CellChat circle plot showing the interaction weights between Basal IV cells and Schwann cell subclusters over time. **(D)** CellChat circle plot showing the interaction weights between IFE subclusters and endothelial subclusters over time. **(E-G)** Dot plots showing signaling strength over time for ligand-receptor pairs sent and received by Basal IV and subclusters associated with neurovascular signaling hub: (**E**) sent by Basal IV and received by proliferating endothelial cells (PEC), (**F**) sent by PEC and received by Basal IV, (**G**) sent by Basal IV and received by pericyte I. (**H**) Dot plot showing relative signaling strength of pathways between Basal IV and Repair SC II cells at D7PW and D15PW.

**Supplemental Figure 24.**
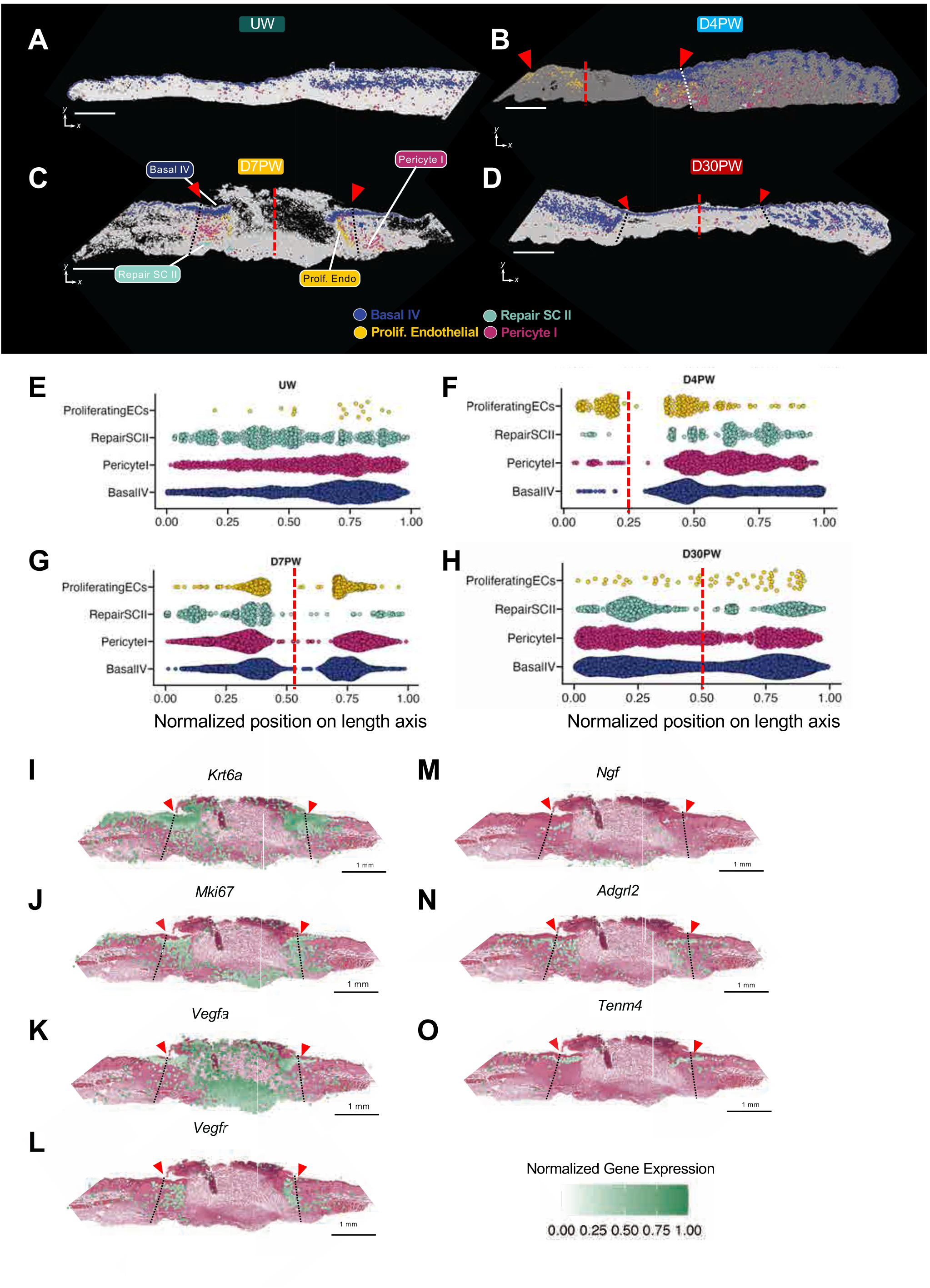
Emergent CO populations colocalize in distinct spatial niches during the inflammation-proliferation phase transition. (**A-D)** Visium HD spatial localization of Basal IV (blue), proliferative endothelial cells (yellow), Repair SCII (teal), Pericyte I (pink) throughout the wound healing time course (UW, D4PW, D7PW, D30PW). Red arrowheads indicate the wound edge in the suprabasal layer, while red dashed lines mark center of the wound. Black dashed lines denote the dermal wound boundaries. This represents one biological replicate per timepoint and shows one representative technical replicate of two. Scale bar represents 1mm **(E-H)** Anatomical distance quantifications of Basal IV (blue), proliferative cells, Repair SCII (teal), Pericyte I (pink) across the anterior-posterior wound plane. The x-axis represents arbitrary spatial units (1 a.u. = 9 mm) corresponding to anterior-posterior distance across the tissue section. Represents two technical replicates per timepoint. Red lines denote approximate location of wound center. **(I-O)** Spatial transcriptomic expression of related angiogenesis and axonogenesis axis (Krt6a (I), Mki67 (J), Vegfa (K), Vegfr (L), Ngf (M), Adgrl2 (N), Tenm4(O)), overlayed with H&E images in D7PW Visium HD sections. Red triangles and dashed lines denote wound edge in epidermis and dermis, respectively.

**Supplemental Figure 25.**
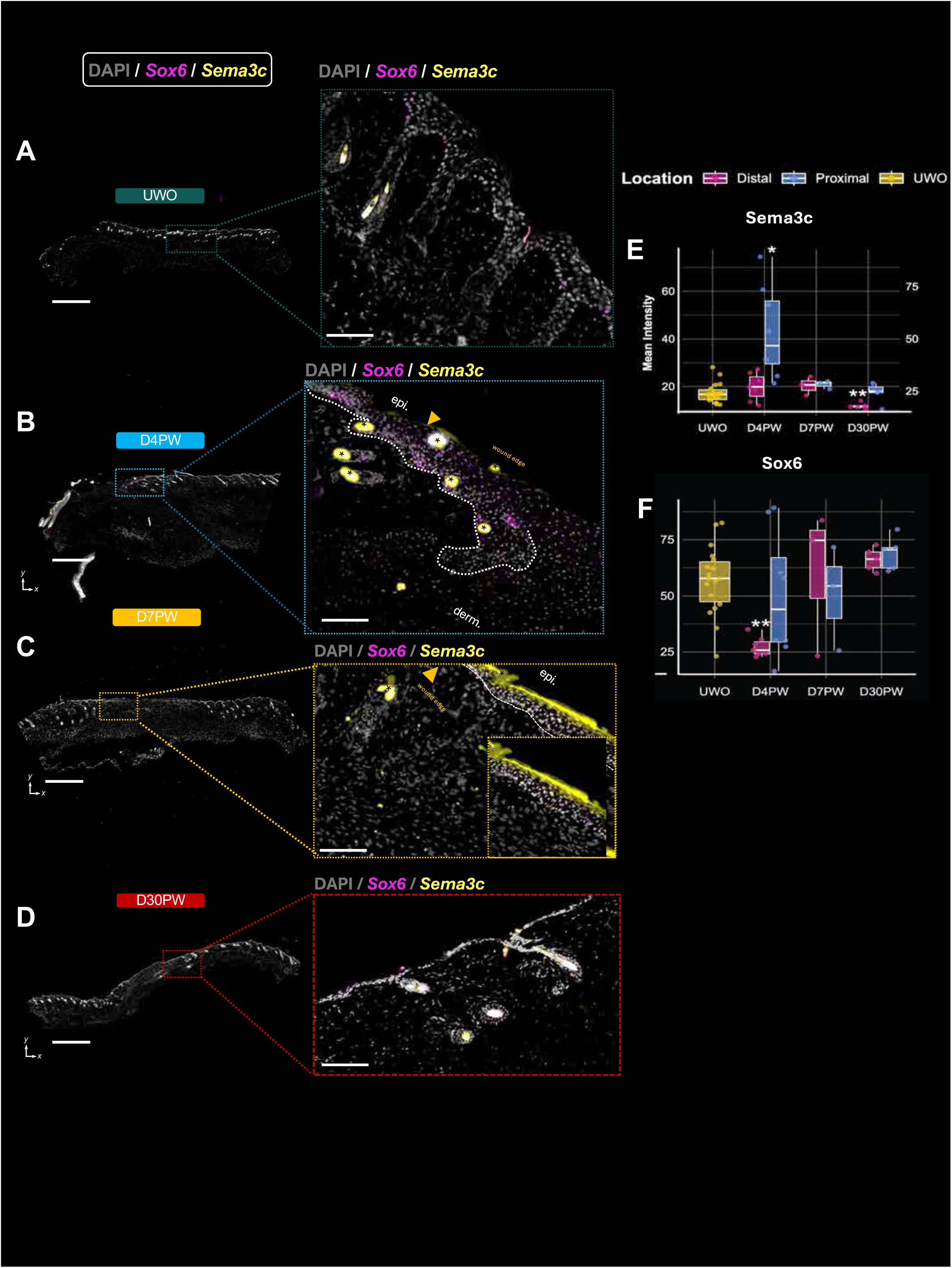
Sema3c colocalizes with Sox6 in the epithelium proximal to the wound site after injury. (**A-D)** Single-molecule RNA FISH (smFISH) and immunofluorescence analysis of Sox6 (magenta) and Sema3C (yellow) mRNA transcripts, merged with DAPI nuclear staining (white) in **(A)** UW, (**B**) D4PW, (C) D7PW, and (D) D30PW skin. Left image displays 10X large image scan (scale bar represents 1mm) and callouts on right display 20X image of zoomed in sections (scale bar represents 10mm). **(E–F)** Boxplots quantifying mean fluorescence intensity of (E) Sox6 and (F) Sema3c smFISH signals across timepoints (UW, D4PW, D7PW, D30PW), comparing unwounded, distal, and proximal wound regions. UW includes three biological replicates, D4PW two biological replicates, and D7PW/D30PW representing one biological replicate. Variability is represented using the interquartile range (IQR). Statistical significance was determined using a Wilcoxon rank-sum test (p < 0.05 = *, p < 0.01 = **).

**Supplemental Figure 26.**
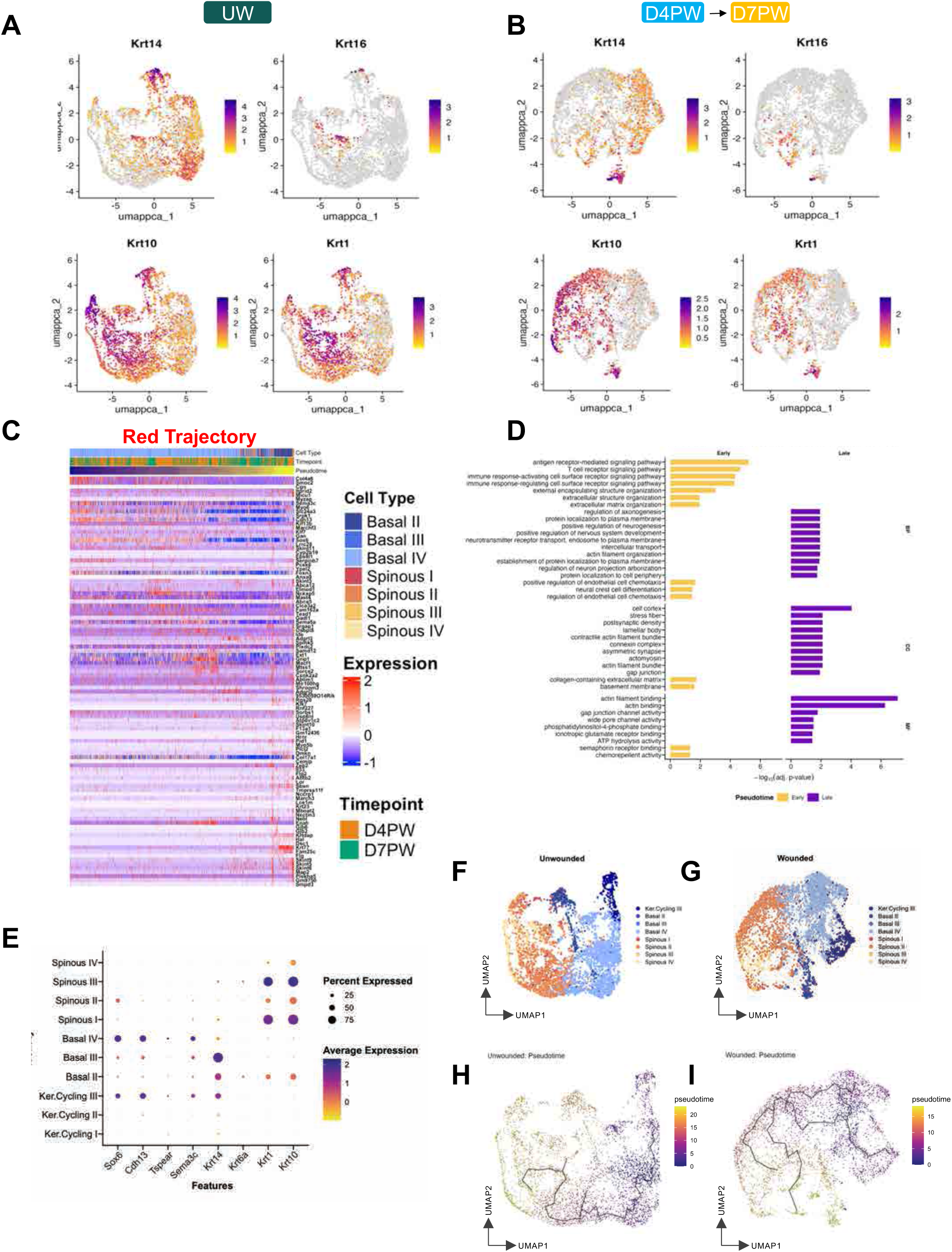
Pseudotime dynamics and transcriptional programs of IFE keratinocytes. **(A–B)** Feature plots showing the expression of Krt14, Krt16, Krt10, and Krt1 across the snRNA-seq UMAP, illustrating basal-to-differentiation transitions in the keratinocyte compartment. (A) Representing unwounded and (B) representing both D4PW and D7PW timepoints. (**C**) Heatmap of the top 50 pseudotime-associated genes in the D4PW/D7PW “red” trajectory keratinocytes, ordered by subcluster and pseudotime value. Arrows indicate genes enriched in early pseudotime (purple) versus late pseudotime (yellow), marking progressive transcriptional state transitions. (**D**) Gene Ontology enrichment results (BP, CC, MF) for genes associated with early and late pseudotime, highlighting distinct biological programs active at different stages of keratinocyte progression. (**E**) DotPlot showing expression of Basal IV, spinous, and pan-keratinocyte markers in IFE subsets across all timepoints. (**F-G**) UMAP plots depicting IFE keratinocytes including keratinocyte cycling cell separated into (**F**) unwounded and (**G**) early wounding (D4PW/D7PW) timepoints, colored by cell type annotation. (**H-I**) UMAP plots showing pseudotemporal ordering of IFE keratinocyte including keratinocyte cycling cell subsets, separated into (**H**) unwounded and (**I**) early wounding (D4PW/D7PW) timepoints.

**Supplemental Figure 27.**
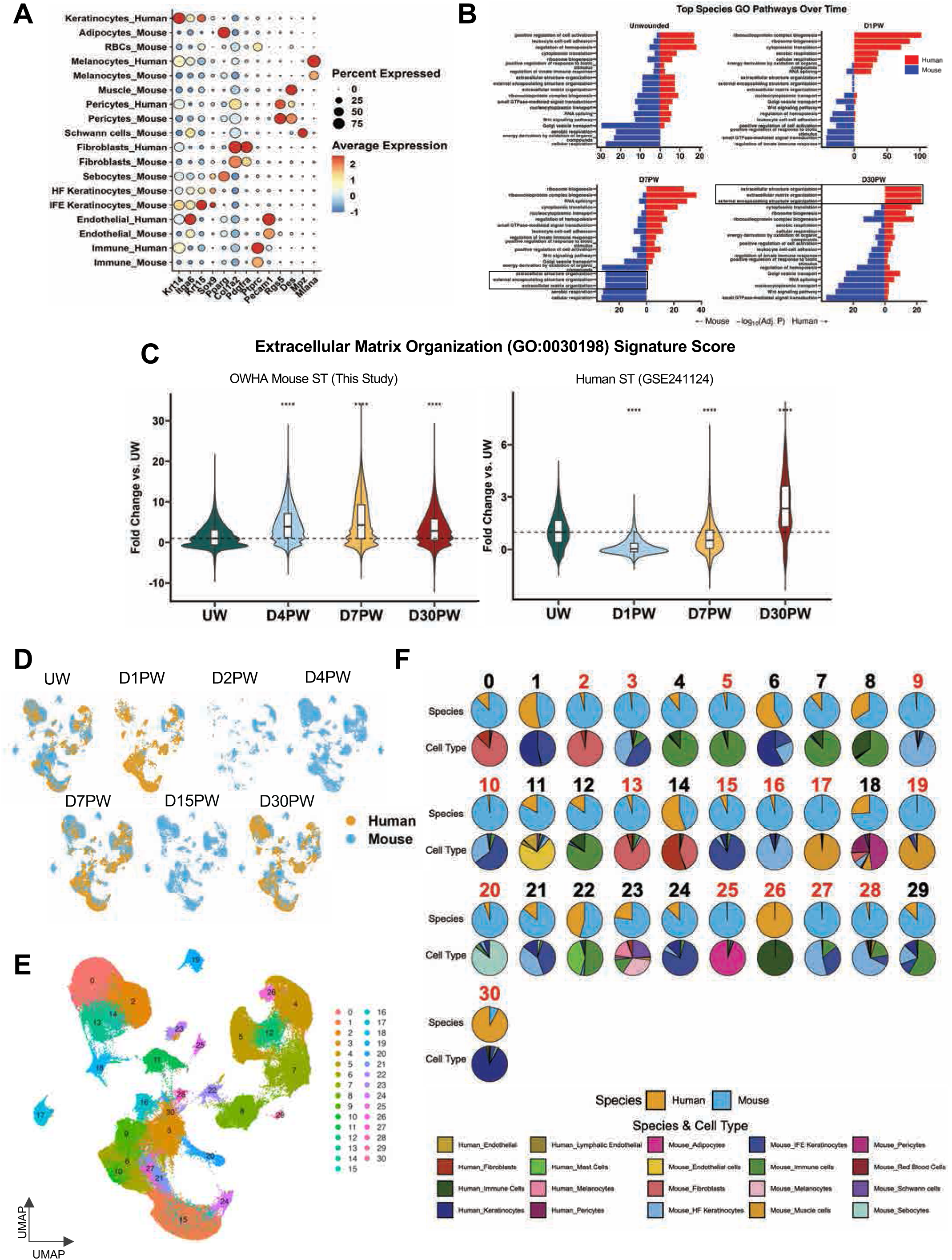
Cross-Species Comparison of Wound-Healing Transcriptional Programs. (**A**) Dot plot showing canonical marker gene expression across major metaclusters in both mouse and human datasets. (**B**) Bar plots of top Gene Ontology Biological Process pathways for both mouse and human across matching timepoints, showing pathways enriched in one species or another at the same healing time. Plots depict adjusted p-value of enriched pathway for each species within one timepoint, with direction of enrichment denoting species-specific induction. Black boxes highlight ECM-associated pathways which display temporal divergence, demonstrating that active repair pathways remain enriched in human wounds compared to mouse. (**C**) Violin plots depicting fold change in extracellular matrix organization GO pathway signature score in mouse ST data (This study, left) and human ST data (GEO accession GSE241124, right). Significance was assessed using Wilcoxon rank-sum test (p < 0.05 = *, p < 0.01 = **, p < 0.001 = ***). Mouse n = 1, human n = 4. (**D**) UMAP projections of the integrated cross-species wound-healing atlas across all timepoints (UW, D1PW, D2PW, D4PW, D7PW, D15PW, D30PW), colored by species. Depicts species composition at each timepoint within the 236,930-cell, 40-sample integrated atlas. (**E**) UMAP plot of the integrated cross-species atlas, colored by Harmony-integrated clustering. (**F**) Pie charts showing the species composition (mouse: blue; human: orange) and major cell-type composition for each Harmony cluster in the cross-species atlas. Clusters composed of >90% cells from a single species are labeled in red.

**Supplemental Figure 28.**
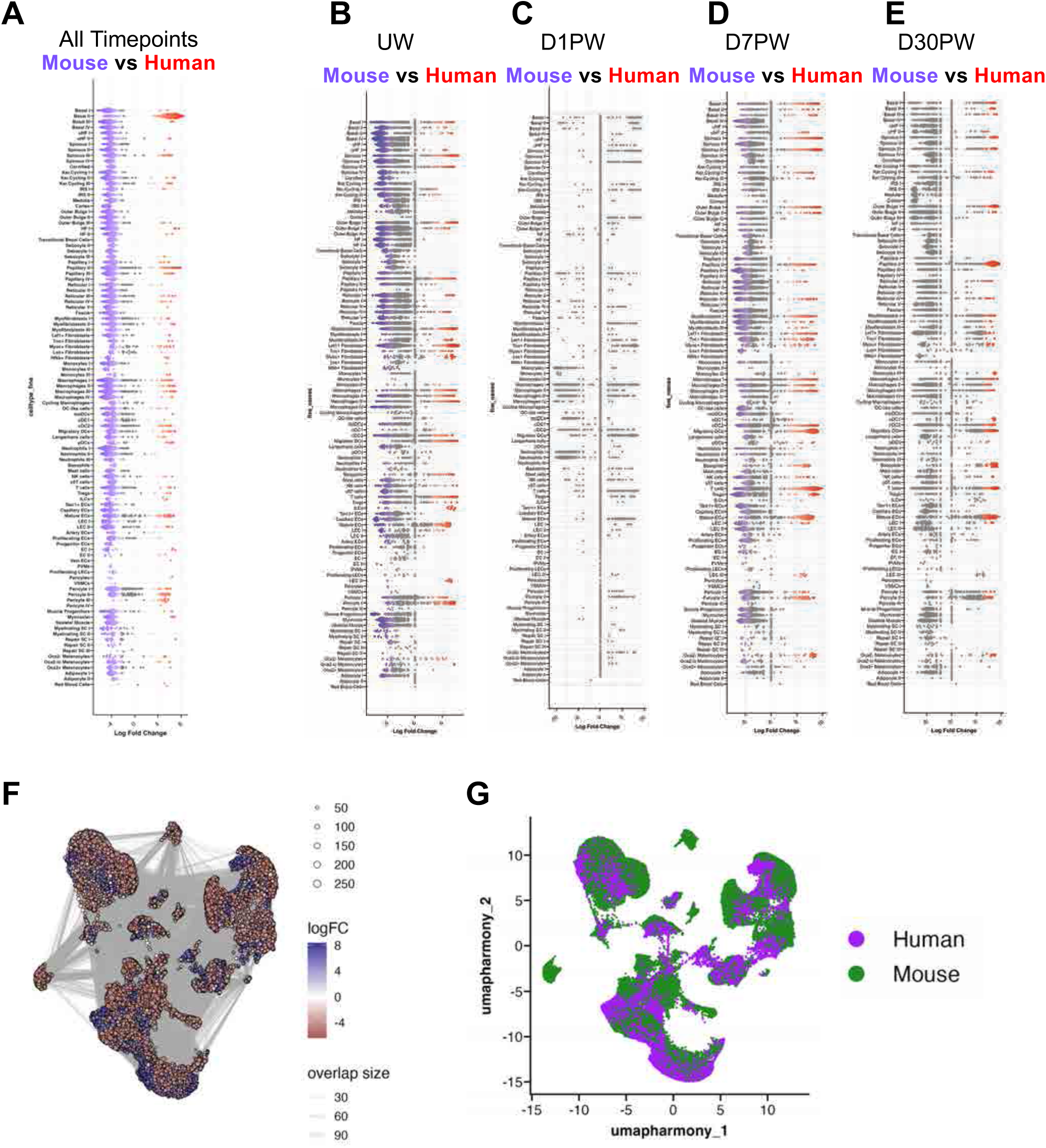
Differential abundance of cell types between species. **(A–D)** Bee swarm plots showing Milo-derived differential abundance for 107 fine cell-type subclusters in the cross-species integrated atlas. Mouse-enriched subclusters are shown in blue, and human-enriched subclusters in red (FDR < 0.01 for matched timepoints). Panels represent: **A)** unwounded (UW), **B)** D1PW, **C)** D7PW, and **D)** D30PW. (**E)** Global comparison of Milo-derived differential abundance values across all subclusters. (**F)** UMAP projection of the cross-species atlas with Milo differential abundance scores overlaid, highlighting species-enriched cellular states (red = human-enriched, blue = mouse-enriched).( **(G)** UMAP projection of harmony-integrated cross-species atlas with mouse and human samples colored (human = purple, mouse = green). Down sampled to 80,000 cells

**Supplemental Figure 29.**
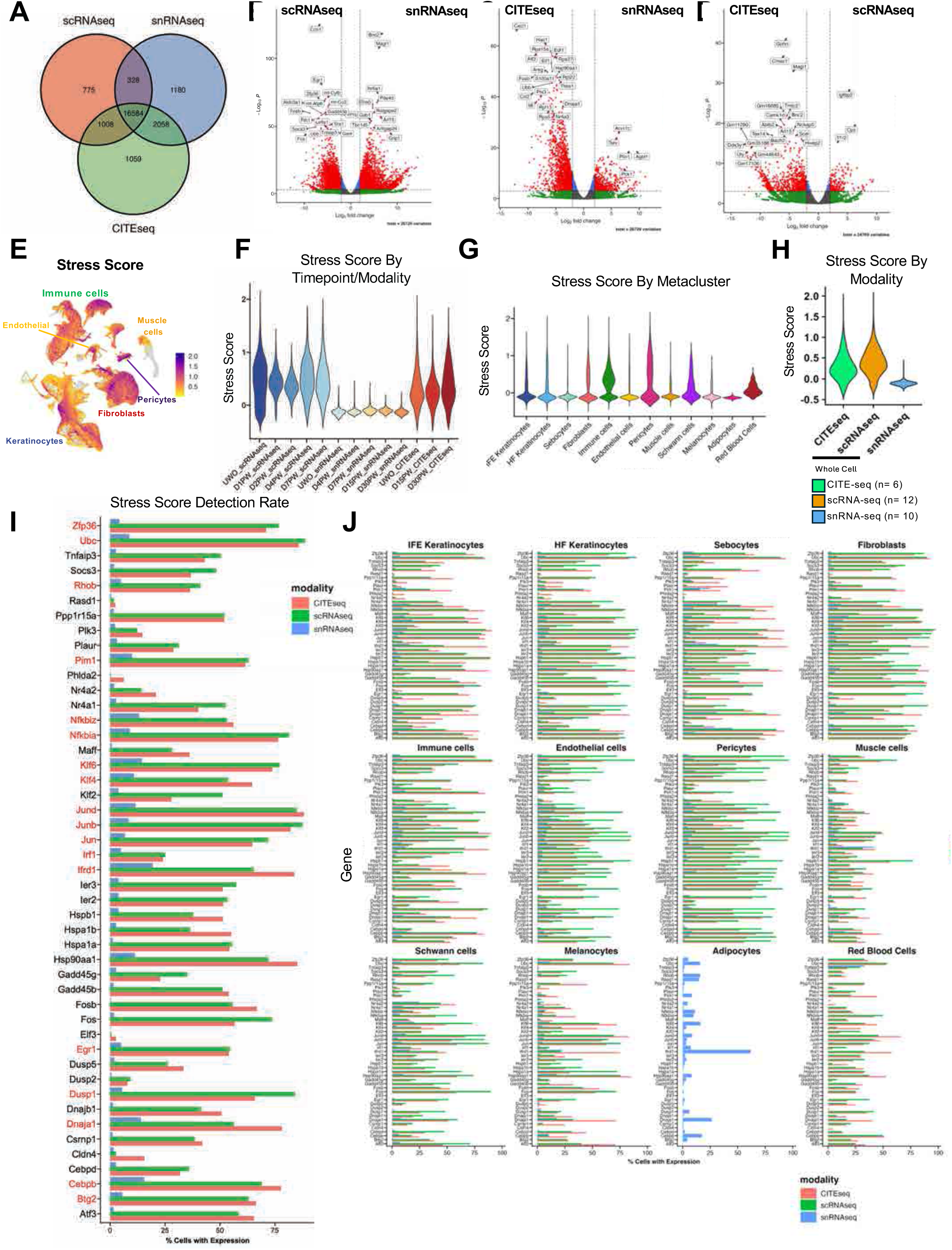
Multimodal gene detection and stress signature benchmarking in the OWHA atlas. **(A)** Venn diagram showing the overlap of genes detected across sequencing modalities included in the OWHA atlas: scRNA-seq (n = 12 samples), snRNA-seq (n = 10), and CITE-seq (n = 6). **(B–D)** Volcano plots showing differential gene expression comparisons between sequencing modalities in unwounded skin. Comparisons include (**B**) scRNA-seq versus snRNA-seq, (**C**) CITE-seq versus snRNA-seq, and (**D**) CITE-seq versus scRNA-seq. Differentially expressed genes were defined by |log₂FC| > 2 and adjusted *p* < 0.01. **(E)** Feature plot visualizing per-cell Stress Score across the RPCA-integrated OWHA UMAP embedding. **(F–H)** Violin plots showing Stress Score distributions across the OWHA atlas, stratified by (**F**) timepoint, (**G**) metacluster, and (**H**) sequencing modality. (**I**) Bar plot showing the percent detection rate for each gene included in the stress gene set, colored by sequencing modality (CITE-seq, red; scRNA-seq, green; snRNA-seq, blue). Genes highlighted in red indicate those retained for regulon-based stress scoring, defined by a detection rate ≥10% in snRNA-seq. (**J**) Bar plots showing percent detection rates of stress-associated genes across sequencing modalities, faceted by metacluster.

**Supplemental Figure 30:**
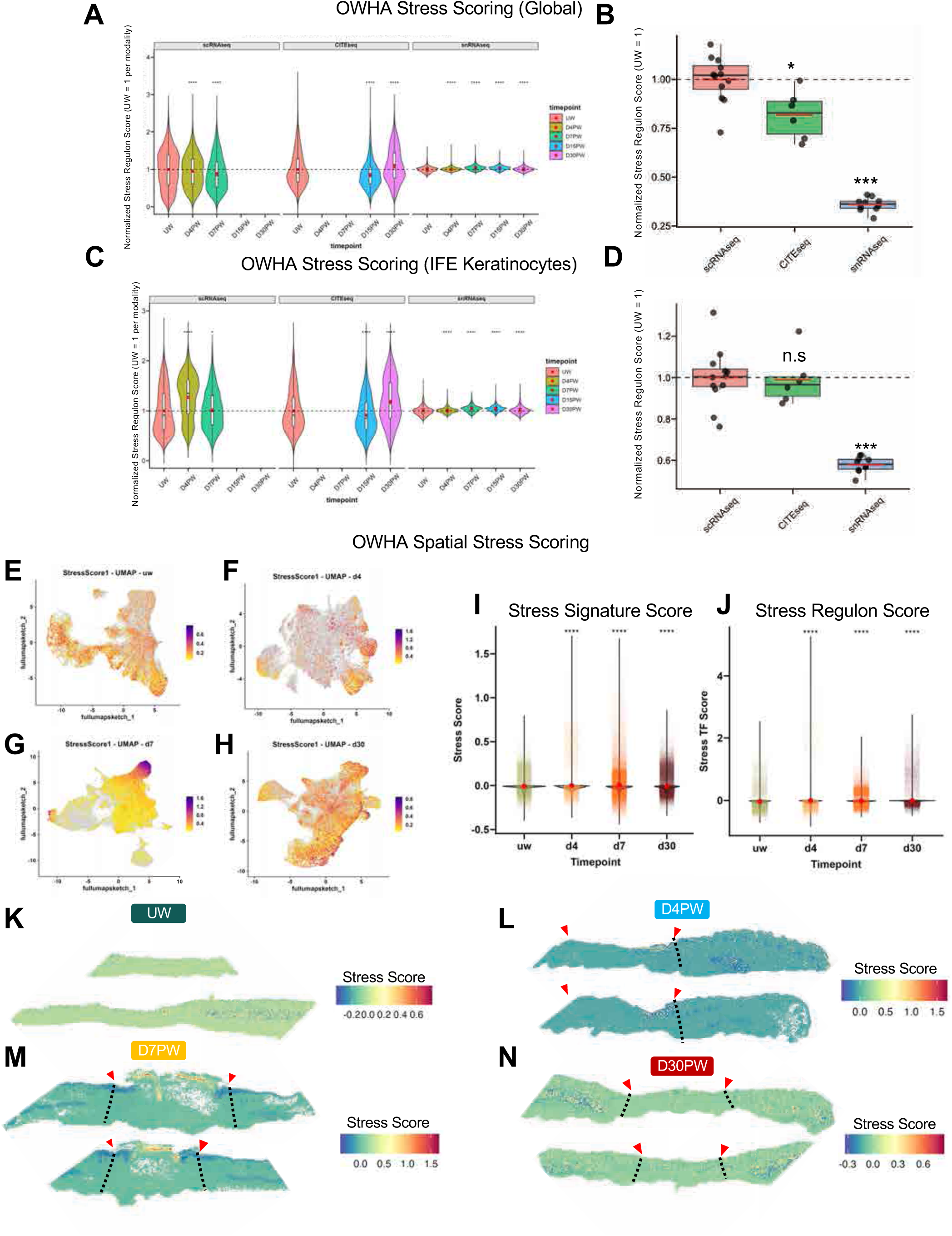
Single-nucleus and spatial transcriptomic datasets exhibit reduced stress score signatures relative to whole-cell sequencing approaches. **(A)** Violin plots showing distributions of the transcription factor–based regulon Stress Score across wound-healing timepoints in all OWHA cells, faceted by sequencing modality. Stress Scores were normalized within each modality to the unwounded (UW) condition (UW = 1). Statistical significance was assessed using a Wilcoxon rank-sum test compared to UW for each modality. **(B)** Boxplots showing mean Stress Scores per sequencing run, colored by modality (scRNA-seq, red; CITE-seq, green; snRNA-seq, blue). Values were normalized to the mean scRNA-seq Stress Score (scRNA-seq = 1). Each point represents an independent sequencing run (n = 28). **(C)** Violin plots showing transcription factor–based regulon Stress Score distributions specifically within interfollicular epidermis (IFE) keratinocytes across timepoints, faceted by modality. Stress Scores were normalized to the UW condition within each modality (UW = 1). Statistical significance was assessed using a Wilcoxon rank-sum test compared to UW for each modality. **(D)** Boxplots showing mean Stress Scores for IFE keratinocytes across sequencing runs, colored by modality (scRNA-seq, red; CITE-seq, red; snRNA-seq, blue). Values were normalized to the mean scRNA-seq Stress Score (scRNA-seq = 1). Each point represents an independent sequencing run (n = 28). **(E–H)** Feature expression UMAP plots showing Stress Score values calculated on Visium HD spatial transcriptomic data for mouse skin at (**E**) UW, (**F**) D4PW, (**G**) D7PW, and (**H**) D30PW. (**I-J**) Violin plots quantifying (**I**) Stress Signature Score using all stress-associated genes and (**J**) transcription factor–based Stress Regulon Score in Visium HD data. Red points show the mean value for each timepoint across n = 2 technical replicates. Significance was assessed using a Wilcoxon rank-sum test compared to UW (p < 0.05 = *, p < 0.01 = **, p < 0.001 = ***). **(K-N)** Spatial feature plots showing Stress Signature Score values across the wound axis for each timepoint in the OWHA spatial atlas: **(K)** UW, **(L)** D4PW, **(M)** D7PW, and **(N)** D30PW.

**Supplemental Figure 31:**
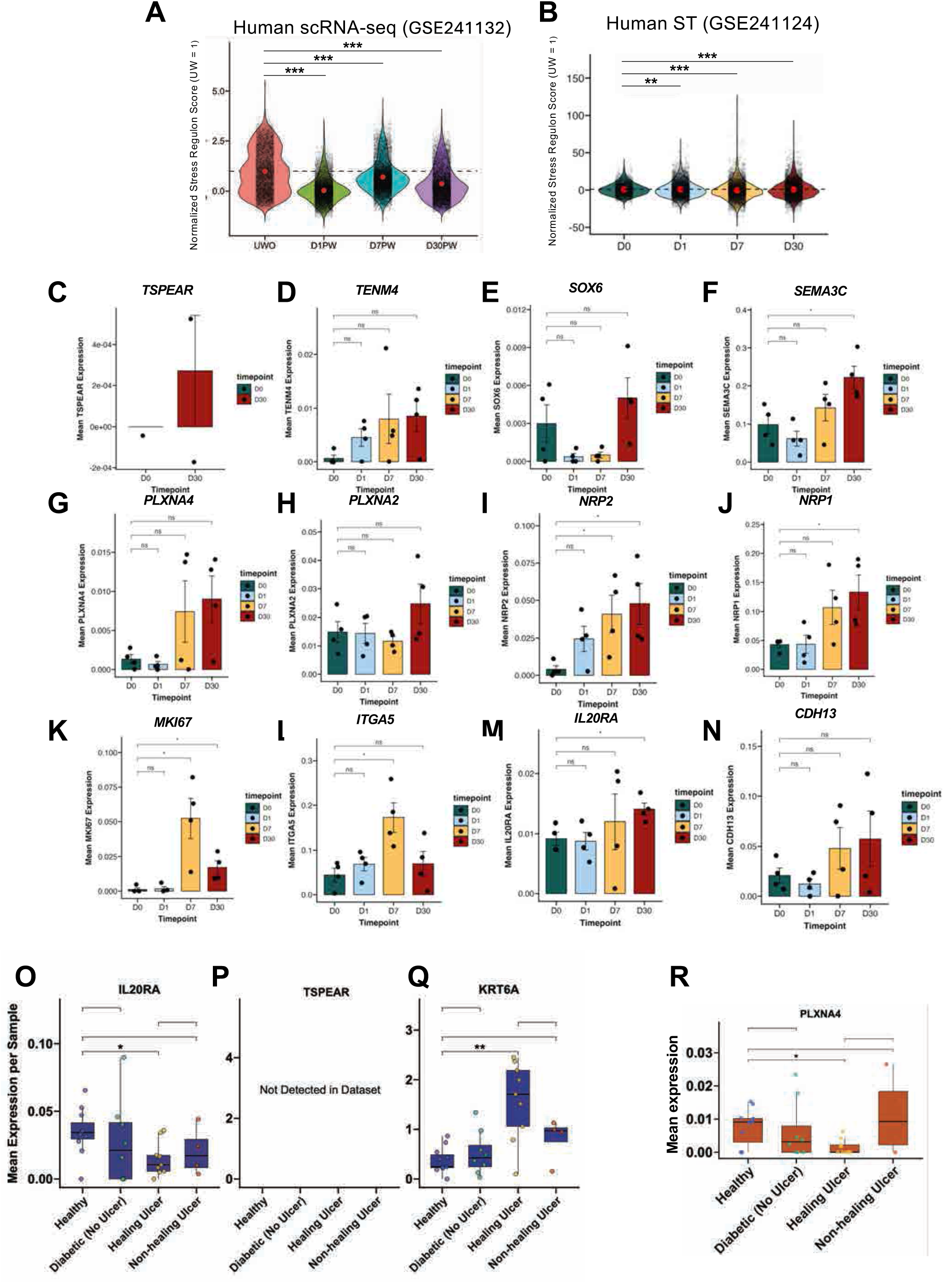
Basal IV signature in human spatial and diabetic data. **(A-B)** Violin plots showing Stress Regulon Score distributions for (**A**) human scRNA-seq data and (**B**) human spatial data (Liu et al., 2024; GSE241132), normalized to the UW mean. Statistical significance was assessed using a Wilcoxon rank-sum test compared to UW. (* p < 0.05; ** p < 0.01, p < 0.001 = ***). **(C-N)** Box plots depicting mean wound edge expression of genes associated with Basal IV cells in human spatial data (Liu et al., 2024; GSE241132) across healing timepoints (D0PW, D1PW, D7PW, D30PW). Each dot represents the mean signal per sequencing run. Statistical significance was assessed using a Wilcoxon rank-sum test relative to D0/UW (* p < 0.05; ** p < 0.01, p < 0.001 = ***). *n* = 4. **(O-Q)** Box plots showing expression of Basal IV markers in keratinocytes from human diabetic wound healing data (GSE265972). Each point represents an individual sequencing run. Statistical significance was assessed using a Wilcoxon rank-sum test compared to healthy wounds. (**R**) Box plot showing expression of SEMA3C-receptor PLXNA4 in endothelial cells from human diabetic wound healing data (GSE265972). Each point represents an individual sequencing run. Statistical significance was assessed using a Wilcoxon rank-sum test compared to healthy wounds.

## Notes

### Competing Interest Statement

The authors have declared no competing interest.

